# Integration of Magnetic Tweezers and Traction Force Microscopy for the Exploration of Matrix Rheology and Keratinocyte Mechanobiology: Model Force- and Displacement-Controlled Experiments

**DOI:** 10.1101/2020.12.03.410704

**Authors:** Waddah I. Moghram, Pratibha Singh, Christian A. VandeLune, Edward A. Sander, John C. Selby

## Abstract

In this work we demonstrate the integration of magnetic tweezers (MT) with substrate deformation tracking microscopy (DTM) and traction force microscopy (TFM) for the investigation of extracellular matrix rheology and human epidermal keratinocyte mechanobiology in the context of human blistering skin diseases. Two model bead-on-gel experiments are described in which an MT device is used to apply a prescribed force or displacement waveform to a fibronectin-coated superparamagnetic bead attached to a type I collagen gel containing a layer of covalently attached red-fluorescent microspheres. Serial fast time-lapse DIC and epifluorescence image acquisitions are used to capture displacements of the bead and microspheres, respectively, in response to the applied force or displacement. Due to the large number of acquired images and the dynamic behavior of substrate microspheres observed during the experiment, new quantitative methods are developed for the tracking and filtering of microsphere displacement data, the selection of L2 regularization parameters used for TFM analysis, and the identification of time intervals within the overall image set that can be approximated as being subject to elastostatic conditions. Two major proof-of-concept applications are described in which integrated MT-DTM/TFM experiments are used to (i) estimate the elastic properties of a fibrillar type I collagen gel substrate and (ii) demonstrate how a force applied to a focal adhesion contact on the apical surface of a living keratinocyte is directly transmitted to basal cell-matrix anchoring junctions as observed by substrate deformations and incremental traction stresses that develop within the collagen subjacent to the cell.

## I. INTRODUCTION

The human epidermis-the outermost layer of skin-comprises an organized assembly of epithelial cells known as keratinocytes^1^. Within the epidermis, basal keratinocytes adhere to one another via specialized cell-cell anchoring junctions known as desmosomes and adherens junctions^1,2^, whereas keratinocytes within the innermost layer of the epidermis are also anchored to the underlying connective tissue of the dermis by specialized cell-extracellular matrix (cell-matrix) anchoring junctions known as hemidesmosomes and focal adhesion contacts^3^. Within individual keratinocytes, keratin intermediate filaments associate with desmosomes and hemidesmosomes, while actin microfilaments converge on focal adhesions and adherens junctions^3,4^. Collectively, these anchoring junctions and their associated cytoskeletal proteins form a transcellular filamentous network superstructure that is critical to the mechanical barrier function of skin. In humans, several different types of congenital and acquired skin fragility disorders exist in which specific components of this cytoarchitectural complex are rendered dysfunctional^2,5–7^. Affected areas of skin form blisters, erosions, and ulcerations, thereby impairing the skin’s ability to regulate temperature and water balance, while permitting invasion by external pathogens^2,8^.

Despite advances in our understanding of the pathophysiology of known skin fragility disorders, from a biophysical standpoint, the mechanisms by which anchoring junctions and transcellular cytoskeletal protein networks endow keratinocytes with such an innate mechanical resilience are not completely understood^2,9^. Fortunately, over the past several decades, numerous experimental methodologies have been developed for exploration of cell mechanobiology and matrix rheology, including magnetic tweezers (MT)^10–14^, deformation tracking microscopy (DTM)^15,16^, and cell traction force microscopy (TFM)^17–20^. With respect to multicellular systems, MT devices have been used to explore local viscosity within embryonic tissues^21^, whereas DTM/TFM has been used to directly measure traction stresses associated with collective cell migration^22^ and to infer cell-cell force transmission within living cell clusters^15,23,24^. To date, only limited work has been done to integrate MT and DTM/TFM experiments for biological applications^25,26^. Towards this end, we set out to demonstrate how integration of MT and DTM/TFM can be used to explore the mechanobiology of the human epidermis in the context of skin fragility disorders. As the design, operation, and calibration of our MT device is comprehensively reviewed in a companion paper^27^, herein we describe two model approaches to *directly* investigate force transmission in reconstituted collagen matrices and living human epidermal keratinocyte tissue constructs *in vitro*: force-control and displacement-control mode MT-DTM/TFM experiments. Explicit details are provided on the relevant physical approximations and computational methods used to analyze the large image-based data sets generated from each of these two types of MT-DTM/TFM experiment. Two major proof-of-concept applications are described in which an integrated MT-DTM/TFM experiment is used (i) to estimate the elastic properties of a fibrillar type I collagen gel substrate and (ii) to demonstrate force transmission through a living human epidermal keratinocyte localized to the periphery of a reconstituted multicellular sheet.

## II. CONCEPT OF AN INTEGRATED MT-DTM/TFM EXPERIMENT

In order to optimally explore mechanical force transmission within a multicellular keratinocyte sheet *in vitro*, we postulate that the experimenter should have the ability to apply a force that is (i) *specific* to a cell-cell or cell-matrix anchoring junction of interest and that is (ii) of *sufficient magnitude* to generate cellular and/or underlying substrate deformations that can be physically observed and characterized within the framework of applied mechanics^28^. Based on these idealized constraints, we chose to integrate the techniques of MT and DTM/TFM to demonstrate how a force applied to a living keratinocyte is transmitted to its underlying substrate. A cartoon schematic of our methodology is detailed in **Fig. 1(a)**. Depending on culture conditions, individual keratinocytes or an interconnected multicellular sheet of keratinocytes are cultured on a soft deformable substrate with fluorescent microspheres embedded on its surface. The substrate must mimic physiologically relevant matrix of the tissue environment *in vivo* while also being compatible with keratinocyte culture *in vitro*, i.e., the surface mechanochemistry must be conducive to normal cell attachment, spreading, and viability. As an additional constraint, the substrate must be mechanically compliant, such that forces and/or displacements applied the MT device result in observable substrate deformations.

**FIG. 1.**
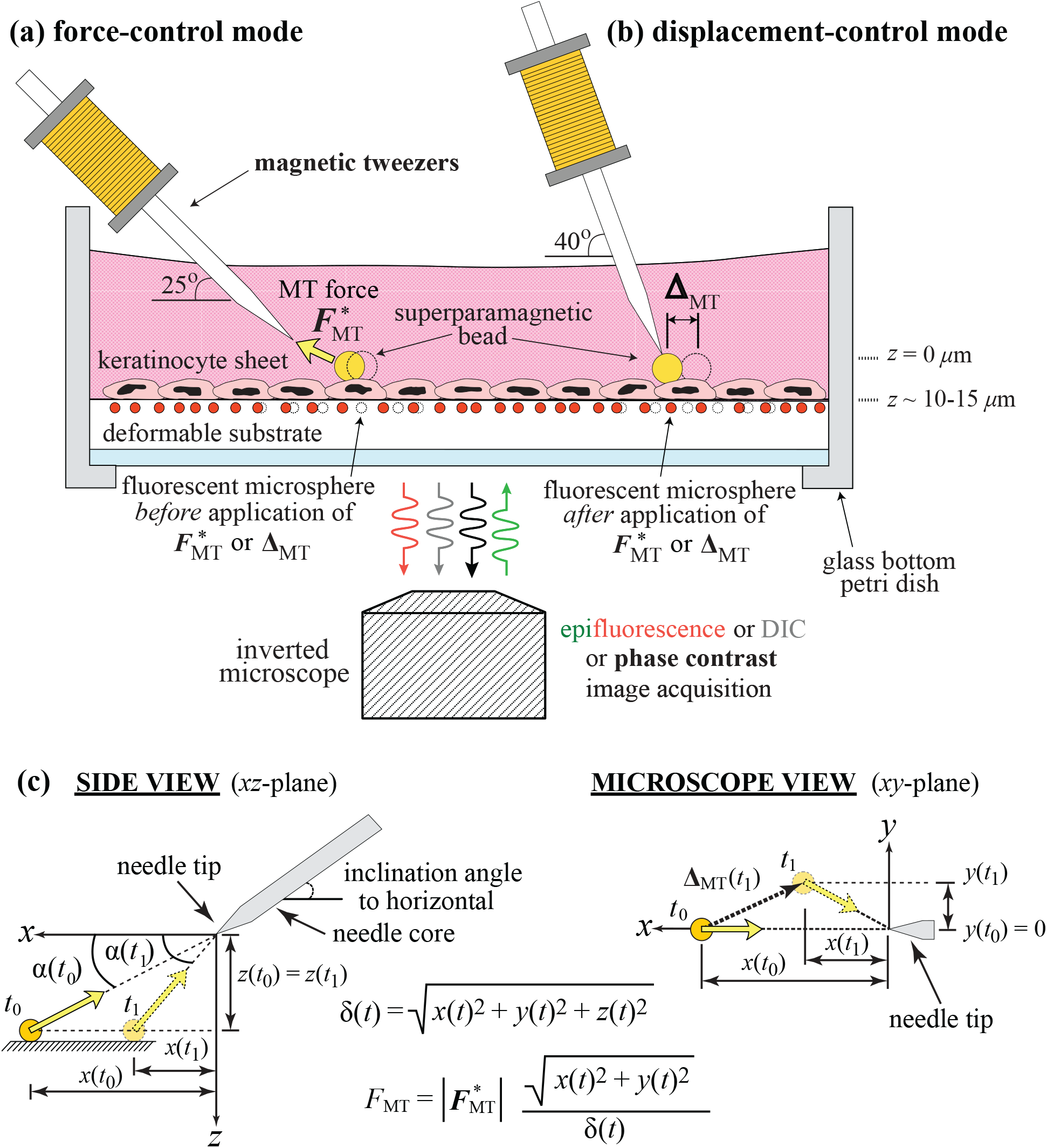
Schematic of an integrated MT-DTM/TFM experiment. Schematic conceptually demonstrates two prototypical modes of operation in an integrated MT-DTM/TFM mechanobiological experiment. **(a)** In force-control (FC) mode, the MT device exerts a prescribed magnetic actuation force or force waveform, 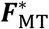, on a superparamagnetic bead attached to a keratinocyte. **(b)** In displacement-control (DC) mode, the MT device is actuated to magnetically clamp a bead to the needle tip. Subsequent translation of the MT needle can then be used to impose a prescribed displacement or displacement waveform on the bead, **Δ_MT_**. Depending on the presence or absence of cell-cell and cell-matrix anchoring junctions, application of 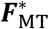 or **Δ_MT_** can generate a deformation in the underlying culture substrate, observed by tracking displacements of individual fluorescent microspheres embedded in its surface. For both modes of operation, experiments require the acquisition of two sequential yet coupled DIC and epifluorescence imaging sets during which an identical 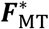 or **Δ_MT_** actuation sequence is applied to the superparamagnetic bead. **(c)** An orthogonal (*x, y, z*)-coordinate system with its origin fixed to the needle tip of the MT device is used to spatially locate the position of the superparamagnetic bead at any instance of time, *t*, a vector defined here as ***δ***(*t*). Bead position is illustrated at two different time points, *t*_0_ and *t*_1_. Yellow block arrows are used to indicate the *xy*- and *xz*-projections of the magnetic actuation force vector, 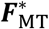. Here, *z*(*t*) is assumed to be constant for the duration of the experiment as motion of the bead is largely restricted to the *xy*-plane. Note in **(c)** how the angle between ***δ***(*t*) and the *xy*-plane changes as a function of bead position, as shown by α(*t*), the projection of this angle in the *xz*-plane. Although the bead and the *y*-axis of the needle are grossly aligned at *t*_0_, small *y*-displacements of the bead (exaggerated in the above schematic) are observed during actuation of the MT device, as imaged within the *xy*-focal plane of the microscope. The bead-tip Euclidean separation distance, δ(*t*), and geometry of the setup are deterministic of the vectoral component of 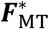 present within the *xy*-plane of the substrate, a variable defined as ***F***_MT_, with a scalar magnitude denoted as *F*_MT_.

In an integrated MT and DTM/TFM experiment, a superparamagnetic bead is covalently functionalized with a specific ligand of interest (e.g., fibronectin, collagen, E-cadherin, desmocollin, desmoglein, etc.). The bead is then allowed to bind to its native ligand(s) or receptor(s) on the surface of an individual cell forming a cell-cell or cell-ECM anchoring junction mimic, i.e., a mechanobiochemical coupling between the superparamagnetic bead and the cell that can be used to experimentally model one or more critical attributes of a true biological anchoring junction. In a force-control mode (FC) experiment (see **Fig. 1(a)**), a defined force or force waveform, 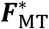, is selectively applied to the anchoring junction mimic using the MT device during two serial imaging experiments. Note that throughout this work, bolded variables are used to represent vector-valued functions, while non-bolded variables are used to represent scalar valued functions. In the first experiment, 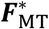 is applied while differential interference contrast (DIC) imaging is used to monitor displacements of the superparamagnetic bead. In the second experiment, 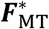 is applied again, but this time epifluorescence imaging is used to capture displacements of the fluorescent microspheres embedded in the surface of culture substrate. Tracked and quantified, microsphere displacements can be used to approximate a continuum representation of the substrate displacement field, **u** (i.e., DTM). If substrate displacements are both small and confined to within the microscopic field of view, it is possible to calculate the *incremental* stress traction vector field, ***T***, that develops on the surface of the substrate in response to 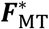 (i.e., TFM). In a displacement-control mode (DC) experiment (see **Fig. 1(b)**), the tip of the MT device is positioned in close proximity to the attached superparamagnetic bead of interest, and then actuated to magnetically clamp the bead to the needle tip. The MT needle is then translated to impose a displacement or displacement waveform, **Δ_MT_**, on the bead in the horizontal plane of the substrate (see **Fig. 1(b)**). As for FC mode, two serial imaging experiments are also performed in DC mode, the first in which DIC imaging is used to capture displacements of the superparamagnetic bead, and a second in which epifluorescence imaging is used to track displacements of substrate microspheres. The same prescribed **Δ_MT_** is applied for both imaging studies. Again, tracking of fluorescent microsphere displacements in the underlying substrate (***u***) allows for calculation of ***T*** that develops in response to **Δ_MT_**. For both modes of operation, it is important to recognize that experiments require the acquisition of *sequential* yet coupled DIC and epifluorescence imaging sets during which an identical 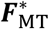 or **Δ_MT_** actuation sequence is being applied to the superparamagnetic bead. The DIC image set records motion of the superparamagnetic bead and the cell, whereas the epifluorescence imaging set captures the kinematics of the microspheres embedded in the culture substrate. Additional phase contrast or DIC imaging can also be used to provide wide-field views of the reconstituted cell sheet.

## III. METHODS

### A. Instrumentation

The instrumentation and control schemes employed to operate our MT device are extensively detailed in a companion paper^27^. With respect to DTM/TFM, in brief, our setup consisted of a Nikon Eclipse Ti-E inverted microscope (Nikon, Melville, NY) controlled by an HP Z72 Workstation running Nikon Elements AR 4.51 software on a non-deterministic 64-bit Windows^®^ operating system. A PCO.EDGE 4.2 sCMOS camera (PCO-Tech, Wilmington, DE) was used to acquire images at a standard frame rate of 40 frames/second (fps) and a resolution of 2048×2044 pixels. Frame rates of up to 100 fps for smaller defined regions of interest (ROIs) were used for some experiments. The microscope was equipped with standard DIC, phase contrast, and epifluorescence imaging modalities. Nikon CFI Plan Fluor DL 10X (0.3 NA), Plan Apo Lambda 10X (0.3 NA) or CFI Plan Apo VC 20X (0.75 NA) air objective lenses coupled with an optional 1.5X magnifier were used in the collection of all experimental image sets. Epifluorescence imaging used a Texas Red bandpass filter cube set (560/40 nm excitation filter, 595 nm dichroic mirror, 630/60 nm emission filter). DIC or phase contrast imaging was used for visualization of keratinocyte sheets. The MT device was controlled via a Dell Optiplex 9010 desktop computer that used a LabVIEW™ Real-Time 15.0.1 deterministic operating system (National Instruments, Austin, TX), equipped with a 16-bit NI PCIe-6341 DAQ board. The DAQ board was used to acquire analog input measurements (i.e., magnetic flux density, solenoid current, and transistor-transistor logic (TTL) pulses marking exposures of the sCMOS camera) while generating analog output voltages to control actuation of the MT device in real-time. The MT device was mounted on an MHW-3 three-axis water hydraulic manual micromanipulator (Narishige, Amityville, NY) that was fitted with an aluminum adaptor arm fabricated in-house to extend the reach of the needle tip over the microscope stage.

### B. Collagen substrates

Although polyacrylamides and silicones are commonly used in TFM experiments, we found that these materials proved incompatible with keratinocyte culture when formulated with the mechanical compliances needed for our integrated MT-DTM/TFM experiments. As such, we elected to use type I collagen gel substrates, prepared as follows. The working surfaces of 50 mm-diameter petri dishes with 35 mm-diameter No. 1.5 coverslip glass bottoms (MatTek, Ashland, MA) were treated with a BD-20AC corona discharge treater (Electro-Technic Products, Chicago, IL) to render the surface hydrophilic due to an increase in hydroxyl moieties on the glass surface^29,30^. Next, the glass was silanized by covering with 97% 3-aminopropyltriethoxysilane (3-APTMS; Sigma Aldrich, St. Louis, MO) for 5 minutes, followed by multiple rinses with deionized (DI) water. The surface was then covered in a freshly prepared 5% glutaraldehyde solution (Sigma Aldrich) for 30 minutes, followed by multiple rinses of DI water^15,31^. Rat tail type I collagen solutions (1.0 to 3.0 mg/mL) were prepared at 4°C from a mixture of phenol-red 10X Dulbecco’s phosphate buffered saline (DPBS; Gibco, Dublin, Ireland), DI water, and rat tail type I collagen (Corning Inc, Corning, NY), titrated to a pH of 7.4 using 1 N NaOH according to the manufacturer’s protocol. The collagen solution was then dispensed into an ~18 mm inner diameter Sylgard 184 polydimethylsiloxane (PDMS; Dow Corning, Midland, MI) circular mold fixed to the glass surface. PDMS molds were fabricated in-house with thicknesses ranging from 250 μm to 750 μm and outer diameters of ~24 mm. Specimens were subsequently incubated at 37°C/5% CO_2_ for 60 minutes to allow for collagen fibrillogenesis and hence gel formation, after which the PDMS molds were removed. A second circular PDMS mold with an inner diameter of ~24 mm and an outer diameter of ~28 mm was then placed around the fully formed collagen gel to create a temporary liquid reservoir. The surface of the collagen gel was then covered with a sonicated solution of 0.5 μm-diameter FluoSpheres™ carboxylate-modified polystyrene red fluorescent microspheres (ThermoFisher Scientific, Grand Island, NY) and incubated at room temperature for 2 minutes. Loosely physiosorbed microspheres were rinsed from the surface via multiple washes with 1X DPBS. To covalently attach the fluorescent microspheres to the collagen fibril network, the PDMS reservoir was filled with a 2 mM solution of 1-ethyl-3-(3-dimethylaminopropyl)carbodiimide (EDAC; ThermoFisher Scientific) in 50 mM 2-(N-morpholino)ethanesulfonic acid hydrate (MES; Sigma Aldrich) at a pH of 6.0 for 15 minutes at room temperature. The EDAC/MES solution was then aspirated and the specimen was rinsed multiple times with 1X DPBS. The final surface density of red fluorescent microspheres was approximately 0.06 spheres/μm^2^. Following the covalent attachment step, PDMS rings were removed, and the entire dish was filled with 1X DPBS to prevent gel drying. Specimens were sealed with parafilm and stored at 4°C for up to 3 weeks prior to use.

### C. Keratinocyte culture and multicellular sheet reconstitution

Primary adult human keratinocytes (HEKs; ATCC, Manassas, VA) were cultured in T75 flasks at 37°C/5% CO2 in keratinocyte serum-free medium (KSFM; Gibco) containing L-glutamine, and supplemented with 50 mg/mL of bovine pituitary extract, 2.5 ng/mL of recombinant human epidermal growth factor, 100 U/mL penicillin, 100 mg/mL of streptomycin, 0.205 mg/mL of sodium deoxycholate and 0.25 μg/mL of amphotericin. Fully supplemented KSFM contains low concentrations of calcium (<0.2 mM), and is otherwise referred to here as low [Ca^2+^] KSFM. Fully supplemented KSFM with a calcium concentration of ~1.2 mM was also prepared via addition of sterile CaCl_2_, referred to here as high [Ca^2+^] KSFM. Sheets were reconstituted by harvesting cells >48 hours following passage 4 or 5. To seed a sheet of keratinocytes on a type I collagen gel substrate, 1X DPBS covering the collagen gel for storage was aspirated and the petri dish containing the gel was sterilized by a 10-to 15-minute exposure to 254 nm ultraviolet light in the biosafety cabinet. Next, 18 μL droplets containing 200 cells/μL in low [Ca^2+^] KSFM were dispensed onto the central area of the gels and sterile DI water was pipetted into the petri dish around the edges of the gel to prevent dehydration while keratinocytes attached and spread over the collagen surface. The specimens were incubated at 37°C/5% CO_2_ for 3 hours, after which the water was aspirated, and the dish was flooded with either low [Ca^2+^] or high [Ca^2+^] KSFM and incubated for an additional 24 to 72 hours. This procedure consistently generated multicellular, mixed monolayer and partially stratified keratinocyte sheets on the central area of the collagen gels, ~3 to 6 mm in diameter. Under low [Ca^2+^] conditions, individual keratinocytes within a sheet adhere to the underlying collagen only via focal adhesion contacts^2,32^. However, under high [Ca^2+^] conditions, keratinocytes within a multicellular sheet become mechanically interconnected via cell-cell adherens junctions and desmosomes^2,15,33–36^.

### D. Functionalization and attachment of superparamagnetic beads

Superparamagnetic beads used in both collagen and keratinocyte MT-DTM/TFM experiments were prepared according to the bead manufacturer’s protocol by incubating 4.5 μm-diameter Dynabeads^™^ M-450 Tosylactivated^®^ superparamagnetic beads with human plasma fibronectin purified protein (Catalog No. FC010; MilliporeSigma, St. Louis, MO) reconstituted at a concentration of 1 mg/mL in a 0.1 M sodium phosphate buffer containing 2 mM of EDAC. Incubation was allowed to proceed with gentle tilting at 4°C for a total of 5 minutes. For experiments probing the rheology of a collagen gel, a small droplet of fibronectin-coated superparamagnetic beads diluted to 15 beads/μL in 1X DPBS was dispersed on the surface of a collagen gel and incubated at 37°C/5% CO_2_ for 10 minutes to allow for bead settling. Following this initial incubation, the dish was gently filled with additional DPBS after which the specimen was incubated overnight at 37°C/5% CO_2_ to promote more robust bead attachment to the substrate via native hydrophobic adhesive interactions between collagen and fibronectin^37^. For experiments involving keratinocytes, fibronectin-coated superparamagnetic beads were again diluted in 1X DPBS. After aspirating the overlying KSFM, the bead suspension was dispensed as a small droplet on the surface of a keratinocyte sheet and incubated at 37°C/5% CO_2_ for 10 minutes. After this brief incubation, the loosely adherent beads and excess solution was removed, and the entire petri dish was then flooded with supplemented KSFM and incubated overnight at 37°C/5% CO_2_. In this manner, keratinocytes form a focal adhesion contact (FAC) with the magnetic bead via adhesive interactions between integrins within the FAC and fibronectin coupled to the surface of the bead^38^.

### E. Displacement tracking

Integrated MT-DTM/TFM experiments based on DIC image sets were analyzed by first selecting a global ROI appropriate for data analysis. Within this global ROI, the spatial position of the centroid of the superparamagnetic bead was quantified within each frame using a custom MATLAB (Mathworks, Inc., Natick, MA) code based on *imfindcircles* and/or the geometric transformation function *imregtform* (restricted to rigid body translation without rotation) applied to a small fixed local ROI (within the global ROI) that captured the bead’s position throughout the overall image sequence. Use of the local ROI for centroid tracking allowed for markedly more efficient computation times. For FC and DC experiments (see Sec. IV.A), positions of the needle tip of the MT device were not tracked. In addition to superparamagnetic bead positions, the code tracked displacements of small corner ROIs confined to the four corners of the global ROI (each ~10% of the global ROI) using the MATLAB function *imregtform*. A substrate drift correction to the superparamagnetic bead position in each frame was calculated by subtracting the average rigid body translation of the four corner ROIs from the raw measured bead position.

Epifluorescence image sets of the microspheres embedded within the surface of our collagen substrates were analyzed to estimate ***u***(*x*\*, *y*\*, *t*), a two-dimensional continuum representation of the substrate displacement field on a time-invariant equally spaced rectangular (*x*\*, *y*\*)-positional grid for each image frame or imaging time, *t*. A concise conceptual outline of the steps required for reduction of microsphere displacement data can be found elsewhere^39^. In this work, we employed a modified version of an open-source MATLAB TFM code originally developed to investigate force modulation in subresolution cellular adhesions^40^. Identical global ROIs were used for both DIC and epifluorescence image sets. For epifluorescence analysis, individual fluorescent microspheres were first detected using a two-dimensional Gaussian fit of the intensity distribution for a point source^41^. After microsphere identification, incremental microsphere positions were tracked using pixel correlation with subpixel fitting (PCSF)^40^, sometimes followed by an additional tracking step based on a modified version of subpixel correlation by image interpolation (SCII)^42^. The template size was set to be larger than the average separation distance between microspheres, and the maximum displacement was set at least equal to the maximum displacement of the superparamagnetic bead observed in the corresponding DIC image set.

After tracking of raw microsphere positions, a multistep temporospatial displacement correction algorithm was applied to the data set. First, local displacement outliers within the global ROI at each time (or imaging frame), *t*, were automatically detected from the data set using a universal normalized median residual test algorithm adapted from particle tracking velocimetry and particle image velocimetry methods^43,44^. Corrected displacements for these spatial outliers were interpolated from the measured displacements of neighboring microspheres. Second, unique to our DTM/TFM analysis, spurious high frequency frame-to-frame noise in the individually tracked microsphere displacements was attenuated by passing the entire data set through a low-pass digital filter with its design specifically tailored to the actuation waveform of the experiment. For FC mode of operation (see Sec. IV.A.1), individual microsphere displacements were temporally filtered using an equiripple linear-phase FIR digital filter with 41 taps, a passband of 2 Hz, and a stopband of 10 Hz. For DC experiments (see Sec. IV.A.2), displacements were temporally filtered using an equiripple linear-phase FIR digital filter with 41 taps, a passband of 10 Hz, and a stopband of 15 Hz. Temporal filtering introduced a time shift in the filtered microsphere position/displacement data, and thus the final 0.5 s (or ~20 imaging frames) of each FC or DC displacement experiment were omitted from analysis. As the last step in our algorithm, spatially corrected and temporally filtered microsphere displacements present within small square ROIs localized to the four corners of the global ROI at *t* = 0 (each ~10% of the global ROI) were used to compute a frame-by-frame substrate drift correction. In each frame, the average displacement of microspheres contained in these corner ROIs was subtracted from the displacement measured for every microsphere contained within the global ROI. As with displacement tracking for DIC image sets, rotational drift corrections were not performed.

As a final step in data reduction, the optimal (*x*\*, *y*\*)-grid was determined as a function of *x*\*- and *y*\*-domains of microsphere positions within the global ROI, the total number of tracked microspheres, and the average spacing of microspheres. Then, for each imaging time, *t*, we computed ***u***(*x*\*, *y*\*, *t*) by applying the MATLAB function *griddata* (set to cubic interpolation) to the spatial-outlier corrected, temporally filtered, and drift-corrected microsphere displacement data set. Due to the random (*x*\*, *y*\*)-distribution of the microsphere data, cubic interpolation failed to estimate displacements for some (*x*\*, *y*\*)-grid positions. To fill these “holes”, biharmonic spline interpolation was used to calculate missing values using the interpolated displacements and their corresponding (*x*\*, *y*\*)-grid positions as inputs to the *griddata* function, *not* the original microsphere displacement data set^45^. In this manner, C^2^ continuity was preserved throughout the interpolated displacement field ***u***(*x*\*, *y*\*, *t*).

### F. Traction force microscopy

Excellent reviews of TFM and the various analytical and numerical approaches used to calculate stress traction vector fields, ***T***(*x*\*, *y*\*, *t*), from observed substrate displacement fields, ***u***(*x*\*, *y*\*, *t*), can be found elsewhere^39,46,47^. In brief, we computed the inverse solution for ***T***(*x*\*, *y*\*, *t*) using data sets where ***u***(*x*\*, *y*\*, *t*) were small, and where ***u***(*x*\*, *y*\*, *t*) along the boundary of the global ROI were tending towards zero for all *t* of the epifluorescence image set. The collagen substrate was modeled as a spatially homogeneous, isotropic, linear elastic half-space, with a Young’s elastic modulus, *E*, and Poisson’s ratio, *v*, which to a first-order approximation holds true, given the relatively small deformations observed in the majority of our experiments. To calculate ***T***(*x*\*, *y*\*, *t*), interpolated (or gridded) displacement fields were filtered in the spatial domain with a Wiener filter (3×3 pixel)^20^ and then padded by a margin of zeros^39,48^. A Hann window was applied to the filtered and padded data set to minimize spectral leakage^39,48^. The gridded, padded, and windowed displacement data set was then transformed into Fourier space^39^. Solutions for ***T***(*x*\*, *y*\*, *t*) were computed for each frame of a collective fast time-lapse image sequence using an open-source MATLAB code employing the methodology of Fourier Transform Traction Cytometry (FTTC) with an L2 regularization scheme^20,40^. FTTC was chosen over other methodologies due to its computational efficiency given the large number of imaging frames to be analyzed per experiment. Algorithmic determination of the L2 regularization parameter(s) employed a Bayesian methodology^49^, as detailed in the **Supplementary Material**(see SI.A.2 and SI.B.2).

Once ***T***(*x*\*, *y*\*, *t*) were determined, the total traction force present at the surface of the substrate at time, *t*, denoted here as ***F***(*t*), and its magnitude, *F*(*t*), were calculated by numerical integration of ***T***(*x*\*, *y*\*, *t*):

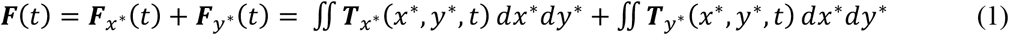

where the subscripts *x*\* and *y*\*represent the vectoral components of ***T***(*x*\*, *y*\*, *t*) and ***F***(*t*) in the *x*\*- and *y*\*-directions, respectively, and integration is carried out over a domain that includes the entire (*x*\*, *y*\*) positional grid configured for the global ROI (see Sec. SI.A.2 in the **Supplementary Material**). In addition to ***T***(*x*\*, *y*\*, *t*), ***F***(*t*), and *F*(*t*), we also computed the strain energy density field, *ρ*(*x*\*, *y*\*, *t*), as

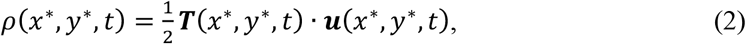

and the total strain energy associated with deformation of the substrate, *U*(*t*), as

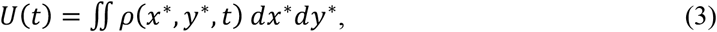

where, again, the limits of integration were set to span the domain of the complete (*x*\*, *y*\*) positional grid configured for the global ROI. Note that in typical TFM analysis of a single cell adherent on a deformable substrate, ***F***(*t*) is identically zero because the adherent cell is mechanically *static* at the time scale under consideration. Thus, the stress traction vector field that the cell produces in the underlying substrate does *not* generate a net force. In contrast, for the MT-DTM/TFM experiments demonstrated in this work, ***F***(*t*) are non-zero because the *incremental* traction stresses imposed on the collagen substrate result from a non-zero force generated by actuation of the MT device. In the idealized case where a superparamagnetic bead directly attached to the surface of a linear elastic substrate is subject to a static magnetic actuation force, 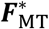 (see **Fig. 1(c)**), the above analysis should yield ***F***(*t*) = ***F***_MT_(*t*), where ***F***_MT_(*t*) represents the vectoral component of 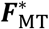 present within the *xy*-plane of the substrate.

As for the mechanical properties of the type I collagen substrates used in our TFM calculations, we assumed ν = 0.4 for all collagen substrates^50^. As a first pass in our TFM analyses, we assumed concentration-dependent shear moduli, *G*(*c*), of 5.0 Pa, 55.4 Pa, and 341.8 Pa for 1 mg/mL, 3 mg/mL, and 7 mg/mL collagen gels, respectively, as measured by the collagen manufacturer (Corning) using parallel plate rheometry^51^. Here, the variable, *c*, is used to denote collagen concentration. In this work, collagen fibrillogenesis was carried out at 37°C, as was done in the Corning study^51^. For other collagen concentrations within the range of 1 mg/mL to 7 mg/mL, *G*(*c*) were estimated from the equation, *G*(*c*) ≈ (5.0 Pa)*c*:.9,^52^ and all concentration-dependent elastic moduli, *E*(*c*), were calculated from the implied linear elastic relationship, *E*(*c*) = 2*G*(*c*)(1 + ν) (assuming ν = 0.4). As an alternative to assuming mechanical properties for our TFM analyses, we also directly estimated *E*(*c*) for each collagen substrate that was tested using our standardized force-control mode of operation, as detailed in Sec. IV.A.1.

### G. Ethics approval

Ethics approval was not required.

## IV. MT-DTM/TFM APPLICATIONS

### A. Rheology of a collagen gel

#### 1. Force-control mode experiment

For a model FC MT-DTM/TFM experiment, we dispersed a dilute suspension of 4.5 μm-diameter, fibronectin-coated superparamagnetic beads directly onto a ~680 μm thick, 1.0 mg/mL type I collagen substrate embedded with a surface layer of covalently attached red fluorescent microspheres. Preparation of the collagen substrate, functionalization of the superparamagnetic beads, and attachment of superparamagnetic beads to the substrate were carried out as detailed in Secs. III.B and III.D. Attached beads were spatially separated by at least one field of view at 30X magnification. With the microscope configured for 30X magnification, an FC experiment was performed using sequential DIC and epifluorescence fast time-lapse image capture steps, during which identical 3-cycle 3 s/7 s ON/OFF magnetic actuation waveform sequences with maximum calibrated ON magnetic flux density setpoints of 175 Gs were applied by the MT device. As opposed to perfectly square ON/OFF waveforms, magnetic actuation in our setup included short stepwise ramps in calibrated magnetic flux density values between ON and OFF states. Specifically, the MT device was stepped up (or down) to ON (or OFF) states passing through intermediate calibrated magnetic flux density values of 25 Gs, 50 Gs, 75 Gs, 100 Gs, 125 Gs, and 150 Gs, maintaining each of these intermediate flux density setpoints for ~70 ms before proceeding to the next setpoint (up or down). As a consequence of this magnetic actuation scheme, there exists a ~500 ms transient interval in magnetic actuation force, 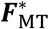, that separates ON and OFF (0 nN) states of a given actuation cycle. Use of stepwise ON/OFF ramps during magnetic actuation was intentional, because it facilitated more reliable tracking of the kinematics of the fluorescent microspheres embedded in our collagen substrates. Explicit details of the actuation of our MT device are described in the **Supplementary Material** (see Sec. SI.A.1 and **Fig. S1**).

The initial (*x, y, z*)-spatial location of the superparamagnetic bead relative to the needle tip of the MT device was *x*(0) = 20.0 μm, *y*(0) = 0 μm, and *z*(0) = 13.0 μm (δ(*t*_0_) ~ 23.9 μm; see **Fig. 1(c)**). Prior to data collection, a trial 3-cycle magnetic actuation waveform was applied to the superparamagnetic bead to precondition the mechanical response of the collagen substrate. Following data collection, all images were analyzed according to the methods presented in the **Supplementary Material** (see Sec. SI.A.2). Of note, TTL pulses marking exposures of the sCMOS camera were used to temporally correlate DIC and epifluorescence image data to the magnetic flux density recorded during actuation of the MT device. Complete real-time videos of the coupled DIC and epifluorescence imaging data analyzed for this experiment can be found in **Vid 1** and **Vid 2**, respectively. In all DIC images, the superparamagnetic bead remains clearly in focus throughout the magnetic actuation waveform sequence, as does the shadow of the needle tip of the MT. Although the shadow of the needle tip is apparent in the epifluorescence image set, the superparamagnetic bead is not easily identified.

In comparing the kinematics of the superparamagnetic bead to the mechanical deformation of the collagen substrate in response to the same magnetic actuation waveform sequence, consider the data shown in **Fig. 2**. Here, in what is referred to here as a bead-on-gel experiment, we define the time point, *t*_OFF_, for each cycle within the overall magnetic actuation waveform sequence as the last imaging frame captured prior to initiation of the stepwise *decrease* of 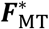 towards a null actuation force 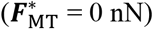. Similarly, we define the time point, *t*_ON_, for each cycle as the last imaging frame captured prior to initiation of the stepwise *increase* of 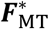 towards the maximum actuation force. **Figs. 2(a)** and **2(b)** depict the DIC and epifluorescence images captured near *t*_OFF_ for the first magnetic actuation cycle, respectively. Now, consider plots of 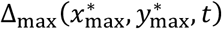 and Δ_MT_(*t*) as shown in **Fig. 2(e)**. Here, 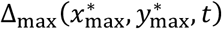 is used interchangeably with Δ_max_(*t*), representative of the magnitude of the displacement vector of the microsphere embedded on the collagen substrate subject to the largest observed displacement during the experiment, or 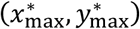. Δ_MT_(*t*) represents the magnitude of the displacement vector of the superparamagnetic bead. As can be seen in **Fig. 2(e)**, at *t* = *t*_OFF_ for cycles #1, #2, and #3, the maximum substrate displacement was ~2.0 μm less than that recorded for the superparamagnetic bead, a quantity on the order of the average radius of the bead (~2.25 μm). In testing numerous superparamagnetic beads attached to collagen substrates in FC MT-DTM/TFM experiments, this finding was repeatable. For superparamagnetic beads that were allowed to establish conformal contact with the substrate for >12 hours, values of Δ_MT_(*t*_OFF_)-Δ_max_(*t*_OFF_) for a given cycle were less than ~2.25 μm for experiments in which Δ_MT_(*t*_OFF_) < 2.25 μm. These values plateaued at ~2.25 μm for experiments in which Δ_MT_(*t*_OFF_) ≥ 2.25 μm. For experiments in which superparamagnetic beads were only allowed to establish conformal contact with the substrate for a short period of time (~1 hour), greater differences were observed between Δ_MT_(*t*_OFF_) and Δ_max_(*t*_OFF_) for Δ_MT_(*t*_OFF_) < 2.25 μm. Moreover, these beads frequently detached from the substrate for Δ_MT_(*t*_OFF_) ≥ 2.25 μm. Based on these physical observations, we speculate that the superparamagnetic bead might both rotate and translate in response to the magnetic actuation exerted by our MT device, dependent on the state of conformational contact between the bead and the substrate. Although previous authors have attempted to model substrate deformations that occur when an attached superparamagnetic bead is subject to an applied torque^53^, contact models characterizing the response (to force) observed here represent a subject of future investigation.

**FIG. 2.**
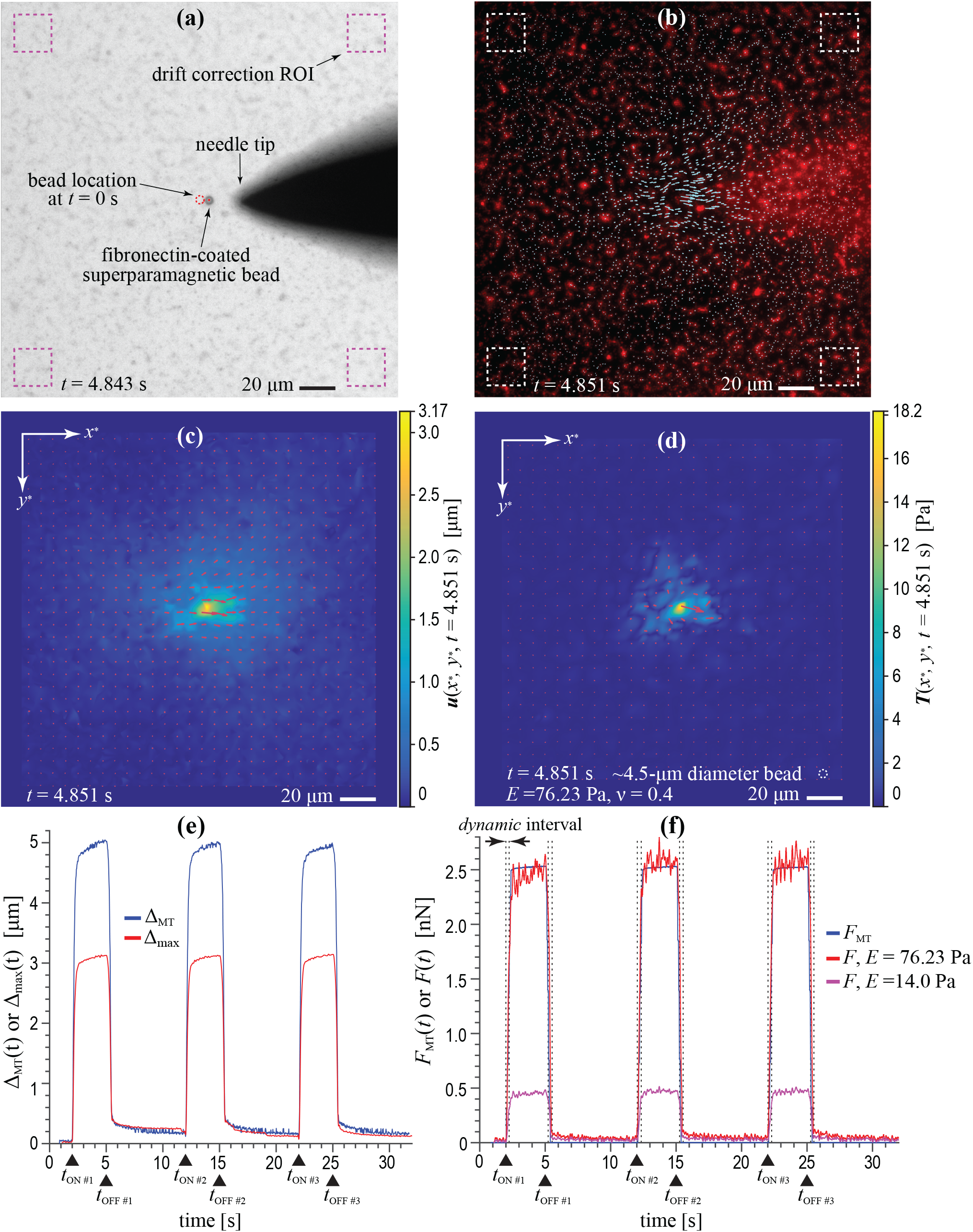
FC mode bead-on-gel experiment. Six-panel figure highlights data from a model FC mode bead-on-gel experiment involving a fibronectin-coated 4.5 μm-diameter superparamagnetic bead attached to the surface of a ~680 μm-thick, 1.0 mg/mL type I collagen substrate embedded with a surface layer of covalently attached red fluorescent microspheres. **(a)** DIC image showing the superparamagnetic bead near its peak displacement during the first ON cycle of the applied magnetic actuation waveform sequence. **(b)** Texas Red epifluorescence image with overlaid quivers that represent the displacement vectors of individual microspheres present within the surface of the collagen substrate at this same time point in the magnetic actuation waveform. Dashed boxes in **(a)** and **(b)** denote the ROIs used for drift correction. The displacement field, ***u***(*x*\*, *y*\*), and stress traction vector field, ***T***(*x*\*, *y*\*), computed for the microsphere displacement field displayed in **(b)**, are shown in **(c)** and **(d)**, respectively, where for aid of visualization, the white dashed circle in **(d)** represents a ~4.5 μm-diameter circle. A plot of the displacement magnitude of the microsphere subject to the largest overall displacement during actuation, Δ_max_(*x*_OPQ_, *y*_OPQ_, *t*), versus the magnitude of the superparamagnetic bead displacement, Δ_MT_(*t*), is shown in **(e)**. A plot of the magnitude of the component of the magnetic actuation force vector in the *xy*-plane of the collagen substrate, *F*_MT_(*t*), versus the magnitude of the integrated total traction force vector, *F*(*t*), is shown in **(f)**, where *F*(*t*) has been computed assuming υ = 0.4 and two different elastic moduli, *E* = 14.0 Pa and *E* = 76.23 Pa. Black triangles in **(e)** and **(f)** denote *t*_ON_ and *t*_OFF_ for each actuation cycle. Vertical dashed lines in **(f)** indicate *dynamic* time intervals in which the assumption of elastostatic conditions is likely invalid.

Real-time videos depicting continuum representations of the substrate displacement field, ***u***(*x*\*, *y*\*, *t*), the stress traction vector field, ***T***(*x*\*, *y*\*, *t*), and the strain energy density field, *ρ*(*x*\*, *y*\*, *t*), for this experiment, assuming *E* = 76.23 Pa and *u* = 0.4, can be found in **Vid 3**, **Vid 4,** and **Vid 5**, respectively. **Figs. 2(c)** and **2(d)** depict ***u***(*x*\*, *y*\*, *t* = 4.851 s) and ***T***(*x*\*, *y*\*, *t* = 4.851 s), respectively, where *t* = 4.851 s is very near *t*_OFF_ for the first actuation cycle. Considering the inherent assumption of linear elasticity, overall, the solution computed for ***T***(*x*\*, *y*\*, *t* = 4.851 s) appears to be physically reasonable for the observed ***u***(*x*\*, *y*\*, *t* = 4.851 s), i.e., a quasi-symmetrical distribution of traction stresses around a localized circular traction stress that has a diameter approximately equal to that of the superparamagnetic bead (~4.5 μm). Integrating ***T***(*x*\*, *y*\*, *t*) over the (*x*\*, *y*\*) positional grid for all time, *t*, provides a mean for computing the total traction force, ***F***(*t*) and its scalar magnitude, *F*(*t*). In **Fig. 2(f)**, we compare *F*(*t*) to the magnitude of the component of the magnetic actuation force exerted by the MT device in the *xy-*plane (and *x*\**y*\*-plane) of the collagen substrate, or *F*_MT_(*t*), as calculated from DIC imaging data (see SI.A.2 of the **Supplementary Material**). With regards to force, note that *F*_MT_(*t*) approximates a square waveform with small ~500 ms transients between ON and OFF states (see SI.A.1 of the **Supplementary Material**). This finding is somewhat counterintuitive, based on the fact that for a constant nominal calibrated magnetic flux density setpoint, the magnetic actuation force, 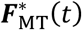, *increases* as the superparamagnetic bead more closely approaches the needle tip^27^. However, based on the geometry of our MT setup, the vectoral component of 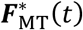 in the *xy-*plane of the collagen substrate remains roughly constant due to increases in the relative angle between ***δ***(*t*) and the *xy-*plane as the bead approaches the needle tip (see **Fig. 1(c)**).

Neglecting viscous and inertial effects, if the collagen substrate behaved purely as an ideal linear elastic material, both Δ_max_ and Δ_MT_ should have exhibited displacement responses that qualitatively mirrored the approximate square waveform of *F*_MT_(*t*). However, as can be seen in **Fig. 2(e)**, both Δ_max_(*t*) and Δ_MT_(*t*) exhibit evidence of a viscoelastic response to the applied magnetic actuation force^54^. Allowing for a viscoelastic response, but assuming that mechanical equilibrium is achieved at *t*_OFF_ for each of the three actuation cycles, one should find that *F*_MT_(*t*_OFF_) = *F*(*t*_OFF_) upon integrating ***T***(*x*\*, *y*\*, *t*_OFF_) over the (*x*\*, *y*\*) domain of the global ROI. However, as can be seen in **Fig. 2(f)**, if one uses *E* = 14.0 Pa as suggested by the rheologic data published for a 1.0 mg/mL type I collagen gel^51^, *F*(*t*_OFF_) ≠ *F*_MT_(*t*_OFF_). As such, we determined the *apparent E* for our collagen substrates by combining information from the DIC and epifluorescence image sets collected during identical time intervals of the magnetic actuation waveform sequence. Specifically, for the final ~500 ms of the ON portion of the second magnetic actuation cycle, we iterated the assumed value of *E* within our ***T***(*x*\*, *y*\*, *t*) calculations until the computed values for *F*(*t*) during this time interval best fit, in a least squares sense, the magnetic actuation force, *F*_MT_(*t*), using the derivative-free, unconstrained minimum search function *fminsearch* in MATLAB. Note that each iteration of the search required re-computation of the L2 regularization parameters for the data set, given its dependence on *E*. As can be seen in **Fig. 2(f)**, an apparent *E* of ~76.23 Pa provides a much closer match between *F*_MT_ and *F* at *t* = *t*_OFF_ for each of the three actuation cycles. However, with increasing modulus, the mean noise floor in *F* during OFF segments of magnetic actuation also increases, from ~0.03 nN for *E* = 14.0 Pa to ~0.2 nN for *E* = 76.23 Pa (also see **Fig. S2** in the **Supplementary Material**). One possible explanation for the observed difference between the reported value of *E* and our best-fit apparent *E* is that the elastic modulus of our collagen substrates was altered by the presence of the fluorescent microspheres, the EDAC-based chemical treatment used to confer covalent attachment of the microspheres to the collagen fibrils, or the brief UVC exposure used to sterilize the substrate. Additionally, subtle variations in reconstitution conditions, e.g., differences in temperature and pH, are also known to affect the fibrillar structure and hence the mechanical properties of type I collagen gels^55^. Alternatively, differences between the reported value of *E* and our best-fit apparent *E* might be attributed to the length scales at which the respective mechanical measurements were performed, i.e., the local elasticity of a collagen fibrillar network as measured by our FC MT-DTM/TFM experiment versus the bulk parallel plate rheometry measurements used by Corning^51,56^.

Using an energetics approach, we also estimated an upper bound for the apparent *E* from this same combined DIC and epifluorescence image set. Using DIC images, we calculated the work done by the MT device in moving the superparamagnetic bead during the ON segment of a specified cycle of the overall magnetic actuation waveform sequence, a variable denoted here as *W*_MT_(*t*), and calculated from tracked superparamagnetic bead positions as:

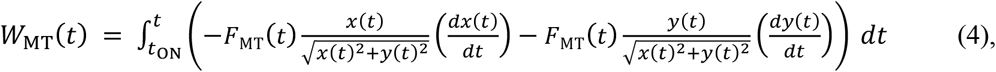

where 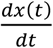 and 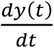 represent differential displacements of the magnetic bead in the *x*- and *y*-directions, respectively, and *t* marks any time point such that *t*_ON_ < *t* ≤ *t*_OFF_. Again, *F*_MT_(*t*) represents the magnitude of the vectoral component of the magnetic actuation force in the *xy*-plane (or *x*\**y*\*-plane) of the collagen substrate. To estimate the upper bound of the apparent *E*, we first computed *W*_MT_(*t* = *t*_OFF_) for the *second* cycle of the 3-cycle magnetic actuation waveform. We then iterated the assumed value of *E* within our ***T***(*x*\*, *y*\*, *t*) calculations for the corresponding epifluorescence image at time, *t* = *t*_OFF_, of the second actuation cycle to find the minimum difference between *U*(*t*_OFF_) and *W*_MT_(*t*_OFF_). Note that again, this required computation of *λ*2_ON_ using the entire data set for each iterated value of *E*. Optimization solutions utilized *fminsearch* in MATLAB. By neglecting energy lost to the system as viscous dissipation within the collagen gel and instead equating *W*_MT_(*t*_OFF_) to *U*(*t*_OFF_) at *t* = *t*_OFF_ (for cycle #2), we approximated an upper bound to the apparent *E* of the collagen substrate. **Fig. 3** demonstrates temporal profiles of *W*_MT_(*t*) and *U*(*t*) for this experiment. *U*(*t*) was calculated assuming both *E* = 14.0 Pa and an apparent *E* = 149.8 Pa, the latter representing the upper bound of *E* estimated for this collagen specimen using an energetics approach.

**FIG. 3.**
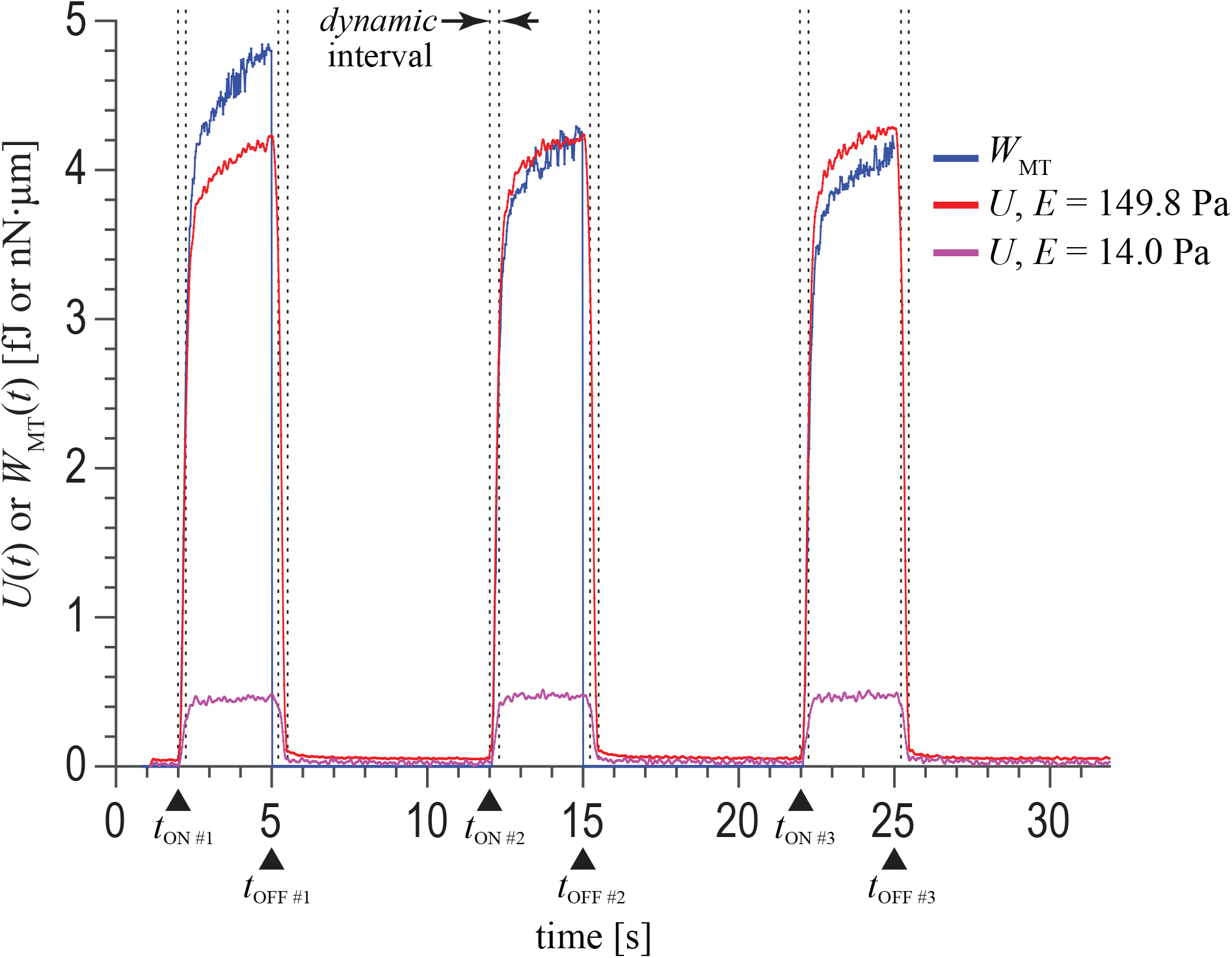
Plot showing *W*_MT_(*t*), the work done by the MT device in moving the superparamagnetic bead during each ON segment for each of the three magnetic actuation cycles, versus *U*(*t*), the total strain energy stored in the collagen substrate during the course of the magnetic actuation waveform sequence. An upper bound to the apparent elastic modulus of the collagen substrate is found by iterating on the assumed value of *E* in computing ***T***(*x*\*, *y*\*, *t*) and *U*(*t*), and then equating *U*(*t* = *t*_OFF_) to *W*_MT_(*t* = *t*_OFF_) for actuation cycle #2. Black triangles denote *t*_ON_ and *t*_OFF_ for each actuation cycle. Vertical dashed lines indicate *dynamic* time intervals in which the assumption of elastostatic conditions is likely invalid.

In this work, we observed that for a given ***u***(*x*\*, *y*\*, *t*), selection of the L2 regularization parameter, λ2, can have a profound effect on the solution for ***T***(*x*\*, *y*\*, *t*), in terms of the distribution and magnitude of the resultant stress traction vector field. In our first attempt to objectively select λ2 for the data set, we used a Bayesian approach^49^ to objectively compute a raw, time-specific value of λ2 for each imaging frame on which to base the solution of ***T***(*x*\*, *y*\*, *t*). However, when these independent stress traction vector fields were assembled into a collective time sequence, subtle non-physical temporal discontinuities in the frame-to-frame evolution of ***T***(*x*\*, *y*\*, *t*) could be observed. Consequently, we devised an algorithmic approach for computing ***T***(*x*\*, *y*\*, *t*) that utilized two global L2 regularization parameters, λ2_ON_ and λ2_OFF_ (as described in Sec. SI.A.2 of the **Supplementary Material**), that provided more physically realistic traction stress fields while minimizing traction noise, especially during the OFF segments of the actuation sequence. A graphical demonstration of the computation of λ2_ON_ and λ2_OFF_ corresponding to the FC mode experiment detailed in **Fig. 2** can be found in **Fig. S3** of the **Supplementary Material**.

Lastly, note that all of the foregoing TFM analysis applied to our epifluorescence image data is based on the solution of a classic *elastostatics* problem, i.e., one that assumes infinitesimal displacements of a spatially homogeneous, isotropic, linear elastic half-space in static equilibrium with one or more applied surface traction forces^57^. However, in an integrated FC MT-DTM/TFM experiment, the magnetic actuation waveform sequence generated by the MT device produces *dynamic* motion of the collagen substrate. Although we compute ***T***(*x*\*, *y*\*, *t*), ***F***(*t*), *ρ*(*x*\*, *y*\*, *t*), and *U*(*t*) corresponding to the measured ***u***(*x*\*, *y*\*, *t*) for every analyzed imaging frame, solutions for ***T***, ***F***, *ρ*, and *U* are only approximately valid at time points during the magnetic actuation sequence when inertial contributions to the global force balance in this boundary-value problem are negligible. Clearly, at time points where inertial contributions are *not* negligible, the stress traction vector field solution, as described in Secs. III.F, would be invalid.

As an *ad hoc* experimental means for identifying time intervals within our overall data set for which the assumptions underlying the elastostatics model are valid, we performed an additional FC experiment, referred to here as a null force control experiment. With the microscope configured for 30X magnification, the identical superparamagnetic bead of interest used in the preceding experiment was positioned such that *x*(0) = 20.0 μm, *y*(0) = 0 μm, and *z*(0) = 13.0 μm with respect to the needle tip. The bead was then actuated with a 3-cycle 3 s/7 s ON/OFF waveform sequence with maximum nominal ON magnetic flux density setpoints of 0 Gs, exactly as done for the previous prototype FC experiment. In other words, the MT device was actuated with a 3-cycle waveform sequence such that ***F***_MT_(*t*) = 0 nN for all time points captured in the sequential DIC and epifluorescence fast time-lapse capture image sets. With a null actuation force, the collagen substrate remains static throughout data acquisition and analysis, in contrast to the dynamic substrate motions observed during the prototype FC experiment (see **Fig. 2**). Real-time videos of the coupled DIC and epifluorescence imaging data analyzed for this null force control experiment, as well as videos computed for ***u***(*x*\*, *y*\*, *t*), ****T****(*x*\*, *y*\*, *t*), and *ρ*(*x*\*, *y*\*, *t*), respectively, can be found in **Vid 6**, **Vid 7**, **Vid 8**, **Vid 9**, and **Vid 10**, respectively, the latter two assuming *E* = 76.23 Pa and *u* = 0.4. **Fig. S2** in the **Supplementary Material** summarizes data from the null force control experiment in a manner that is analogous to data presented for the prototype FC experiment shown in **Fig. 2**. The global ROIs used to analyze both the null FC experiment and the prototype FC experiment were identical, i.e., the central 1024 × 1022 pixels of each image.

In this null force control experiment, 3266 microspheres were tracked over 1236 imaging frames. As done for the model bead-on-gel FC experiment, measured microsphere displacements for the null force control experiment were corrected for spatial outliers, temporally filtered, and drift-corrected (see Sec. III.E). For each imaging frame in the data set, we computed the *x*\*- and *y*\*-vectoral components of the displacement tracked for each microsphere embedded within the collagen substrate, denoted here as 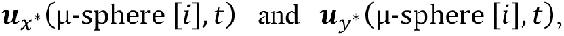 respectively, where μ-sphere [*i*] denotes a unique microsphere present in the collagen substrate identified by the index, *i*. Starting with the third imaging frame, we also computed the *x*\*- and *y*\*-components of microsphere accelerations, denoted here as 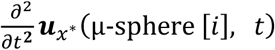 and 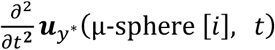, respectively, using a backward difference methodology. For each analyzed frame, we then computed averages for the *x*\*- and *y*\*-components of displacements and accelerations observed over all 3266 microspheres (*i* = 1 thru 3266), in addition to statistical upper and lower bounds of this data defined by ±6 standard deviations (±6SD). Disregarding data from the first and second imaging frames where accelerations were undefined, we computed the global means of the average displacements, accelerations, and 6SD upper⁄lower bounds observed over the remaining 1234 imaging frames (see **Fig. S4** of the **Supplementary Material**). Because none of the 3266 microspheres experienced a *true* displacement or acceleration during the null force control experiment, the mean 6SD upper⁄lower bounds on these displacement and acceleration observations represent estimates of the physical detection limits (and thus measurement uncertainty) intrinsic to our experimental setup and microsphere tracking algorithms.

Assuming identical microscope settings, global ROI selection, and microsphere tracking procedures as described for the model bead-on-gel experiment, we assert that any microsphere displacement or acceleration observed during an FC experiment as described here cannot be distinguished as being *true*, versus *artifactual*, if it falls within the mean 6SD upper⁄lower bounds of the *x*\*- and *y*\*-displacements and accelerations computed for the null force control experiment. Accordingly, consider the prototype FC experiment originally detailed in **Fig. 2**, and more specifically, the microsphere embedded within the collagen substrate that was subject to the overall largest displacement during magnetic actuation, or 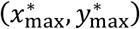. The *x*\*- and *y*\*-components of the displacement and acceleration of this microsphere, denoted here as 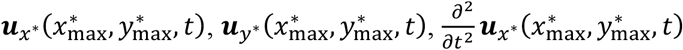 and 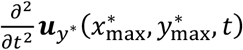, respectively, are plotted in **Fig. 4**. The mean 6SD upper⁄lower bounds for *x*\*- and *y*\*-displacements and accelerations as derived from the null force control experiment are included with each plot (see **Fig. S4**). In **Fig. 4**, one finds that in terms of *x*\*- and *y*\*-accelerations, the prototype FC experiment is potentially dynamic within small 275 to 325 ms (11 to 13 imaging frames) or 150 to 225 ms (6 to 9 imaging frames) time intervals, respectively, correlating with transients that follow the start of each ON or OFF segment of the overall magnetic actuation waveform sequence. Knowing *a priori* that the microsphere defined by 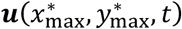 would have experienced the largest accelerations during magnetic actuation, we classified each imaging frame within the original data set of the prototype FC experiment as being elastostataic if 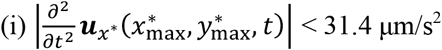, and if 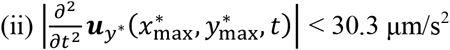, and if (iii) the frame was continuous within a sequence of more than 15 other elastostatic imaging frames. If all three conditions were not met, the frame was classified as being *dynamic* and therefore potentially in violation of the elastostatic assumptions inherent to our solution of the stress traction vector field. Dynamic time intervals for our prototype FC experiment are shown in **Fig. 2(f)** and **Fig. 3**. Similarly, images of diminished brightness are used to indicate dynamic imaging frames in **Vid. 4** and **Vid. 5**. Based on the above analysis for ±6SD acceleration bounds, we argue that the vast majority of imaging frames collected for our prototype FC experiment are elastostatic, and therefore associated with valid solutions for ***T***(*x*\*, *y*\*, *t*), ***F***(*t*), *ρ*(*x*\*, *y*\*, *t*), and *U*(*t*). Alternatively, reduced upper and lower acceleration bounds could be employed using this same algorithm, e.g., ±3 standard deviations of the recorded acceleration data, and accordingly, this would result in a decreased number of imaging frames being classified as elastostatic.

**FIG. 4.**
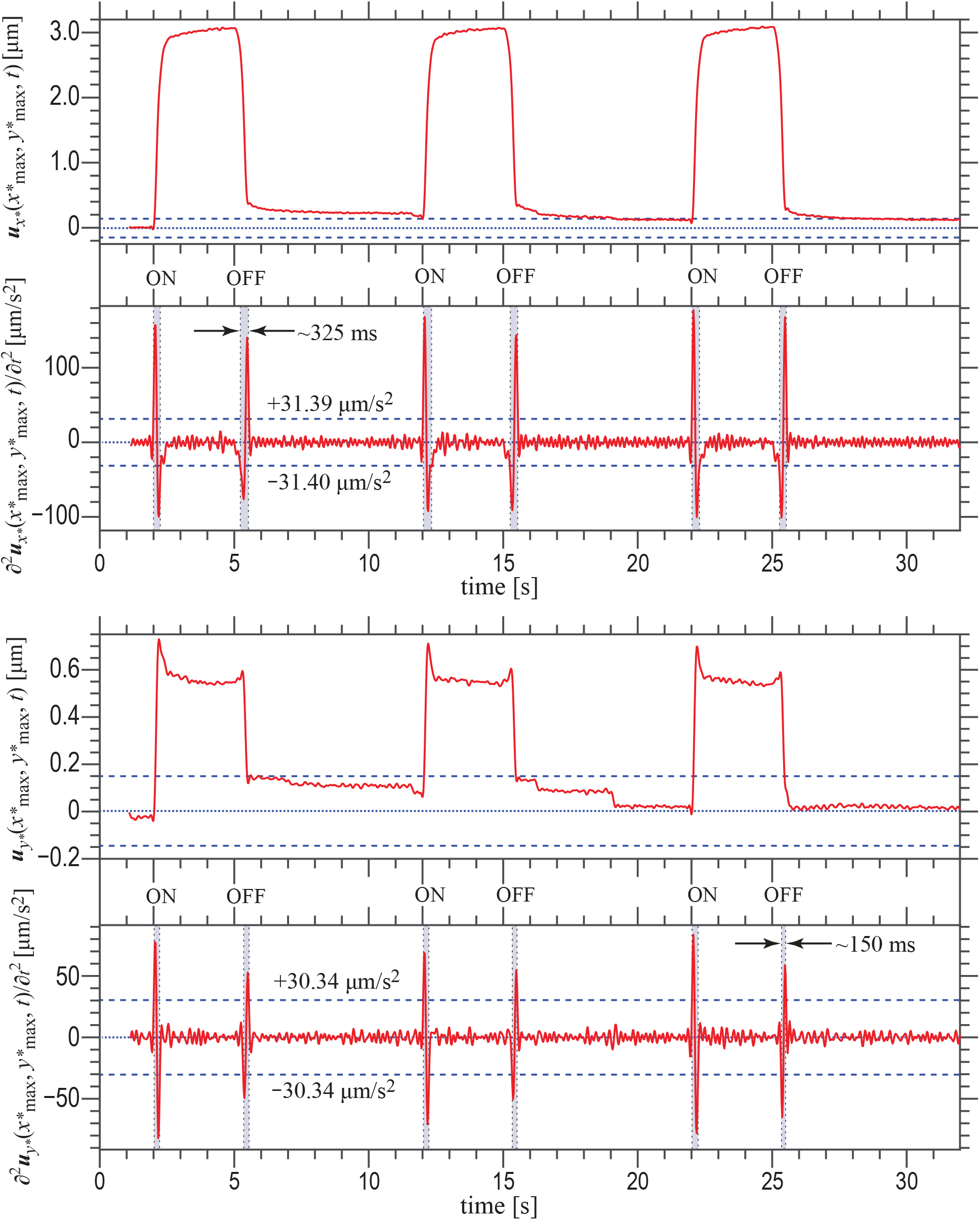
Microsphere dynamics in the FC mode bead-on-gel experiment. Plots showing time-resolved vectoral components in the *x*\*- and *y*\*-directions of the displacement, 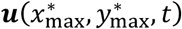, and acceleration, 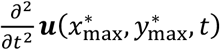, of the microsphere embedded within the collagen substrate that was subject to the overall largest displacement during magnetic actuation in the prototype FC mode bead-on-gel experiment. Horizontal dashed lines in each plot represent the 6 standard deviations (6SD) upper- and lower-bounds representative of the physical detection limits for measured microsphere displacements and accelerations, as derived from a null force control experiment (see Sec. IV.A.1). Gray-shaded vertical boxes indicate *dynamic* time intervals during which inertial contributions to the force balance of the boundary-value problem used to find solutions for the stress traction vector field are likely to be non-negligible.

#### 2. Displacement-control mode experiment

In most MT-based experiments, displacements of the superparamagnetic bead are small, i.e., <5 μm, and associated with actuation forces on the order of 5 nN or less. However, in exploring the mechanobiology of an epithelial sheet, it is conceivable that selective application of large forces (>5 nN) and/or large displacements (>5 μm) might be required to interrogate force transmission via specific cell-cell or cell-matrix anchoring junctions. Such is the case in our work, and thus the impetus for the development of the displacement-control mode MT-DTM/TFM experiment presented here. As a prototype demonstration of a DC experiment, 4.5 μm-diameter, fibronectin-coated superparamagnetic beads were allowed to attach to a ~700 μm thick, 1.0 mg/mL type I collagen substrate embedded with an embedded surface layer of fluorescent microspheres (see Secs. III.B and III.D). With the microscope configured for 30X magnification, two separate FC experiments were performed using 3-cycle 3 s/7 s ON/OFF magnetic actuation waveform sequences, each with maximum nominal ON magnetic flux density setpoints of 175 Gs, exactly as described in Sec. SI.A.1 of the **Supplementary Material**. For both experiments, the initial *xyz*-spatial location of the superparamagnetic bead relative to the needle tip of the MT device was *x*(0) = 20.0 μm, *y*(0) = 0 μm, and *z*(0) = 13.0 μm. DIC and epifluorescence images collected during the second FC experiment were used to determine the apparent elastic modulus for the collagen substrate (*E* = 52.73 Pa, assuming *u* = 0.4) using a global ROI set to the central 1024 × 1022 pixels of each image.

A second superparamagnetic bead a few fields of view away from this original bead was then identified. With the microscope now configured for 20X magnification and DIC fast time-lapse image acquisition at 40 fps (2048×2044 pixels per frame), the bead was magnetically clamped to the needle tip by actuating the MT device with a magnetic flux density setpoint of 175 Gs. The needle was then *manually* translated away from this initial position in 1 μm-step increments, holding each position for ~1 s before proceeding with the next incremental step until a maximum needle translation of 10 μm was reached. After a ~2 s hold at this maximum displacement, the needle was manually translated back to its initial position in an identical stepwise fashion. After a brief 1 s to 5 s pause, the MT was actuated to achieve ***F***_MT_(*t*) = 0 nN, releasing the superparamagnetic bead from the needle tip. The exact same sequence of events was then carried out with the microscope configured for fast time-lapse epifluorescence imaging. More specific procedural details for this prototype DC MT-DTM/TFM experiment can be found in Sec. SI.B.1 of the **Supplementary Material**. For a static null displacement control experiment (with respect to collagen substrate motion), the needle was positioned near the same superparamagnetic bead of interest such that δ < 1 μm. The MT device was actuated such that ***F***_MT_(*t*) = 0 nN, i.e., a null magnetic clamping force. Serial fast time-lapse image acquisitions at 40 fps (2048×2044 pixels per frame) were then performed in DIC and epifluorescence imaging modes while the needle tip was translated to and from the same maximum displacement of ~10.0 μm, exactly as done in the model DC experiment. The substrate remained undeformed throughout the duration of the null displacement control experiment.

Following data collection, all image data was analyzed according to the methods presented in Sec. SI.B.2 of the **Supplementary Material**. Real-time videos of the DIC image data (full field and zoom) and the epifluorescence image data (full field), as well as videos for ***u***(*x*\*, *y*\*, *t*), ***T***(*x*\*, *y*\*, *t*), and *ρ*(*x*\*, *y*\*, *t*) for the prototype DC MT-DTM/TFM experiment can be found in **Vid 11**, **Vid 12**, **Vid 13**, **Vid 14**, **Vid 15**, and **Vid 16**, respectively, the latter three videos computed assuming *E* = 52.73 Pa and *u* = 0.4. The analogous set of videos for the null displacement control experiment can also be found in **Vid 17**, **Vid 18**, **Vid 19**, **Vid 20**, **Vid 21**, and **Vid 22**, respectively, the latter three videos also computed assuming *E* ~ 52.73 Pa and *u* = 0.4. Quantitative determination of imaging intervals where microsphere dynamics might preclude implementation of our elastostatic TFM analysis was not done for this DC experiment as was previously shown for the prototype FC experiment (see Sec. IV.A.1). Although our null displacement control experiment provides a data set amenable to such analysis, identification of these time intervals does not offer any new insight into understanding this potential limitation of our DC MT-DTM/TFM methodology.

In contrast to an FC experiment, the prototype DC experiment described here does not employ a programed and measured magnetic actuation waveform that can be used to temporally correlate data from the serially acquired DIC and epifluorescence image sets. Translation of the needle tip was *manually* initiated for both imaging sequences and did not necessarily start at the same raw time point during each respective fast time-lapse image acquisition. As such, we chose to correlate the independent data sets by temporally aligning the first significant microsphere displacement observed during the epifluorescence imaging sequence to the first significant superparamagnetic bead displacement observed during the DIC imaging sequence. Experimentally significant displacements were determined via analysis of our null displacement control experiments, as can be found in the **Supplementary Material**(**Fig. S5** and **Fig. S6**). In brief, we first searched the epifluorescence data set to find the raw time point, *t*_EPI_, at which 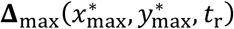 first becomes ≥ 0.15 μm, i.e., the raw time, *t*, marking the first physically significant microsphere displacement measurement (see **Fig. S6**). We then determined *t*_DIC_, the first time point at which **Δ_MT_**(*t*) ≥ 0.28 μm in the DIC imaging data, i.e., the time marking the first physically significant superparamagnetic bead displacement (see **Fig. S5**). Knowing *t*_DIC_ and *t*_EPI_, we temporally shifted the entire epifluorescence data set, re-defining the temporal variable for this data set, *t*, as *t* = *t*_r_ + (*t*_DIC_ − *t*_EPI_). With this alignment of the epifluorescence and DIC image data, the first experimentally significant microsphere and superparamagnetic bead displacements are temporally coincident. For the prototype DC experiment shown here, *t*_EPI_ = 2.65278 s while *t*_DIC_ = 5.33058 s. Time stamps shown in **Vid 13**, **Vid 14**, **Vid 15**, and **Vid 16** have all been shifted by 2.6778 s.

Alignment of the data sets in this manner assumes that superparamagnetic bead and substrate microsphere displacements are temporally coincident, though this is not known to be true *a priori*. However, in our FC experiments, Δ_MT_ and Δ_max_ were observed to be temporally coincident despite differences in the observed magnitudes of these displacements (see **Fig. 2(e)**). Thus, we believe this to be a reasonable assumption for analyses presented here. In future DC experiments, we plan to use a piezo nanopositioner to control the translation of the needle tip, similar to the setup described for MT-based pick and place magnetic particle assembly^58^. DIC and epifluorescence imaging data will then be temporally correlated by measurements of the actuator’s position acquired in parallel with the TTL pulses marking exposures of the sCMOS camera during each respective DIC or epifluorescence fast time-lapse imaging sequence.

A 6-panel plot that highlights key findings from the prototype DC experiment is shown in **Fig. 5**. A similar 6-panel plot for the static null displacement control experiment and a graphical depiction of our computation of the L2 regularization parameters, λ2_ON_ and λ2_OFF_, for the prototype DC experiment detailed in **Fig. 5** can be found in **Fig. S7** and **Fig. S8**, respectively, of the **Supplementary Material**. In comparing the displacement of the superparamagnetic bead, Δ_MT_(*t*), and the displacement of the microsphere subject to the largest displacement during translation of the needle tip, Δ_max_(*t*), we first note that the data sets are not perfectly correlated in time (see **Fig. 5(e)**). We attribute this to the fact that displacement control was achieved via manual translation of the needle tip using our three-axis micromanipulator. With manual control, translation step and hold times were not identical for the DIC and epifluorescence image sets. Despite this fact, several important findings are still evident from the coupled data sets. Foremost, we see a discrepancy between the maximum of Δ_MT_(*t*) (~10.3 μm) compared to the maximum of Δ_max_(*t*) (~8.9 μm) observed during the DC experiment. Similar to our FC experiment (see Sec. IV.A.1), the difference between maximums in Δ_MT_ and Δ_max_ (~1.4 μm) is less than the radius of the superparamagnetic bead. As previously hypothesized for our FC experiments, we also speculate here that the superparamagnetic bead can both rotate and translate during application of the magnetic clamping force exerted by the MT device. Relative amounts of rotation and translation are likely dependent on the state of conformational contact between the bead and the collagen substrate. Another possibility to consider is that fibronectin coating of the superparamagnetic bead results in the formation of short fibronectin fibrils^59^. As the bead is displaced, stretching of fibronectin fibrils could potentially account for the some of the discrepancy between Δ_MT_ and Δ_max_, depending on the relative mechanical compliances of collagen and fibronectin fibrils.

**FIG. 5.**
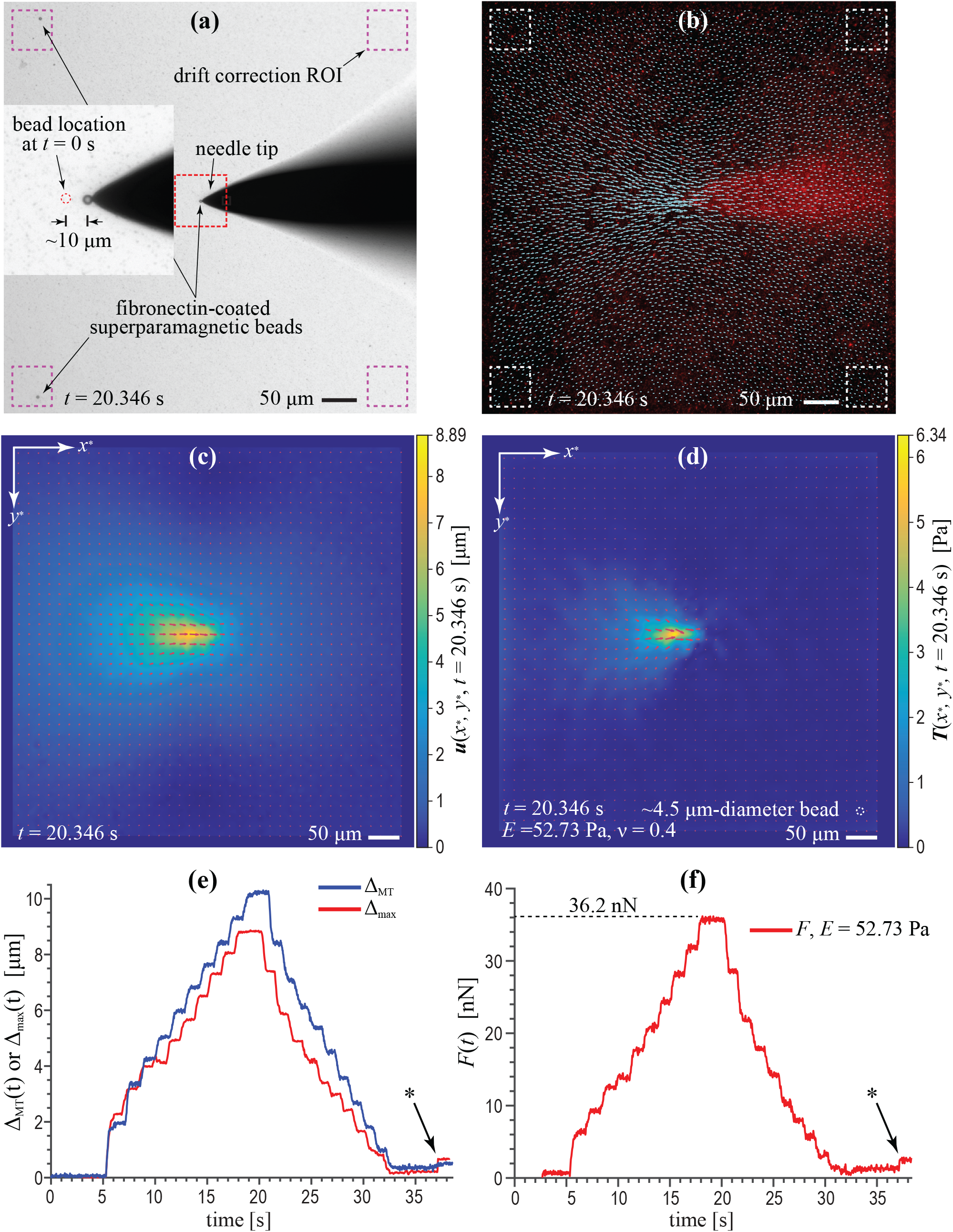
DC mode bead-on-gel experiment. Six-panel figure highlights data from a prototype DC mode bead-on-gel experiment involving a fibronectin-coated 4.5 μm-diameter superparamagnetic bead attached to the surface of a ~700 μm-thick, 1.0 mg/mL type I collagen substrate containing a surface layer of covalently attached red fluorescent microspheres. **(a)** DIC image showing the superparamagnetic bead near its peak displacement following a ~10 μm translation of the needle tip. The inset image shows a magnified view of the region denoted by the red dashed square. **(b)** Texas Red epifluorescence image with overlaid quivers representing the displacement vectors of individual microspheres at the same time point. Dashed boxes in **(a)** and **(b)** denote the ROIs used for drift correction. The displacement field, ***u***(*x*\*, *y*\*), and stress traction vector field, ***T***(*x*\*, *y*\*), computed for the microsphere displacement field displayed in **(b)**, are shown in **(c)** and **(d)**, respectively, where for aid of visualization, the white dashed circle in **(d)** represents a ~4.5 μm-diameter circle. A plot of the displacement of the microsphere subject to the largest displacement during translation of the needle tip, Δ_max_(*t*), versus superparamagnetic bead displacement, Δ_MT_(*t*), is shown in **(e)**. A plot of the magnitude of the integrated total traction force vector, *F*(*t*), is shown in **(f)**, where *F*(*t*) has been computed assuming *υ* = 0.4 and *E* = 52.73 Pa. The asterisks (*) shown in **(e)** and **(f)** denote the time points at which the magnetic clamping force was removed, releasing the bead from the needle tip.

Though not observed in this experiment, in other trial DC experiments, we did observe (with DIC imaging) relative sliding contact motion between the superparamagnetic bead and the needle tip that occurred during the translation of the needle tip away from its initial position following application of the magnetic force clamp. Thus, it is possible that relative bead-tip motion contributed to the non-uniform initial displacement steps in Δ_max_(*t*) that can be observed for 5 s < *t* < 13 s in **Fig. 5(e)**. However, as the substrate becomes increasingly deformed, the magnitude of displacement steps in Δ_max_(*t*) and Δ_MT_(*t*) become nearly identical, suggestive that no further relative motion occurs between the superparamagnetic bead and the needle tip. As can be seen in **Fig. 5(e)**, small residual step displacements in both Δ_max_(*t*) and Δ_MT_(*t*) are observed following demagnetization of the needle tip and removal of the clamping force at *t* ≈ 37 s (also see **Fig. 5(f)**). Assuming that the collagen behaves as a linear elastic substrate, this displacement could be accounted for by hysteresis in the micromanipulator used to translate the needle tip. Alternatively, assuming no hysteresis in translation of the needle tip, residual step displacements might be attributed to the viscous and/or plastic constitutive material properties of the collagen substrate in response to the applied deformation cycle.

With regards to our continuum models of the measured displacement field and computed stress traction vector fields, note that although three superparamagnetic beads were present within the field of view for this DC experiment (see **Fig. 5(a)**), substrate deformations and traction stress were localized only to the near field of the bead that was magnetically clamped to the needle tip, as can be seen in **Figs. 5(c)** and **5(d)**, respectively. Due to the needle’s concentrated magnetic field gradients, our MT device possesses the ability to *selectively* clamp a specific bead of interest without introducing forces on neighboring superparamagnetic beads positioned >385 μm away from the needle tip. Smaller neighborhoods may be possible, but the exact spatial selectivity of our MT device was not further quantified in this work.

With that understanding, compare the maximum stress generated in our prototype DC experiment, ~6.34 Pa (as shown in **Fig. 5(d)**), to the maximum stress generated in our prototype FC experiment, ~18.2 Pa (as shown in **Fig. 2(d)**). Admittedly, this result is physically inconsistent with what one would expect given that the deformation in the DC experiment was much larger than that observed for the FC experiment, even considering the known differences in apparent elastic moduli of the collagen substrates used for these two independent experiments (*E* ~ 52.73 Pa for the DC experiment, whereas *E* ~ 76.23 Pa for the FC experiment). However, it is important to recognize that our solution for the stress traction vector field was *unconstrained*, meaning that we did not restrict solutions for the traction stress field to consist only of tractions localized to within the area of bead-substrate contact. By happenstance, the peak traction stress solution for the FC experiment predicted a small localized traction stress confined to a 4.5 μm-diameter circular area, roughly the same size of a superparamagnetic bead. In contrast, the peak traction stress solution for the DC experiment predicted a traction stress field localized to a linear streak-like area with a length of ~3 superparamagnetic bead diameters. In future experiments, we will consider adopting an iterative solution scheme that constrains stress tractions to an area of presumed conformational contact between the collagen substrate and the superparamagnetic bead, analogous to the methodology originally described by Butler et al. for constraining cellular focal adhesion tractions to spatial locations present within the projected boundary of cell^60^.

As a final discussion point, we note that our prototype DC experiments tended to fail by one of three mechanisms: (i) mechanical fracture within the collagen substrate, (ii) detachment of the superparamagnetic bead from the collagen substrate, or (iii) the development of substrate traction stresses that ultimately exceeded the magnetic clamping force holding the bead to the needle tip. Fracture of the collagen substrate was only observed for extremely large deformations of soft gels with collagen concentrations of ≤1 mg/mL. When fracture occurred, grossly visible amounts of collagen debris could be observed sticking to the superparamagnetic bead. For our fibronectin-coated beads (see Sec. III.D), bead detachment from the collagen substrate proved to be a rare event when beads were incubated for >12 hours at 37°C to allow sufficient time to establish conformal contact via hydrophobic adhesive interactions between fibronectin and collagen. Consequently, bead-tip separation represented the most common mode of experimental failure.

As can be seen in **Fig. 5(f)**, the total traction force vector, *F*, reached a maximum of ~36.2 nN during this prototype DC experiment, suggesting a bead-tip clamping force of at least the same magnitude. In other DC experiments using 1.0 mg/mL collagen gels, our TFM analysis suggested *F* = ~43.0 nN for peak collagen displacements of Δ_max_ = ~11.0 μm. As such, we believe that our MT device operates with a bead-tip clamping force of at least 40 nN. Although maximum displacements of Δ_MT_ = ~20.0 μm were possible with 1.0 mg/mL gels, unfortunately, tracking of substrate microspheres for these experiments proved to be exceedingly difficult. Moreover, even if we were able to track substrate microspheres for these large displacement experiments, the finite deformations resultant in the collagen gel would preclude accurate modeling of traction stresses using our TFM code given that our computational solution for *T* is based on a theory of infinitesimal deformations. In future work, we plan to explore alternative needle tip geometries and different preparations of both collagen gels and superparamagnetic and/or ferromagnetic beads in an attempt to better define the maximum range of bead-tip clamping forces that characterize the DC mode of operation for our integrated MT-DT/TFM apparatus.

### B. Keratinocyte Force Transmission

Type I collagen gel substrates, 2.0 mg/mL, ~500 μm to 600 μm thick and embedded with a surface layer of fluorescent microspheres, were prepared as previously described (see Sec. III.B). Multicellular sheets of primary normal human epidermal keratinocytes were reconstituted on these collagen substrates and cultured using either low [Ca^2+^] or high [Ca^2+^] KSFM for ~24 hours (see Sec. III.C). Phase contrast images of the keratinocyte sheets at low and high [Ca^2+^] conditions and epifluorescence images of their respective underlying fluorescent microsphere distributions are shown in **Fig. 6**. As can be seen in **Figs. 6(a)** and **6(c)**, keratinocytes adherent to 2.0 mg/mL gels exhibited grossly normal morphology and were found to survive for ~72 hours in culture, at which time our experiments were terminated. Keratinocytes grown on gels ≤1.0 mg/mL did *not* exhibit normal cell morphology or viability. Under low [Ca^2+^] conditions, keratinocytes within the multicellular sheet are known to lack true cell-cell adherens junctions or desmosomes^2,15,32–36^. As can be seen **Fig. 6(b)**, individual keratinocytes at low [Ca^2+^] conditions-mechanically coupled to the underlying substrate via focal adhesion contacts-each form a distinct cell-sized zone of increased microsphere density in their immediate subjacent collagen substrate, referred to here as a *microsphere footprint*. Whether or not this keratinocyte-organized microsphere footprint reflects true re-organization and compaction of collagen fibrils remains an open question subject to further investigation. Regardless, microsphere footprints of individual cells were similar in size and microsphere density whether or not a keratinocyte was present within the central area or the peripheral edge of a multicellular sheet. In stark contrast, keratinocytes cultured under high [Ca^2+^] conditions-known to be mechanically coupled to one another via both cell-cell adherens junctions and desmosomes^2,15,32–36^-exhibited a dramatic intensification of microsphere footprints within cells present at the peripheral edge of a multicellular sheet (see **Fig. 6(d)**). Qualitatively, these data suggest that under low [Ca^2+^] conditions, keratinocytes within a multicellular sheet generate *independent* cellular traction stresses to maintain adhesion to the underlying substrate, whereas for keratinocytes cultured at high [Ca^2+^], cellular traction stresses are *cooperatively* localized to the periphery of the multicellular sheet. Collectively, these observations are congruent with the traction stress measurements and modelling that has previously been reported for small colonies of murine keratinocytes^24,61^.

**FIG. 6.**
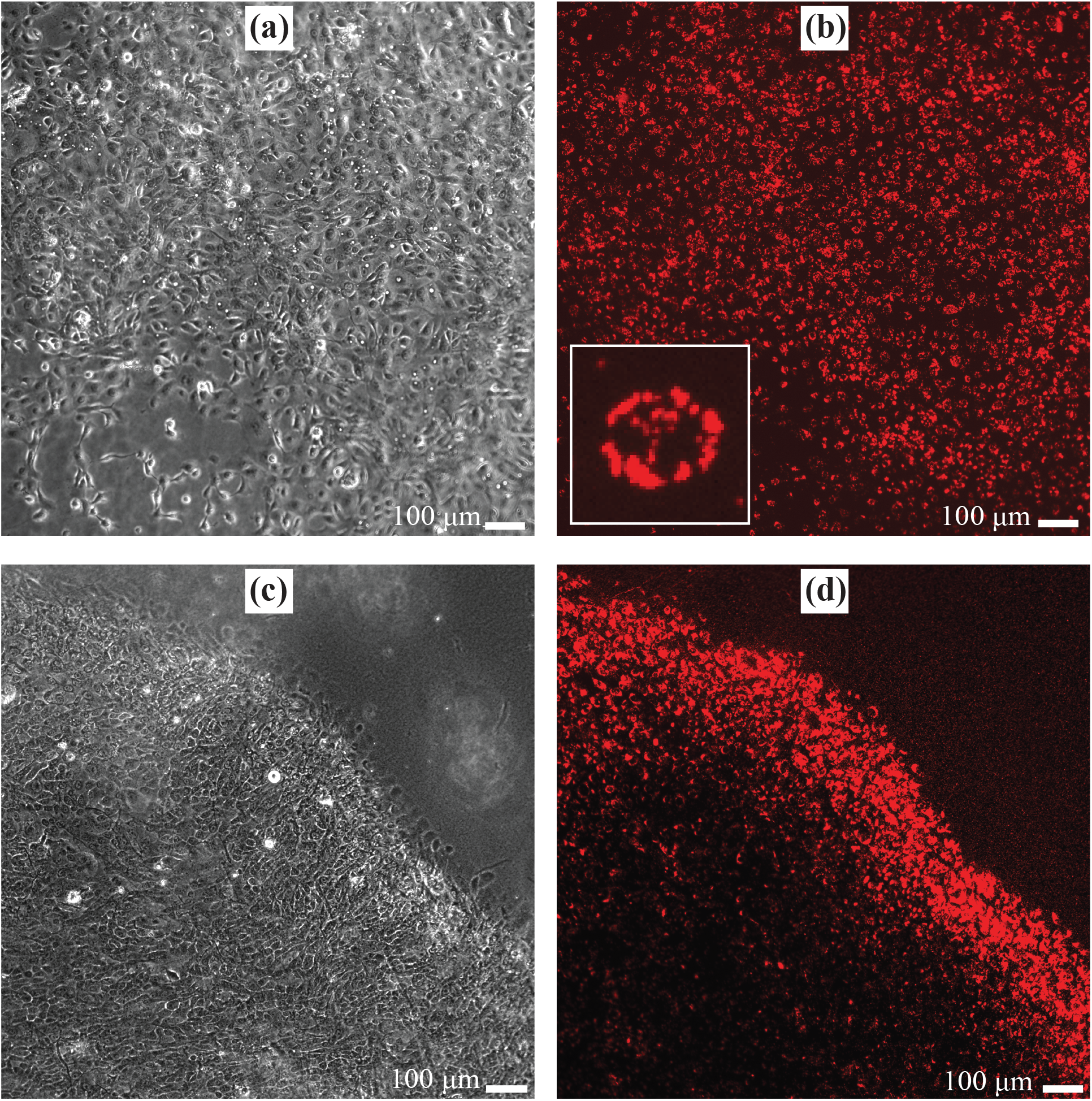
Paired phase contrast and Texas Red epifluorescence images of select areas of reconstituted multicellular keratinocyte sheets cultured for ~24 hours with low [Ca^2+^] **(a, b)** or high [Ca^2+^] **(c, d)** conditions. Phase contrast images allow visualization of individual and collective keratinocyte morphology, whereas the epifluorescence images capture microsphere distributions within the underlying 2.0 mg/mL type I collagen substrate. The image inset in **(b)** represents a magnified view of the microsphere footprint characteristic of an isolated keratinocyte cultured using low [Ca^2+^] medium.

One multicellular keratinocyte sheet cultured in low [Ca^2+^] medium was selected for further interrogation. Fibronectin-coated superparamagnetic beads were allowed to attach, at random, to cells within this sheet using the protocol described in Sec. III.D. A superparamagnetic bead located in a cell-free area adjacent to the keratinocyte sheet was subjected to our prototype FC MT-DTM/TFM experiment as described in Sec. IV.A.1. From this data, we quantified an apparent elastic modulus of *E* ~ 138.3 Pa (assuming *u* = 0.4) for this 530 μm-thick 2.0 mg/mL collagen substrate, employing the analysis described in Sec V.A.1. Next, in what is referred to as a bead-on-cell experiment, a superparamagnetic bead attached to the apical surface of a keratinocyte present at the periphery of the multicellular sheet was identified and subjected to two sequential 3-cycle 4 s/7 s ON/OFF FC mode actuation cycles with maximum nominal ON magnetic flux density setpoints of 175 Gs, exactly as described in Sec. SI.A.1 of the **Supplementary Material**. A tiled phase contrast image of the multicellular sheet and the relative location of the specific keratinocyte interrogated within this sheet can be found in the **Supplementary Material**(see **Fig. S9**). The initial *xyz*-spatial location of the superparamagnetic bead relative to the needle tip of the MT device for this experiment was *x*(0) = 11.0 μm, *y*(0) = 0 μm, and *z*(0) = 13.0 μm (δ(*t*_0_) ~ 17.0 μm). Image data from the second 3-cycle FC MT-DTM/TFM experiment were analyzed according to the methods presented in Sec. SI.A.2 of the **Supplementary Material**. Real-time videos of the DIC and epifluorescence imaging data for this bead-on-cell experiment can be found in **Vid 23** and **Vid 24**, respectively. Videos for ***u***(*x*\*, *y*\*, *t*), ***T***(*x*\*, *y*\*, *t*), and *ρ*(*x*\*, *y*\*, *t*) can be found in **Vid 25**, **Vid 26**, and **Vid 27**, respectively, all computed assuming *E* = 138.3 Pa and *u* = 0.4. A graphical determination of the L2 regularization parameters used for the solution of ***T*** and *ρ* in this experiment can also be found in the **Supplementary Material**(**Fig. S10**). Though possible, quantitative determination of imaging intervals where microsphere dynamics prevent implementation of an elastostatic TFM analysis was not done for this experiment as was previously shown for the prototype FC experiment (see Sec. IV.A.1).

Important findings from this bead-on-cell experiment investigating force transmission through a keratinocyte cultured under low [Ca^2+^] conditions are graphically summarized in **Fig. 7**. Here, Figs. **7(a)**, **7(b)**, **7(c)**, and **7(d)** represent the DIC image, epifluorescence image, ***u***(*x*\*, *y*\*), and ***T***(*x*\*, *y*\*), respectively, captured or calculated at *t*_OFF_ for cycle #3 (*t* ≈ 27.14 s) of the overall magnetic actuation waveform. Magnified views of ***u*** and ***T*** within the white dashed boxes shown in **Figs. 7(c)** and **7(d)**, respectively, are shown in **Fig. 8**. In the discussion that follows, it is extremely important to note that ***u*** measured for this bead-on-cell experiment represent substrate displacements that arise in response to magnetic actuation of the superparamagnetic bead attached to the apical surface of keratinocyte #1 (see **Fig. 7(a)**). In other words, substrate displacements, ***u***, are defined with respect to a reference state that consists of a substrate that has already been mechanically deformed by the overlying cell layer, *not* an undeformed substrate. Consequently, in this analysis, ***T*** represents the *incremental* stress traction field that develops in response to magnetic actuation, *not* the active stress tractions generated by cells in establishing adhesive contact with the underlying collagen substrate.

**FIG. 7.**
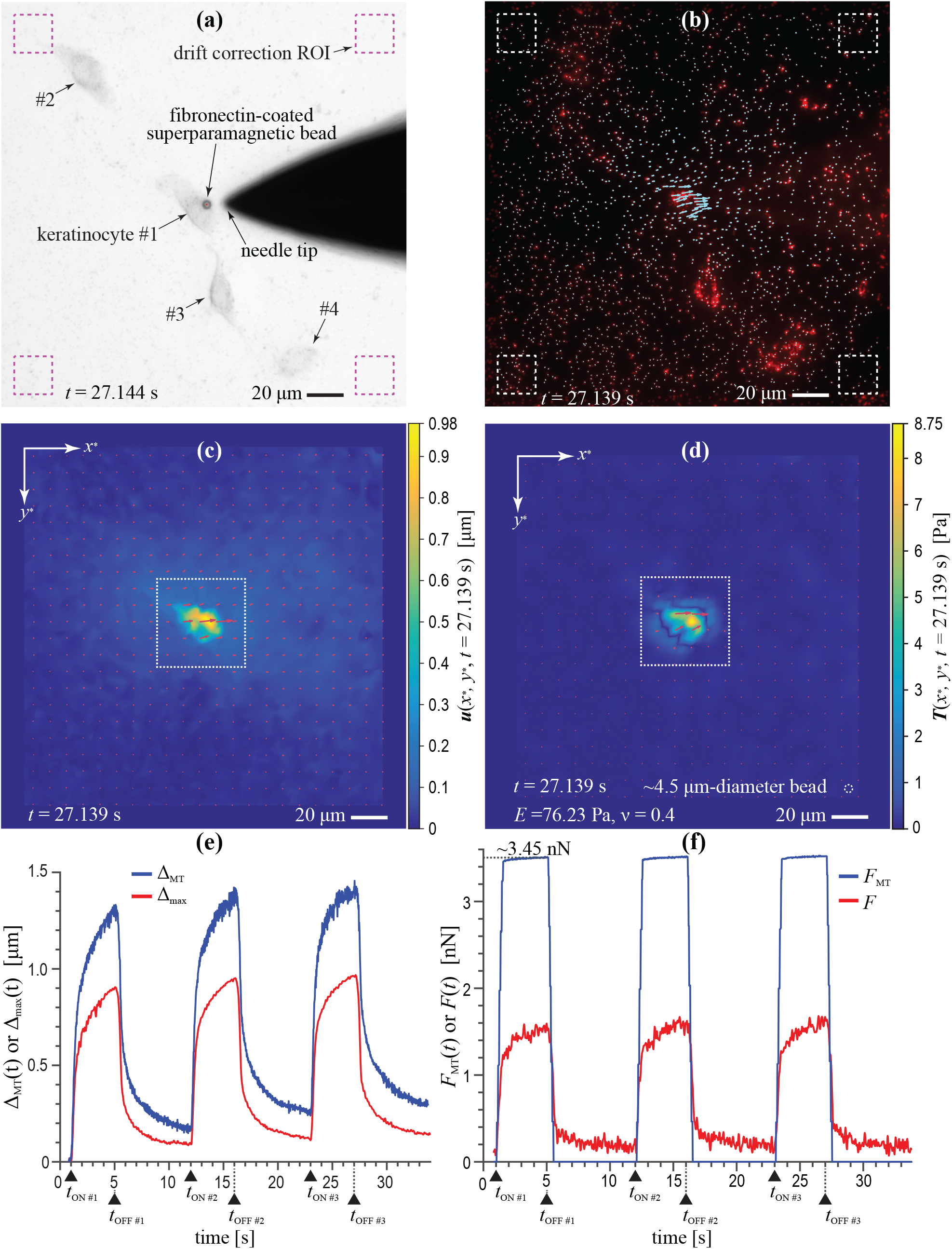
FC mode bead-on-cell experiment. Six-panel figure highlights data from an FC mode bead-on-cell experiment involving a fibronectin-coated 4.5 μm-diameter superparamagnetic bead attached to the apical surface of normal human epidermal keratinocyte that is adherent to a ~530 μm-thick, 2.0 mg/mL, type I collagen substrate containing a surface layer of covalently attached red fluorescent microspheres. The keratinocyte is present at the peripheral edge of a multicellular sheet reconstituted under low [Ca^2+^] conditions (see **Fig. S9** of the **Supplementary Material**). **(a)** DIC image showing the superparamagnetic bead at its peak displacement near *t*_OFF_ of actuation cycle #3. **(b)** Texas Red epifluorescence image with quivers that represent the displacement vectors of individual microspheres within the surface of the collagen substrate at this same time point. The displacement field, ***u***(*x*\*, *y*\*), and the corresponding stress traction vector field, ***T***(*x*\*, *y*\*), are shown in **(c)** and **(d)**, respectively, where for aid of visualization, the white dashed circle in **(d)** represents a ~4.5 μm-diameter circle. The white dashed square boxes in **(c)** and **(d)** denote the area of collagen substrate subjacent to keratinocyte #1. Magnified views of this region are contained in **Fig. 8**. A plot of the displacement of the microsphere subject to the largest displacement during magnetic actuation, Δ_max_(*t*), versus superparamagnetic bead displacement during magnetic actuation, Δ_MT_(*t*), is shown in **(e)**. A plot of the magnitude of the component of the magnetic actuation force vector in the *xy*-plane of the collagen substrate, *F*_MT_(*t*), versus the magnitude of the integrated total traction force vector, *F*(*t*), is shown in **(f)**. Both ***T***(*x*\*, *y*\*, *t*) and *F*(*t*) were computed assuming *υ* = 0.4 and *E* = 138.3 Pa. Black triangles in **(e)** and **(f)** denote *t*_ON_ and *t*_OFF_ for each actuation cycle.

**FIG. 8.**
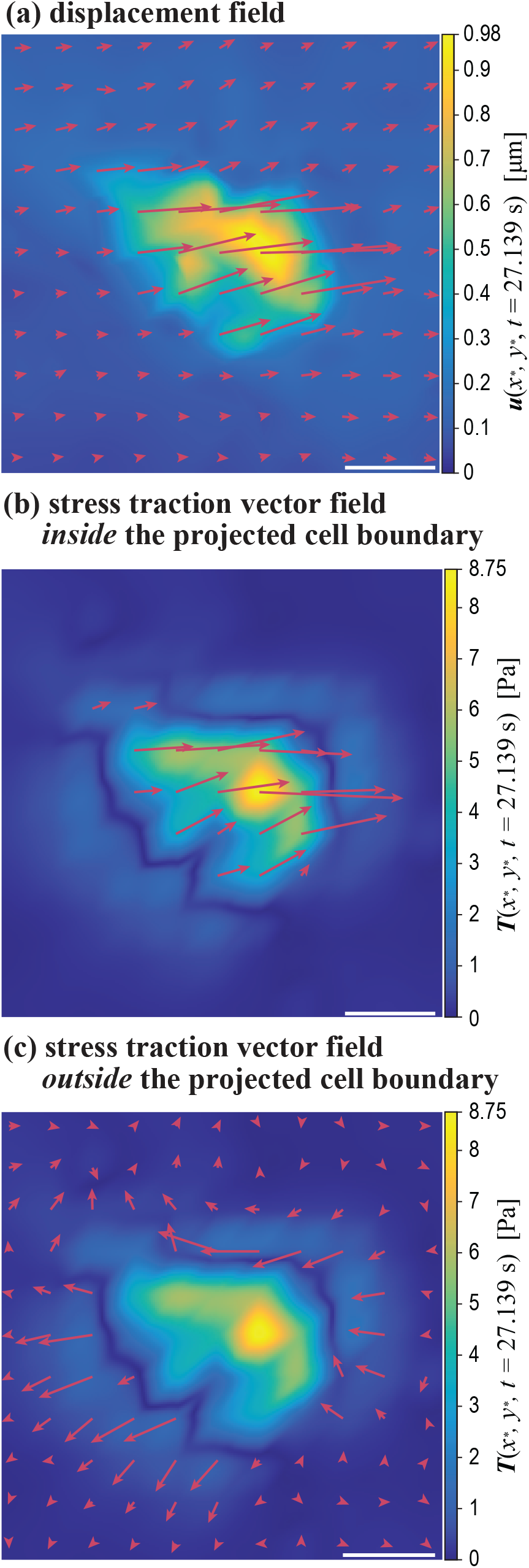
***u***(*x*\*, *y*\*, 27.139 s) and ***T***(*x*\*, *y*\*, 27.139 s) underlying keratinocyte #1. Images depict magnified views of **(a)** the displacement field, ***u***(*x*\*, *y*\*), and **(b**, **c)** the stress traction vector field, ***T***(*x*\*, *y*\*), for the area of substrate subjacent to keratinocyte #1 at near *t*_OFF_ of actuation cycle #3 of the FC mode bead-on-cell experiment detailed in **Fig. 7**(*t* = 27.139 s). For ease of visualizing the local directionality of the stress traction vector field, traction quivers *external* to the projected keratinocyte boundary have been removed in **(b)**, whereas quivers *within* the area of substrate subjacent to the cell boundary have been omitted in **(c)**. Scale bars = 10 μm.

Foremost, as can be seen in **Fig. 7(a)**, note that there are four keratinocytes present within the field of view for this experiment, with keratinocyte #1 having the fibronectin-coated superparamagnetic bead attached to its apical surface. Keratinocyte #3 is in close apposition to keratinocyte #1, and it is possible that their cell membranes are in direct physical contact. As clearly visible in **Vid 24**, one can appreciate that each of these four keratinocytes has created a microsphere footprint in their immediate subjacent collagen substrate *prior to magnetic actuation*. Now, observe how in both **Fig. 7(b)** and **Vid 24** that only keratinocyte #1 develops obvious displacements in its subjacent collagen substrate in response to magnetic actuation of the superparamagnetic bead. Perhaps more clearly evident in ***u***(*x*\*, *y*\*, *t*), during magnetic actuation, the subjacent collagen develops several focal peaks in displacement that are only localized to substrate areas within the projected boundary of keratinocyte #1 (see **Fig. 7(c)** and **Vid 25**). No significant displacements are observed in the collagen substrate underlying keratinocytes #2, #3, and #4. In comparing the displacement field that develops during this bead-on-cell experiment to the displacement field of the bead-on-gel experiment, note how the latter exhibits a single local displacement peak that monotonically decays out into the surrounding substrate (see **Fig. 2(b)** and **Vid 3**). In contrast, the displacement field for the bead-on-cell experiment shown in **Fig. 8(a)**, though continuous, exhibits a very sharp gradient in ***u***(*x*\*, *y*\*) as displacements vary from 0 μm in the substrate immediately adjacent to the cell border, but then abruptly rise to ~1 μm within the collagen subjacent to the cell.

With regards to the mechanical response of the superparamagnetic bead, Δ_MT_(*t*), we see more evidence of an overall viscoelastic response in the bead-on-cell compared to the bead-on-gel experiment. With a cycle #1 peak displacement of ~1.3 μm (**Fig. 7(e)**) in response to a ~3.45 nN load (**Fig. 7(f)**), the data for our keratinocyte experiment are both qualitatively and quantitatively similar to the mechanical response observed for a 4.5 μm-diameter fibronectin-coated Dynabead™ that has been attached to a murine embryonic fibroblast and actuated with an MT device^62^. In comparing Δ_MT_(*t*) to Δ_max_(*t*), we note that the observed difference between Δ_MT_(*t*_OFF_) and Δ_max_(*t*_OFF_) for each of the three actuation cycles varies from ~0.4 μm to ~0.5 μm (**Fig. 7(e)**). However, Δ_MT_(*t*) and Δ_max_(*t*) are temporally coincident without any observable phase shift between the two displacement signals, which is indicative of elastic mechanical coupling between the bead and the underlying collagen substrate. Collectively, our findings strongly suggest that in this FC bead-on-cell experiment, the force applied to the superparamagnetic bead during magnetic actuation was transmitted from the apical surface of the keratinocyte through the cytoskeleton and ultimately to focal adhesion contacts along the interface with the subjacent collagen substrate. Evidence of cell-cell force transmission is lacking.

As one would expect from the displacement field shown in **Fig. 8(a)**, the solution for ***T***(*x*\*, *y*\*, *t* = 27.139 s) exhibits focal traction stresses within the collagen subjacent to the cell that are directionally correlated with the underlying displacement field (see **Figs. 8(b)** and **8(c)**). However, these tractions are also continuous with an annular zone of ~0 Pa traction stress that is surrounded by an outer zone of stress tractions with a directionality opposite of the central cell tractions. Sometimes referred to as “ringing” in TFM, this finding in our solution for ***T*** is a mathematical consequence of the input displacement field regardless of whether the solution was computed using FTTC or the boundary element method (see **Fig. S11** of the **Supplementary Material**)^40^. Low pass filters in Fourier space are often used in FTTC-based approaches to TFM to remove high frequency noise from the gridded displacement data^39^. However, in this instance, ringing in the computed solution for ***T*** is a consequence of an actual measured step-like gradient in ***u***, and therefore filtering out the high-frequency content of ***u*** would not be appropriate.

Lastly, we note that in the bead-on-gel experiment (see Sec. IV.A.1), the collagen substrate was shown to exhibit mechanical properties that approximate a linear elastic solid. However, in this bead-on-cell experiment, individual keratinocytes adherent to the gel have presumably re-organized the fibrillar structure of the collagen, as evidenced by the increased microsphere density in their immediate subjacent substrate. As such, we postulate that the displacement gradient observed in our bead-on-cell experiment developed as a consequence of localized incremental traction stresses applied to a fibrillar collagen substrate with constitutive mechanical properties that are *spatially non-homogenous*. Practically speaking, collagen substrates that have undergone significant remodeling by adherent cells no longer behave as a linear elastic material, invalidating the assumptions inherent to our TFM stress traction solutions. However, until higher-order TFM models are developed that allow for spatially non-homogeneous substrates, to a first-order approximation, we pose that our solutions for ***T*** still stand as semiquantitative observations that can still provide potentially useful insights into future investigations of force transmission in multicellular keratinocyte constructs.

## V. CONCLUSIONS

As motivation for this work, we proposed that integration of magnetic tweezers (MT) with deformation tracking microscopy and traction force microscopy (DTM/TFM) could be used to explore unanswered questions in the field keratinocyte mechanobiology. Towards this end, we have developed a methodology for integrating these techniques, as demonstrated via two distinct model bead-on-gel experiments referred to as force-control (FC) mode and displacement-control (DC) mode MT-DTM/TFM experiments. In the model FC mode bead-on-gel experiment, we showed how the integrated methodology could be used to characterize the rheological behavior of a type I collagen gel substrate. Specifically, we presented two quantitative approaches based on TFM analysis, namely, a force-based method for estimating the apparent elastic modulus of the gel, and an energy-based technique for quantifying an absolute upper bound for the elastic modulus. Because FC mode experiments are inherently dynamic, an *ad hoc* criterion was established to validate the existence of elastostatic conditions inherent to the TFM analysis used as a basis for these calculations. In a model DC mode bead-on-gel experiment, we substantiated the idea that the integrated methodology can be implemented in experiments requiring either large substrate deformations (>5 μm) and/or large forces (>5 nN). Important considerations for DC mode experiments were identified, including a classification of bead-tip clamp failure mechanisms and estimation of the maximum bead-tip magnetic clamping force. Lastly, in a proof-of-concept FC mode bead-on-cell experiment, we applied a defined MT force to a fibronectin-coated superparamagnetic bead attached to a keratinocyte present at the periphery of a multicellular sheet reconstituted under low calcium conditions. Both the measured substrate displacement fields and the calculated incremental traction stress fields observed for this bead-on-cell experiment suggest that a force applied to the apical surface of the keratinocyte is directly transmitted through the cell to its underlying substrate. Collectively, the integrated MT-DTM/TFM experiments and rigorous approaches to data reduction presented in this work will provide a necessary basis for more widespread application of the integrated methodology in future explorations of keratinocyte mechanobiology in the context of human blistering skin disease.

## VI. SUPPLEMENTARY MATERIAL

A comprehensive list of abbreviations and symbols used in this work can be found in the **Supplementary Material**, as are detailed narratives of the experimental procedures and methods of data reduction developed for FC mode and DC mode MT-DTM/TFM experiments. Additional materials found in the **Supplementary Material** include: sample magnetic actuation waveform data for FC mode experiments; analysis of our FC mode null force control and DC mode null displacement control experiments; data utilized for L2 regularization parameter determination in our FC and DC mode experiments; and a tiled and stitched phase contrast image showing the reconstituted multicellular keratinocyte sheet used for the FC mode bead-on-cell experiment detailed in Sec. IV.B.

## Supporting information

Vid 1 FC DIC

Vid FC EPI

VID 3 FC displacement

Vid 4 FC traction

Vid 5 FC strain energy density

Vid 6 FC null DIC

Vid 7 FC null EPI

Vid 8 FC null displacement

Vid 9 FC null traction

Vid 10 FC null strain energy density

Vid 11 DC DIC

VID 12 DC DIC zoom

Vid 13 DC EPI

Vid 14 DC displacement

Vid 15 DC traction

Vid 16 DC strain energy density

Vid 17 DC null DIC

Vid 18 DC null DIC zoom

Vid 19 DC null EPI

Vid 20 DC null displacement

Vid 21 DC null traction

Vid 22 DC null strain energy density

Vid 23 cell DIC

Vid 24 cell EPI

Vid 25 cell displacement

Vid 26 cell traction

Vid 27 cell strain energy density

## VII. ACKNOWLEDGEMENTS

The authors acknowledge Dan Witte and George DeBeck (Nikon Instruments, Inc.) for their expert technical assistance with the setup and operation of our microscope. J.C.S. thanks the Dermatology Foundation for its generous physician scientist career development award. E.A.S. recognizes the National Science Foundation (CAREER 1452728).

## VIII. AUTHOR’S CONTRIBUTIONS

All authors contributed equally to this work.

## IX. DATA AVAILABILITY

The data that supports the findings of this study are available within the article and its supplementary material.

## Notes

### Competing Interest Statement

The authors have declared no competing interest.

## REFERENCES

1. J.A. McGrath and J. Uitto, in Rook’s Textb. Dermatology, edited by C.E.M. Griffiths, J. Barker, T. Bleiker, R. Chalmers, and D. Creamer, 9th ed. (John Wiley & Sons, Ltd, Chichester, UK, 2016), pp. 1–48.

2. J.C. Selby, in Mechanobiol. Cell-Cell Cell-Matrix Interact., edited by A. Wagoner Johnson and B.A.C. Harley (Springer US, Boston, MA, 2011), pp. 169–210.

3. D. Tsuruta, T. Hashimoto, K.J. Hamill, and J.C.R. Jones, J. Dermatol. Sci. 62, 1 (2011).

4. M. Hatzfeld, R. Keil, and T.M. Magin, Cold Spring Harb. Perspect. Biol. 9, a029157 (2017).

5. I. Turcan and M.F. Jonkman, in Autoimmune Bullous Dis. Text Rev., edited by M.F. Jonkman (Springer International Publishing, Cham, 2016), pp. 113–117.

6. C.M. Hammers and J.R. Stanley, Annu. Rev. Pathol. Mech. Dis. 11, 175 (2016).

7. R. Ogawa and C.-K. Hsu, J. Cell. Mol. Med. 17, 817 (2013).

8. K. Karlmark and R. Eming, in Blistering Dis. Clin. Featur. Pathog. Treat., edited by D.F. Murrell (Springe-Verlagr Berlin Heidelberg, Berlin, Heidelberg, 2015), pp. 21–33.

9. C.-K. Hsu, H.-H. Lin, H.I.-C. Harn, M.W. Hughes, M.-J. Tang, and C.-C. Yang, J. Dermatol. Sci. 90, 232 (2018).

10. F. Ziemann, J. Rädler, and E. Sackmann, Biophys. J. 66, 2210 (1994).

11. A.R. Bausch, F. Ziemann, A.A. Boulbitch, K. Jacobson, and E. Sackmann, Biophys. J. 75, 2038 (1998).

12. J.K. Fisher, J.R. Cummings, K. V. Desai, L. Vicci, B. Wilde, K. Keller, C. Weigle, G. Bishop, R.M. Taylor, C.W. Davis, R.C. Boucher, E.T. O’Brien, and R. Superfine, Rev. Sci. Instrum. 76, 053711 (2005).

13. M. Tanase, N. Biais, and M. Sheetz, in Cell Mech., edited by Y. Wang and D.E. Discher (Elsevier Inc., 2007), pp. 473–493.

14. C. Aermes, A. Hayn, T. Fischer, and C.T. Mierke, Sci. Rep. 10, 13453 (2020).

15. H. Zarkoob, S. Bodduluri, S. V. Ponnaluri, J.C. Selby, and E.A. Sander, Cell. Mol. Bioeng. 8, 32 (2015).

16. H. Zarkoob, S. Chinnathambi, S.A. Halberg, J.C. Selby, T.M. Magin, and E.A. Sander, Cell. Mol. Bioeng. 11, 163 (2018).

17. U.S. Schwarz and J.R.D. Soiné, Biochim. Biophys. Acta - Mol. Cell Res. 1853, 3095 (2015).

18. J.H.-C. Wang and J.-S. Lin, Biomech. Model. Mechanobiol. 6, 361 (2007).

19. C.M. Kraning-Rush, S.P. Carey, J.P. Califano, and C.A. Reinhart-King, in Methods Cell Biol., edited by A.R. Asthagiri and A.P. Arkin (Elsevier Inc., 2012), pp. 139–178.

20. B. Sabass, M.L. Gardel, C.M. Waterman, and U.S. Schwarz, Biophys. J. 94, 207 (2008).

21. L. Selvaggi, L. Pasakarnis, D. Brunner, and C.M. Aegerter, Rev. Sci. Instrum. 89, 045106 (2018).

22. X. Trepat, M.R. Wasserman, T.E. Angelini, E. Millet, D.A. Weitz, J.P. Butler, and J.J. Fredberg, Nat. Phys. 5, 426 (2009).

23. M.R. Ng, A. Besser, J.S. Brugge, and G. Danuser, Elife 3, e03282 (2014).

24. A.F. Mertz, Y.L. Che, S. Banerjee, J.M. Goldstein, K.A. Rosowski, S.F. Revilla, C.M. Niessen, M.C. Marchetti, E.R. Dufresne, and V. Horsley, Proc. Natl. Acad. Sci. 110, 842 (2013).

25. Y. Yang, J. Lin, R. Meschewski, E. Watson, and M. Valentine, Biotechniques 51, 29 (2011).

26. J. Bush and V. Maruthamuthu, AIP Adv. 9, 035221 (2019).

27. W.I. Moghram, A. Kruger, E.A. Sander, and J.C. Selby, Rev. Sci. Instrum. Submitted (2020).

28. I. Muhamed, F. Chowdhury, and V. Maruthamuthu, Bioengineering 4, 12 (2017).

29. K. Haubert, T. Drier, and D. Beebe, Lab Chip 6, 1548 (2006).

30. S. Ettori, J.-C. Peraud, and J. Barton, U.S. Patent No. 4,824,458 (1989).

31. T. Yeung, P.C. Georges, L.A. Flanagan, B. Marg, M. Ortiz, M. Funaki, N. Zahir, W. Ming, V. Weaver, and P.A. Janmey, Cell Motil. Cytoskeleton 60, 24 (2005).

32. G.B. Zamansky, U. Nguyen, and U. Lih-Nan Chou, J. Invest. Dermatol. 97, 985 (1991).

33. E.J. O’Keefe, R.A. Briggaman, and B. Herman, J. Cell Biol. 105, 807 (1987).

34. F.M. Watt, D.L. Mattey, and D.R. Garrod, J. Cell Biol. 99, 2211 (1984).

35. J.C.R. Jones and R.D. Goldman, J. Cell Biol. 101, 506 (1985).

36. V. Vasioukhin, C. Bauer, M. Yin, and E. Fuchs, Cell 100, 209 (2000).

37. L.I. Gold and E. Pearlstein, Biochem. J. 186, 551 (1980).

38. V. Auernheimer, L.A. Lautscham, M. Leidenberger, O. Friedrich, B. Kappes, B. Fabry, and W.H. Goldmann, J. Cell Sci. 128, 3435 (2015).

39. R.W. Style, R. Boltyanskiy, G.K. German, C. Hyland, C.W. MacMinn, A.F. Mertz, L.A. Wilen, Y. Xu, and E.R. Dufresne, Soft Matter 10, 4047 (2014).

40. S.J. Han, Y. Oak, A. Groisman, and G. Danuser, Nat. Methods 12, 653 (2015).

41. M.K. Cheezum, W.F. Walker, and W.H. Guilford, Biophys. J. 81, 2378 (2001).

42. L. Gui and S.T. Wereley, Exp. Fluids 32, 506 (2002).

43. J. Westerweel and F. Scarano, Exp. Fluids 39, 1096 (2005).

44. J. Duncan, D. Dabiri, J. Hove, and M. Gharib, Meas. Sci. Technol. 21, 057002 (2010).

45. D.T. Sandwell, Geophys. Res. Lett. 14, 139 (1987).

46. J.A. Mulligan, F. Bordeleau, C.A. Reinhart-King, and S.G. Adie, in Biomech. Oncol., edited by C. Dong, N. Zahir, and K. Konstantopoulos (Springer International Publishing, Cham, 2018), pp. 319–349.

47. X. Tang, A. Tofangchi, S. V Anand, and T.A. Saif, PLOS Comput. Biol. 10, e1003631 (2014).

48. S. V. Plotnikov, B. Sabass, U.S. Schwarz, and C.M. Waterman, in Methods Cell Biol., edited by J.C. Waters and T. Wittman (Elsevier Inc., 2014), pp. 367–394.

49. Y. Huang, C. Schell, T.B. Huber, A.N. Şimşek, N. Hersch, R. Merkel, G. Gompper, and B. Sabass, Sci. Rep. 9, 539 (2019).

50. B. Lee, X. Zhou, K. Riching, K.W. Eliceiri, P.J. Keely, S.A. Guelcher, A.M. Weaver, and Y. Jiang, PLoS One 9, e111896 (2014).

51. K. Slater, J. Patridge, and H. Nandivada, Tuning the Elastic Moduli of Corning® Matrigel® and Collagen I 3D Matrices by Varying the Protein Concentration (Corning Inc., Bedford, MA, 2017).

52. Y. Yang, L.M. Leone, and L.J. Kaufman, Biophys. J. 97, 2051 (2009).

53. S.M. Mijailovich, M. Kojic, M. Zivkovic, B. Fabry, and J.J. Fredberg, J. Appl. Physiol. 93, 1429 (2002).

54. A.R. Bausch, W. Möller, and E. Sackmann, Biophys. J. 76, 573 (1999).

55. E.A. Sander and V.H. Barocas, in Collagen Struct. Mech., edited by P. Fratzl (Springer US, Boston, MA, 2008), pp. 475–504.

56. J. Ferruzzi, Y. Zhang, D. Roblyer, and M.H. Zaman, in Multi-Scale Extracell. Matrix Mech. Mechanobiol., edited by Y. Zhang (Springer International Publishing, Cham, 2020), pp. 343–387.

57. L.D. Landau and E.M. Lifshitz, Theory of Elasticity, 3rd ed. (Butterworth-Heinemann, Oxford, United Kingdom, 1986).

58. Z. Cenev, H. Zhang, V. Sariola, A. Rahikkala, D. Liu, H.A. Santos, and Q. Zhou, Adv. Mater. Technol. 3, 1700177 (2018).

59. M. Cantini, C. González-García, V. Llopis-Hernández, and M. Salmerón-Sánchez, in ACS Symp. Ser. (American Chemical Society, 2012), pp. 471–496.

60. J.P. Butler, I.M. Tolić-Nørrelykke, B. Fabry, and J.J. Fredberg, Am. J. Physiol. Physiol. 282, C595 (2002).

61. A.F. Mertz, S. Banerjee, Y. Che, G.K. German, Y. Xu, C. Hyland, M.C. Marchetti, V. Horsley, and E.R. Dufresne, Phys. Rev. Lett. 108, 198101 (2012).

62. B. Fabry, A.H.H. Klemm, S. Kienle, T.E.E. Schäffer, and W.H.H. Goldmann, Biophys. J. 101, 2131 (2011).

