## Supplementary Material for "Integration of Magnetic Tweezers and Traction Force Microscopy for the Exploration of Matrix Rheology and Keratinocyte Mechanobiology: Model Force- and Displacement-Controlled Experiments"

### LIST OF ABBREVIATIONS AND SYMBOLS

| | |
|---|---|
| $B_{\text{blunt}}(t)$ | *measured* magnetic flux density emanating from the blunt end of the needle core of the MT device |
| $B_{\text{blunt}}^{\text{OFF}}$ | setpoint in magnetic flux density emanating from the blunt end of the MT device during OFF intervals in the magnetic actuation waveform sequence |
| $B_{\text{blunt}}^{\text{ON}}$ | setpoint in magnetic flux density emanating from the blunt end of the MT device during ON intervals in the magnetic actuation waveform sequence |
| $B_{\text{blunt}}^{\text{ON}*}$ | *calibrated* control setpoint in magnetic flux density of the MT device during ON intervals in the magnetic actuation waveform sequence |
| $B_{\text{blunt}}^{\text{perm}}$ | *calibrated* magnetic flux density of the MT device during OFF intervals in any magnetic actuation waveform sequence that is needed to nullify the *permanent* magnetic field present at the needle tip following degaussing of the needle core |
| $B_{\text{blunt}}^{\text{rem}}$ | additional *calibrated* corrective magnetic flux density setpoint of the MT device during OFF intervals in any magnetic actuation waveform sequence that is needed to nullify the remnant magnetic field present at the needle tip following magnetization of the core |
| $c$ | collagen concentration |
| $\boldsymbol{\delta}(t)$ | vector defining the position of the centroid of the superparamagnetic bead with respect to an origin fixed to the needle tip of the MT device |
| $\delta(t)$ | scalar Euclidean separation distance between the centroid of the superparamagnetic bead and the needle tip of the MT device |
| $\boldsymbol{\Delta}_{\text{max}}(t)$ | displacement vector within the $x^*y^*$-plane of the microsphere embedded within the collagen substrate that experiences the largest overall displacement in response to $\boldsymbol{F}_{\text{MT}}(t)$; a quantity equal to $\boldsymbol{u}(x^*_{\text{max}}, y^*_{\text{max}}, t)$ |
| $\Delta_{\text{max}}(t)$ | scalar magnitude of $\boldsymbol{\Delta}_{\text{max}}(t)$ |
| $\boldsymbol{\Delta}_{\text{MT}}$ | displacement vector of the superparamagnetic bead in the $xy$-plane (surface) of the collagen substrate defined with respect to the initial $(x, y, z)$ spatial coordinates of the bead |
| $\Delta_{\text{MT}}$ | scalar magnitude of $\boldsymbol{\Delta}_{\text{MT}}$ |

| | |
|---|---|
| $E(c)$ | collagen concentration-dependent Young's elastic modulus |
| $\boldsymbol{F}(\boldsymbol{t})$ | total traction force vector present in the $x^{*}y^{*}$-surface plane of the collagen substrate at time, $t$ |
| $F(t)$ | scalar magnitude of the total traction force vector |
| $\boldsymbol{F}_{x^{*}}(t)$ | vectoral component of $\boldsymbol{F}(t)$ in the $x^{*}$-direction |
| $\boldsymbol{F}_{y^{*}}(t)$ | vectoral component of $\boldsymbol{F}(t)$ in the $y^{*}$-direction |
| $\boldsymbol{F}^{*}_{\mathrm{MT}}\left(\delta(t), B^{\mathrm{ON}*}_{\mathrm{blunt}}\right)$ | magnetic actuation force vector generated by the MT device as a calibrated function of $\delta(t)$ and $B^{\mathrm{ON}*}_{\mathrm{blunt}}$ |
| $\boldsymbol{F}_{\mathrm{MT}}(t)$ | component of the magnetic actuation force vector in the $xy$-plane of the collagen substrate |
| $F_{\mathrm{MT}}(t)$ | scalar magnitude of $\boldsymbol{F}_{\mathrm{MT}}(t)$ |
| $G(c)$ | collagen concentration-dependent shear modulus |
| $i$ | index used to define individual microspheres present on the surface of a collagen substrate |
| $\lambda 2(n)$ | raw optimal L2 regularization parameter at a generic frame number, $n$ |
| $\lambda 2(t)$ | raw optimal L2 regularization parameter at time, $t$ |
| $\lambda 2_{\mathrm{ON}}$ | global L2 regularization parameter used to compute solutions for $\boldsymbol{T}(x^{*}, y^{*}, t)$ associated with all ON segments of an FC or DC MT-DTM/TFM experiment |
| $\lambda 2_{\mathrm{OFF}}$ | global L2 regularization parameter used to compute solutions for $\boldsymbol{T}(x^{*}, y^{*}, t)$ associated with all OFF segments of an FC or DC MT-DTM/TFM experiment |
| $\nu$ | Poisson's ratio |
| μ-sphere $[i]$ | microsphere identified by the index, $i$, embedded within the surface of the collagen substrate |
| $n$ | generic frame number |
| $n_{\mathrm{OFF}}$ | total number of imaging frames binned as OFF data |
| $n_{\mathrm{ON}}$ | total number of imaging frames binned as ON data |

| | |
|---|---|
| $n_{\mathrm{TOTAL}}$ | total number of imaging frames |
| $n_{\mathrm{TRANS}}$ | total number of imaging frames that captured transients between ON and OFF segments with the overall magnetic actuation waveform of an FC MT-DTM/TFM experiment |
| $\rho(x^*, y^*, t)$ | strain energy density (scalar) field in the $x^*y^*$-surface plane of the collagen substrate at time, $t$ |
| σ | best-fit Rayleigh distribution parameter for a given single epifluorescence imaging frame in a null displacement control experiment |
| $\bar{\sigma}$ | mean best-fit Rayleigh distribution parameter over all epifluorescence imaging frames in a null displacement control experiment |
| $t$ | time |
| $t_0$ | time $t$ = 0 s |
| $t_{\mathrm{OFF}}$ | time marking the last imaging frame captured prior to the start of the OFF segment of a specified cycle within an overall magnetic actuation waveform sequence of an FC mode experiment |
| $t_{\mathrm{ON}}$ | time marking the first imaging frame captured following the start of the ON segment of a specified cycle within an overall magnetic actuation waveform sequence of an FC mode experiment |
| $t_{\mathrm{EPI}}$ | raw time point within the epifluorescence image sequence of a DC mode experiment marking the first statistically significant microsphere displacement measurement |
| $t_{\mathrm{DIC}}$ | raw time point within the DIC image sequence of a DC mode experiment marking the first statistically significant superparamagnetic bead displacement measurement |
| $t_{\mathrm{r}}$ | variable used to denote raw times of the epifluorescence data set within a DC MT-DTM/TFM experiment |
| $\boldsymbol{T}(x^*, y^*, t)$ | stress traction vector field in the $x^*y^*$-plane of the collagen substrate at time, $t$ |
| $\boldsymbol{T}_{x^*}(x^*, y^*, t)$ | vectoral component of $\boldsymbol{T}(x^*, y^*, t)$ in the $x^*$-direction |
| $\boldsymbol{T}_{y^*}(x^*, y^*, t)$ | vectoral component of $\boldsymbol{T}(x^*, y^*, t)$ in the $y^*$-direction |

| | |
|---|---|
| $\boldsymbol{u}(x^*, y^*, t)$ | interpolated two-dimensional continuum representation of the collagen substrate displacement (vector) field in the $x^*y^*$-plane at time, $t$ |
| $\boldsymbol{u}_{x^*}(x^*, y^*, t)$ | vectoral component of $\boldsymbol{u}(x^*, y^*, t)$ in the $x^*$-direction |
| $\boldsymbol{u}_{y^*}(x^*, y^*, t)$ | vectoral component of $\boldsymbol{u}(x^*, y^*, t)$ in the $y^*$-direction |
| $\boldsymbol{u}(\mu\text{-sphere } [i], t)$ | displacement vector of μ-sphere $[i]$ at time, $t$ |
| $\boldsymbol{u}_{x^*}(\mu\text{-sphere } [i], t)$ | vectoral component of the displacement vector of μ-sphere $[i]$ at time, $t$, in the $x^*$-direction |
| $\boldsymbol{u}_{y^*}(\mu\text{-sphere } [i], t)$ | vectoral component of the displacement vector of μ-sphere $[i]$ at time, $t$, in the $y^*$-direction |
| $\frac{\partial^2}{\partial t^2}\boldsymbol{u}_{x^*}(\mu\text{-sphere } [i],\ \ t)$ | vectoral component of the acceleration vector of μ-sphere $[i]$ at time, $t$, in the $x^*$-direction |
| $\frac{\partial^2}{\partial t^2}\boldsymbol{u}_{y^*}(\mu\text{-sphere } [i],\ \ t)$ | vectoral component of the acceleration vector of μ-sphere $[i]$ at time, $t$, in the $y^*$-direction |
| $\lvert\boldsymbol{u}_{6\overline{\sigma}}(\mu\text{-sphere } [i])\rvert$ | $6\overline{\sigma}$ upper limit of microsphere displacements as measured in a null displacement control experiment |
| $\boldsymbol{u}_{x^*}(x^*_{\max}, y^*_{\max}, t)$ | vectoral component in the $x^*$-direction at time, $t$, of the displacement vector of the microsphere embedded within the collagen substrate that was subject to the overall largest displacement during an FC or DC mode experiment |
| $\boldsymbol{u}_{y^*}(x^*_{\max}, y^*_{\max}, t)$ | vectoral component in the $y^*$-direction at time, $t$, of the displacement vector of the microsphere embedded within the collagen substrate that was subject to the overall largest displacement during an FC or DC mode experiment |
| $\frac{\partial^2}{\partial t^2}\boldsymbol{u}_{x^*}(x^*_{\max}, y^*_{\max}, t)$ | vectoral component in the $x^*$-direction at time, $t$, of the acceleration vector of the microsphere embedded within the collagen substrate that was subject to the overall largest displacement during an FC or DC mode experiment |
| $\frac{\partial^2}{\partial t^2}\boldsymbol{u}_{y^*}(x^*_{\max}, y^*_{\max}, t)$ | vectoral component in the $y^*$-direction at time, $t$, of the acceleration vector of the microsphere embedded within the collagen substrate that was subject to the overall largest displacement during an FC or DC mode experiment |
| $U(t)$ | total strain energy associated with deformation of the collagen substrate at time, $t$ |

| | |
|---|---|
| $W_{\mathrm{MT}}(t)$ | work done by the MT device in moving the superparamagnetic bead during the ON segment of a specified cycle within the overall magnetic actuation waveform sequence of an FC mode experiment |
| $x$ | orthogonal reference coordinate used to locate the position of the superparamagnetic bead in the horizontal imaging plane (collagen substrate surface) with respect to the needle tip of the MT device |
| $x^*$ | one of two orthogonal reference coordinates used to define a horizontal positional grid of material points at the surface of the collagen substrate |
| $x^*_{\mathrm{max}}$ | $x^*$coordinate at time $t = 0$ of the microsphere embedded in the collagen substrate that experiences the largest overall displacement during an FC or DC mode experiment |
| $y$ | orthogonal reference coordinate used to locate the position of the superparamagnetic bead in the horizontal imaging plane (collagen substrate surface) with respect to the needle tip of the MT device |
| $y^*$ | one of two orthogonal reference coordinates used to define a horizontal positional grid of material points as a continuum representation of the collagen substrate |
| $y^*_{\mathrm{max}}$ | $y^*$coordinate at time $t = 0$ of the microsphere embedded in the collagen substrate that experiences the largest overall displacement during an FC or DC mode experiment |
| $z$ | orthogonal reference coordinate used to locate the vertical position of the superparamagnetic bead with respect to the needle tip of the MT device |

### SI. MT-DTM/TFM MODES OF OPERATION

#### A. Force-control (FC) mode

##### *1. Experimental procedure*

For nominal FC experiments conducted on both collagen gels and keratinocytes, the needle core of the MT device was first fixed in the standard configuration at an inclination angle of ~25° with respect to the microscope stage plane (the horizon). Under live DIC imaging (~14 to 40 fps) at 30X magnification, the needle tip was positioned in the vicinity of the superparamagnetic bead to be actuated such that, at time $t$ = 0 (or $t_0$), the bead position with respect to the needle tip was 10 µm to 20 µm horizontally away from the tip, $x(t_0)$; 10 µm to 15 µm vertically below the tip, $z(t_0)$; and at the same lateral $y$-position, $y(t_0)$ = 0 µm (see **Fig. 1(c)** of main article). Here, the orthogonal $(x, y, z)$-coordinate system used to define the position of the superparamagnetic bead has an origin fixed to the needle tip of the MT device. Initial positioning of the needle tip and superparamagnetic bead of interest required multiple sequential adjustments to both the motorized stage of the microscope and the manual micromanipulator that translates the needle tip. Once initialized, the needle tip was assumed to remain spatially fixed for the duration of the experiment.

Following configuration of the MT device, the microscope was set to image in DIC mode, focusing on the superparamagnetic bead at 30X magnification. The MT device was triggered to deploy a user-selected ON/OFF magnetic actuation force waveform sequence, simultaneously initiating a fast time-lapse image acquisition at 40 fps (2048x2044 pixels per frame) in Nikon Elements. ON time intervals ranged from 1 s to 5 s, and OFF times ranged from 1 s to 7 s. An initial 1 to 2 s OFF interval preceded the start of the overall ON/OFF magnetic waveform actuation sequence. Due to the non-deterministic operating system running the Nikon Elements software, fast time-lapse image acquisition started at a random time point within this initial 1 to 2 s OFF interval for each experiment. The superparamagnetic bead remained in focus throughout the duration of the magnetic actuation waveform sequence. As such, bead displacements in the $z$-direction were assumed to be negligible.

The nominal ON magnetic actuation force was determined by the *calibrated* magnetic flux density control setpoint configured for the blunt end of the needle ($B_{\mathrm{blunt}}^{\mathrm{ON}*}$) and the calculated Euclidean distance between the superparamagnetic bead and the needle tip, $\delta(t)$, as discussed in SI.A.2 (see also **Fig. 1(c)** of the main article). Based on a detailed force calibration procedure[1], nominal applied loads ranging from 0 nN to 25 nN were possible for $B_{\mathrm{blunt}}^{\mathrm{ON}*}$ setpoints ranging from 0 to 175 Gs. Magnetic flux density setpoints configured for the OFF segments of the actuation sequence, denoted here as $B_{\mathrm{blunt}}^{\mathrm{OFF}}$, employed empiric calibrated corrective schemes to null both the permanent ($B_{\mathrm{blunt}}^{\mathrm{perm}}$) and remnant magnetization ($B_{\mathrm{blunt}}^{\mathrm{rem}}$) of the needle core. In this manner, our MT device generates a null magnetic actuation force during designated

OFF time intervals ($\boldsymbol{F}_{\mathrm{MT}}^{*}$ = 0 nN) of any given actuation cycle[1]. As opposed to perfectly square ON/OFF waveforms, magnetic actuation in our setup included short stepwise ramps in magnetic flux density between ON and OFF states. Specifically, the MT device stepped up (or down) to the desired magnetic flux density control setpoints by passing through intermediate *calibrated* values of $B_{\mathrm{blunt}}^{\mathrm{ON}*}$ (25 Gs, 50 Gs, 75 Gs, 100 Gs, 125 Gs, and 150 Gs), maintaining each of these intermediate flux density setpoints for ~70 ms before proceeding to the next setpoint (up or down). Here, $B_{\mathrm{blunt}}^{\mathrm{ON}*}$ represents the *calibrated* setpoint, defined here as the difference between $B_{\mathrm{blunt}}^{\mathrm{ON}}$ and $B_{\mathrm{blunt}}^{\mathrm{perm}}$[1]. As a consequence of this magnetic actuation scheme, there exists a ~500 ms transient interval in magnetic actuation force, $\boldsymbol{F}_{\mathrm{MT}}^{*}$, that separates ON and OFF states of a given actuation cycle. Use of stepwise ON/OFF ramps during magnetic actuation was intentional, because it facilitated more reliable tracking of the kinematics of the fluorescent microspheres embedded in our collagen substrates. For the FC mode bead-on-gel experiment presented in **Fig. 2** of the main article, the magnetic actuation waveform sequence consisted of 3 identical 3 s/7 s ON/OFF cycles, each with $B_{\mathrm{blunt}}^{\mathrm{ON}*}$ setpoints of 175 Gs. Measured values of magnetic flux density emanating from the blunt end of the MT device, $B_{\mathrm{blunt}}(t)$, were recorded throughout the entire actuation sequence. Illustrative data from the first cycle of the FC mode bead-on-gel experiment presented in **Fig. 2** of the main article is shown in **Fig. S1**.

After capturing the response of the superparamagnetic bead to the prescribed magnetic actuation waveform sequence under DIC imaging, the microscope was set to image with Texas Red epifluorescence, again at 30X magnification. The focus was adjusted to optimize imaging of the fluorescent microspheres embedded within the surface of the collagen substrate. Because the microscope focus was determined by vertical translation of the objective and not the microscope stage, the $z$-coordinate defining the position of the superparamagnetic bead with respect to the needle tip did not change during this re-focusing procedure. The horizontal and lateral positioning of the microscope stage also remained fixed during this re-focusing procedure, however, the $(x, y)$-coordinates of the superparamagnetic bead were subject to small changes depending on the mechanical response of bead to the *first* magnetic actuation waveform sequence. From a biophysical standpoint, the response of the superparamagnetic bead was directly related to the constitutive mechanical properties of the cell or substrate to which the bead was attached. After a time delay of ~60 s required to configure the microscope for optimal epifluorescence imaging, the MT device was triggered to deploy a *second* OFF/ON magnetic actuation force waveform sequence, identical to the first, concurrently with a *second* fast time-lapse image acquisition at 40 fps (2048x2044 pixels per frame). Thus, for every magnetic actuation force waveform sequence tested, two sequential yet coupled data sets were collected: a set of DIC images used to track the displacement of the superparamagnetic bead, and a set of epifluorescence images used to track localized displacements of the collagen substrate.

*2. Data reduction*

To analyze data acquired during a FC experiment, TTL pulses marking sCMOS camera exposures were used to temporally correlate the acquired epifluorescence or DIC image sets with the measured magnetic flux density waveform data. For DIC image sets, the substrate drift-corrected positions of the centroid of the superparamagnetic bead were tracked as described in Sec. III.E of the main article. The central 1024 x 1022 pixels of each image were used as the global ROI for data reduction. With this positional data, we computed the displacement vector of the centroid of the superparamagnetic bead relative to its initial position (at $t = 0$) in the $xy$-plane of the collagen substrate as a function of time, denoted here as $\boldsymbol{\Delta}_{\mathrm{MT}}(t)$, and its corresponding scalar magnitude, $\Delta_{\mathrm{MT}}(t)$, (see **Fig. 1(c)** of main article). Knowing the initial spatial configuration of the needle tip with respect to the centroid of the superparamagnetic bead and assuming that the needle tip position remains fixed and bead displacements in the $z$-direction are negligible throughout the duration of the experiment ($z(t) = z(t_0)$), we calculated the scalar Euclidean bead-tip separation distance as a function of time, $\delta(t) = \sqrt{x(t)^2 + y(t)^2 + z(t_0)^2}$, where $x(t)$ and $y(t)$ represent the positional coordinates of the superparamagnetic bead in the $xy$-plane of the substrate. The magnetic actuation force vector generated by the MT device, $\boldsymbol{F}^{*}_{\mathrm{MT}}(\delta(t), B_{\mathrm{blunt}}(t))$, was calculated using $\delta(t)$ and the measured $B_{\mathrm{blunt}}(t)$ values recorded for the prescribed magnetic actuation waveform sequence. Specifically, for all values of $t$ where $B_{\mathrm{blunt}}(t) > 0$ Gs, we first computed the magnitude of $\boldsymbol{F}^{*}_{\mathrm{MT}}(t)$ by applying the corresponding value of $\delta(t)$ to the calibration power law associated with the largest of the six discrete nominal calibrated values of $B^{\mathrm{ON}^*}_{\mathrm{blunt}}$ for which $B_{\mathrm{blunt}}(t) > B^{\mathrm{ON}^*}_{\mathrm{blunt}}$ [1]. In this manner, $\boldsymbol{F}^{*}_{\mathrm{MT}}(t)$ represents the applied MT force as a function of time with inaccuracies limited to temporal transients in $B_{\mathrm{blunt}}(t)$ that occur between calibrated setpoints in $B^{\mathrm{ON}^*}_{\mathrm{blunt}}$. By design, $\boldsymbol{F}^{*}_{\mathrm{MT}}(t)$ always points towards the needle tip (see **Fig. 1(c)** of main article). For integrated MT-DTM/TFM experiments, we restricted our analysis to the vectoral component of $\boldsymbol{F}^{*}_{\mathrm{MT}}(t)$ that resides within the $xy$-plane (and thus $x^*y^*$-plane) of the collagen substrate, a quantity defined here as $\boldsymbol{F}_{\mathrm{MT}}(t)$, with a magnitude defined as $F_{\mathrm{MT}}(t) = |\boldsymbol{F}^{*}_{\mathrm{MT}}(t)|\left(\sqrt{x(t)^2 + y(t)^2}/\delta(t)\right)$. Based on calibrations of our MT device[1], we estimate maximum relative uncertainties in $F_{\mathrm{MT}}(t)$ of ±25% for superparamagnetic beads configured with initial $\left(x(t_0), y(t_0), z(t_0)\right)$-coordinates within the ranges previously designated, subject to the condition that the beads developed $x$- and $y$-displacements of at most ±5.5 μm following magnetic actuation. Note that with our experimental setup, only the *initial* force applied to the superparamagnetic bead can be controlled by the experimenter. Our current setup is not configured for the real-time feedback adjustments in $\delta(t)$ that would be required to maintain a constant force as the bead begins to move towards the needle tip in response to the applied magnetic field[2].

For Texas Red epifluorescence image sets in an FC experiment, microsphere displacements were tracked and used to generate spatial outlier-corrected, temporally filtered, and drift-corrected interpolated displacement fields, $\boldsymbol{u}(x^*, y^*, t)$, using the procedure outlined in Sec. III.E of the main article. By default, $\boldsymbol{u}(x^*, y^*, 0) = 0$. The central 1024 x 1022 pixels of each image were used as the global ROI for data reduction. Videos superimposing the original epifluorescence images overlaid with arrows representing scaled individual microsphere displacement vectors were generated, as well as heatmaps of $\boldsymbol{u}(x^*, y^*, t)$. For FC experiments, the initial (at $t$ = 0 s = $t_0$) spatial location of the microsphere on the collagen substrate subject to the largest displacement during a given experiment was identified as $(x^*_{\text{max}}, y^*_{\text{max}})$. The measured (non-interpolated) microsphere displacement vector as a function of time at this location was defined as $\boldsymbol{\Delta}_{\text{max}}(t) = \boldsymbol{u}(x^*_{\text{max}}, y^*_{\text{max}}, t)$, with a scalar magnitude denoted by $\Delta_{\text{max}}(t)$. When $\boldsymbol{u}(x^*, y^*, t)$ along the $x^*y^*$-boundary of the global ROI was roughly zero, the stress traction vector field, $\boldsymbol{T}(x^*, y^*, t)$, was calculated for all time $t$, i.e., for each frame in the overall fast time-lapse sequence of acquired images. Once $\boldsymbol{T}(x^*, y^*, t)$ were known, the total traction force, $\boldsymbol{F}(t)$; the magnitude of the total traction force, $F(t)$; the strain energy density field, $\rho(x^*, y^*, t)$; and the total strain energy, $U(t)$; were calculated using the procedures outlined in Sec. III.F of the main article.

In solving for $\boldsymbol{T}(x^*, y^*, t)$, we employed an algorithmic approach for selection of the L2 regularization parameter, $\lambda 2$, described as follows. First, the raw optimal regularization parameter at each instance of time, $\lambda 2(t)$, was determined using a Bayesian approach, given its intrinsic computational efficiency compared to other methodologies[3]. Reduced (gridded) displacement data was first spatially filtered with a Weiner filter (3 x 3 pixel) and then padded by a margin of zeros. Displacements within small square ROIs localized to the four corners of the global ROI at each discrete time instance, $t$, (each square ~10% of the global ROI) were used as a fiducial sample of displacement noise. We binned each $\lambda 2(t)$ data point as occurring within an ON, OFF, or transient (TRANS) segment of the overall magnetic actuation waveform, reassigning the independent variable, $t$, to a generic frame number, $n$, or $\lambda 2(t) \rightarrow \lambda 2(n)$. Transient imaging frames occurring between ON and OFF segments were omitted from the analysis. We then defined two global L2 regularization parameters based on the mean of the logarithms of $\lambda 2(n)$ for each binned data set, or

$$\lambda 2_{\text{ON}} = 10^{\left(\frac{\sum_1^{n_{\text{ON}}} \log_{10}(\lambda 2(n))}{n_{\text{ON}}}\right)} \quad \text{and} \quad \lambda 2_{\text{OFF}} = 10^{\left(\frac{\sum_1^{n_{\text{OFF}}} \log_{10}(\lambda 2(n))}{n_{\text{OFF}}}\right)}, \qquad \text{(S1)}$$

where $n_{\text{ON}}$ represents the total number of analyzed imaging frames binned as ON data and $n_{\text{OFF}}$ represents the total number of frames binned as OFF data. By rule, $n_{\text{ON}} + n_{\text{OFF}} + n_{\text{TRANS}} = n_{\text{TOTAL}}$, where $n_{\text{TRANS}}$ and $n_{\text{TOTAL}}$ represent the number of transient frames and the total number of frames analyzed for the image set, respectively. For a given image set, $\lambda 2_{\text{ON}}$ was used to compute $\boldsymbol{T}(x^*, y^*, t)$ for image data captured within all ON and temporally transient segments of the overall magnetic actuation waveform sequence, whereas $\lambda 2_{\mathrm{OFF}}$ was used to compute stress traction solutions for image data captured with all OFF segments. Of note, $\lambda 2_{\mathrm{ON}}$ and $\lambda 2_{\mathrm{OFF}}$ both depend on the assumed values of $E$ and ν used to define the mechanical properties of the substrate.

#### *3. Movies*

All real-time annotated movies associated with FC mode experiments were created using the original DIC and epifluorescence fast-time lapse data sets captured at a rate of 40 fps. Given their large size, these videos were digitally compressed for uploading to the journal website using open-source video transcoder software HandBrake software v.1.3.3 (www.handbrake.fr). The original movie and data files are available from the authors upon reasonable request.

### B. Displacement-control (DC) mode

#### *1. Experimental procedure*

For DC experiments, the needle core of the MT device was inclined at ~40° with the horizontal (see **Fig. 1(b)** of the main article). Under live DIC imaging mode (~16 to 40 fps) at 20X magnification, the MT device was actuated to maintain a null magnetic force ($\boldsymbol{F}_{\mathrm{MT}}$ = 0 nN). The needle was then translated to a position near a superparamagnetic bead of interest such that $\boldsymbol{\delta}$ ~ 1 μm. During translation of the needle, we intentionally avoided contact between the needle tip and the surface of the collagen gel or keratinocyte sheet. The MT device was then actuated with a magnetic flux density setpoint of $B_{\mathrm{blunt}}^{\mathrm{ON}^*}$ = 175 Gs, which magnetically clamped the superparamagnetic bead to the needle tip. Displacements of the superparamagnetic bead in the $x$-, $y$- and $z$-directions that occurred during the magnetic clamping procedure were often negligible, and at most ~1 μm. When measurable displacements were induced by the magnetic clamping procedure, with the bead still clamped, the needle tip was translated to return the superparamagnetic bead back to the initial $(x, y)$-location that was observed for the bead prior to magnetic actuation. With the superparamagnetic bead in its original pre-clamped position, a fast time-lapse image acquisition at 40 fps (2048 x 2044 pixels per frame) was then initiated within Nikon Elements. After a ~1 s to 5 s pause, the needle was then *manually* translated away from this initial position in 1 μm-step increments, holding each position for ~1 s before proceeding with the next incremental step, until a maximum translation of 10 μm was reached. After a ~2 s hold at this maximum displacement of 10 μm, the needle was again manually translated back to its initial position in an identical stepwise fashion. After a brief 1 s to 5 s pause, the magnetic flux density of the MT device was set to achieve a null force

($F_{\mathrm{MT}} = 0$ nN), releasing the superparamagnetic bead from the needle tip. Image acquisition was then terminated.

Upon termination of the DIC imaging experiment, the needle was translated to a spatial location remote from the superparamagnetic bead of interest and demagnetized[1]. With the microscope still configured for live DIC imaging, the needle was then translated back to the bead of interest and the clamping procedure as previously described was conducted, including, if necessary, re-configuration of the superparamagnetic bead back to its original pre-clamped position. Imaging was then switched to epifluorescence mode and the focus of the microscope was adjusted to optimize viewing of the fluorescent microspheres embedded in the surface of the collagen substrate. Magnification remained constant at 20X. The position of the needle tip remained fixed. After a ~60 s-delay to account for adjustments in microscope settings, a second fast time-lapse image acquisition (2048x2044 pixels per frame) was initiated within Nikon Elements. The needle was stepwise translated to a maximum displacement of 10 μm and then back to the initial position exactly as was done during the preceding DIC imaging experiment. After a brief 1 s to 5 s pause, the magnetic flux density of the MT device was set to achieve a null force ($F_{\mathrm{MT}}$ = 0 nN) and image acquisition was terminated.

#### *2. Data reduction*

Data reduction for coupled DIC and epifluorescence image sets was carried out using the full field as the global ROI (2048x2044 pixels). Analyses used identical variables as previously defined for FC experiments (see Sec. SI.A.2). Temporal correlation between data derived from the DIC and epifluorescence image sets was achieved by first searching the epifluorescence data set to find the raw time point, $t_{\mathrm{EPI}}$, at which $\boldsymbol{\Delta}_{\mathrm{max}}(x^*_{\mathrm{max}}, y^*_{\mathrm{max}}, t_{\mathrm{r}})$ first becomes ≥0.15 μm, i.e., the raw time, $t_{\mathrm{r}}$, marking the first physically significant microsphere displacement measurement (see **Fig. S6**). We then temporally shifted the entire epifluorescence data set, re-defining the temporal variable for this data set, $t$, as $t = t_{\mathrm{r}} + (t_{\mathrm{DIC}} - t_{\mathrm{EPI}})$, where $t_{\mathrm{DIC}}$ was defined as the first time point at which $\boldsymbol{\Delta}_{\mathrm{MT}}(t) \geq 0.28$ μm, i.e., the time marking the first physically significant superparamagnetic bead displacement as derived from DIC imaging data (see **Fig. S5**). With this alignment of the epifluorescence and DIC data sets, the first experimentally significant microsphere and superparamagnetic bead displacements are temporally coincident. Once the two image sequences were temporally aligned, $\boldsymbol{u}(x^*, y^*, t)$ and $\boldsymbol{\Delta}_{\mathrm{max}}(x^*_{\mathrm{max}}, y^*_{\mathrm{max}}, t)$ were calculated as previously described. In solving for $\boldsymbol{T}(x^*, y^*, t)$, we again employed an algorithmic approach for selection of the L2 regularization parameter. As was done for force-control mode, the raw optimal regularization parameter at each instance of time, $\lambda 2(t)$, was determined using a Bayesian methodology[3]. We then binned each $\lambda 2(t)$ data point as ON or OFF, reassigning the independent variable,

$t$, to a generic frame number, $n$, or $\lambda 2(t) \rightarrow \lambda 2(n)$. ON frames were defined as images in which $t > t_{\text{DIC}}$. All other imaging frames were binned as OFF frames. We then computed two global L2 regularization parameters, $\lambda 2_{\text{ON}}$ and $\lambda 2_{\text{OFF}}$, based on the mean of the logarithms of $\lambda 2(n)$ for each ON or OFF binned data set, as per Eq. (S1). For a given image within the overall epifluorescence data set, $\lambda 2_{\text{OFF}}$ or $\lambda 2_{\text{OFF}}$ was used to compute $\boldsymbol{T}(x^*, y^*, t)$, $\boldsymbol{F}(t)$, $\rho(x^*, y^*, t)$, and $U(t)$ based on whether or not the frame was binned as an ON or OFF frame. $F_{\text{MT}}(t)$ was not evaluated for displacement-control mode of MT-DTM/TFM operation.

#### *3. Movies*

All real-time annotated movies associated with DC mode experiments were created using the original DIC and epifluorescence fast-time lapse data sets captured at a rate of 40 fps. Given their large size, these videos were digitally compressed for uploading to the journal website using open-source video transcoder software HandBrake v.1.3.3 (www.handbrake.fr). The original movie and data files are available from the authors upon reasonable request.

### SII. REFERENCES


1. W.I. Moghram, A. Kruger, E.A. Sander, and J.C. Selby, Rev. Sci. Instrum. Submitted (2020).
2. P. Kollmannsberger and B. Fabry, Rev. Sci. Instrum. **78**, 114301 (2007).
3. Y. Huang, C. Schell, T.B. Huber, A.N. Şimşek, N. Hersch, R. Merkel, G. Gompper, and B. Sabass, Sci. Rep. **9**, 539 (2019).
4. S.J. Han, Y. Oak, A. Groisman, and G. Danuser, Nat. Methods **12**, 653 (2015).

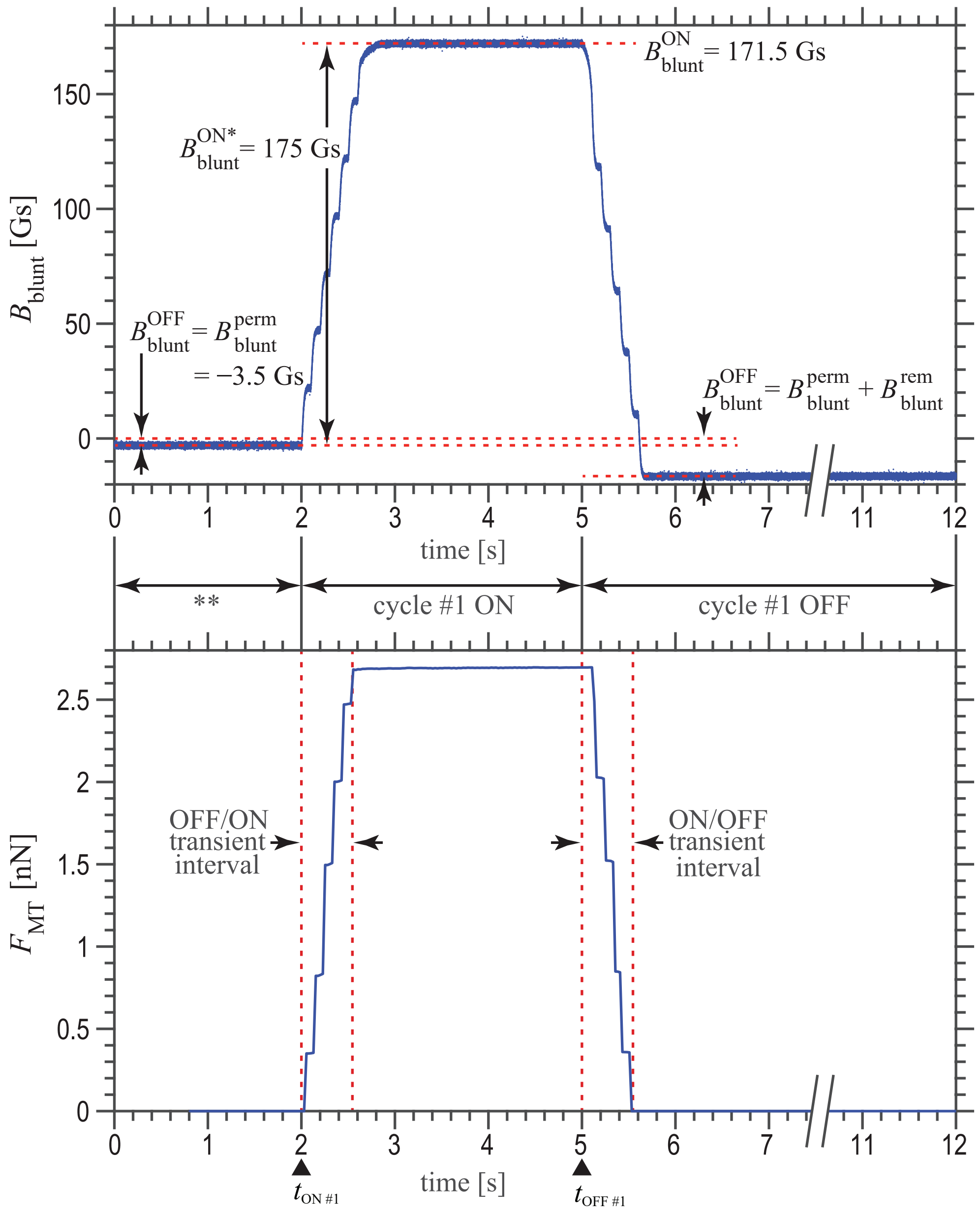


**FIG. S1.** Panel **(a)** shows the measured magnetic flux density measurements, $B_{\text{blunt}}(t)$, observed during cycle #1 of the 3 s/7 s ON/OFF FC mode bead-on-gel experiment detailed in **Fig. 2** of the main article. Panel **(b)** shows the computed magnitude of the magnetic tweezer force vector in the $xy$-plane of the collagen substrate, $F_{\text{MT}}(t)$ (see Sec. SI.A.2). The time interval denoted with ** indicates the 2 s-gap preceding actuation of the MT device during which fast time-lapse image acquisition begins at a random time point due to the non-deterministic operating system used to run Nikon Elements software.

**FIG. S2.** Six-panel figure highlights data from an FC mode null force control experiment involving a fibronectin-coated 4.5 μm diameter superparamagnetic bead attached to the surface of a ~680 μm thick, 1.0 mg/mL type I collagen substrate containing a surface layer of covalently attached red fluorescent microspheres. In this experiment, by design, $F_{\mathrm{MT}}(t)$ =0 nN for all time, $t$. **(a)** DIC image demonstrating that the superparamagnetic bead has no observable displacement near the end of the first ON cycle of the overall null force magnetic actuation waveform. **(b)** Texas red epifluorescence image with quivers showing displacement vectors with grossly zero magnitudes for all individual microspheres embedded within the surface of the collagen substrate at this same time point in the overall null force magnetic actuation waveform. The displacement field, $\boldsymbol{u}(x^*, y^*)$, and stress traction vector field, $\boldsymbol{T}(x^*, y^*)$, computed for the microsphere displacements displayed in **(b)** are shown in **(c)** and **(d)**, respectively. A plot of the magnitude of the maximum microsphere displacement, $\Delta_{\mathrm{max}}(t)$, versus the magnitude of the superparamagnetic bead displacement, $\Delta_{\mathrm{MT}}(t)$, is shown in **(e)**, suggesting a noise floor in bead displacements of ~0.1 μm. A plot of the null $F_{\mathrm{MT}}(t)$ versus the magnitude of the integrated total traction force vector, $F(t)$, is shown in **(f)**, where $F(t)$ has been computed assuming $\upsilon = 0.4$ and two different elastic moduli, $E = 14.0$ Pa and $E = 76.23$ Pa. The noise floor in $F(t)$ has a clear dependence on the assumed value of $E$.

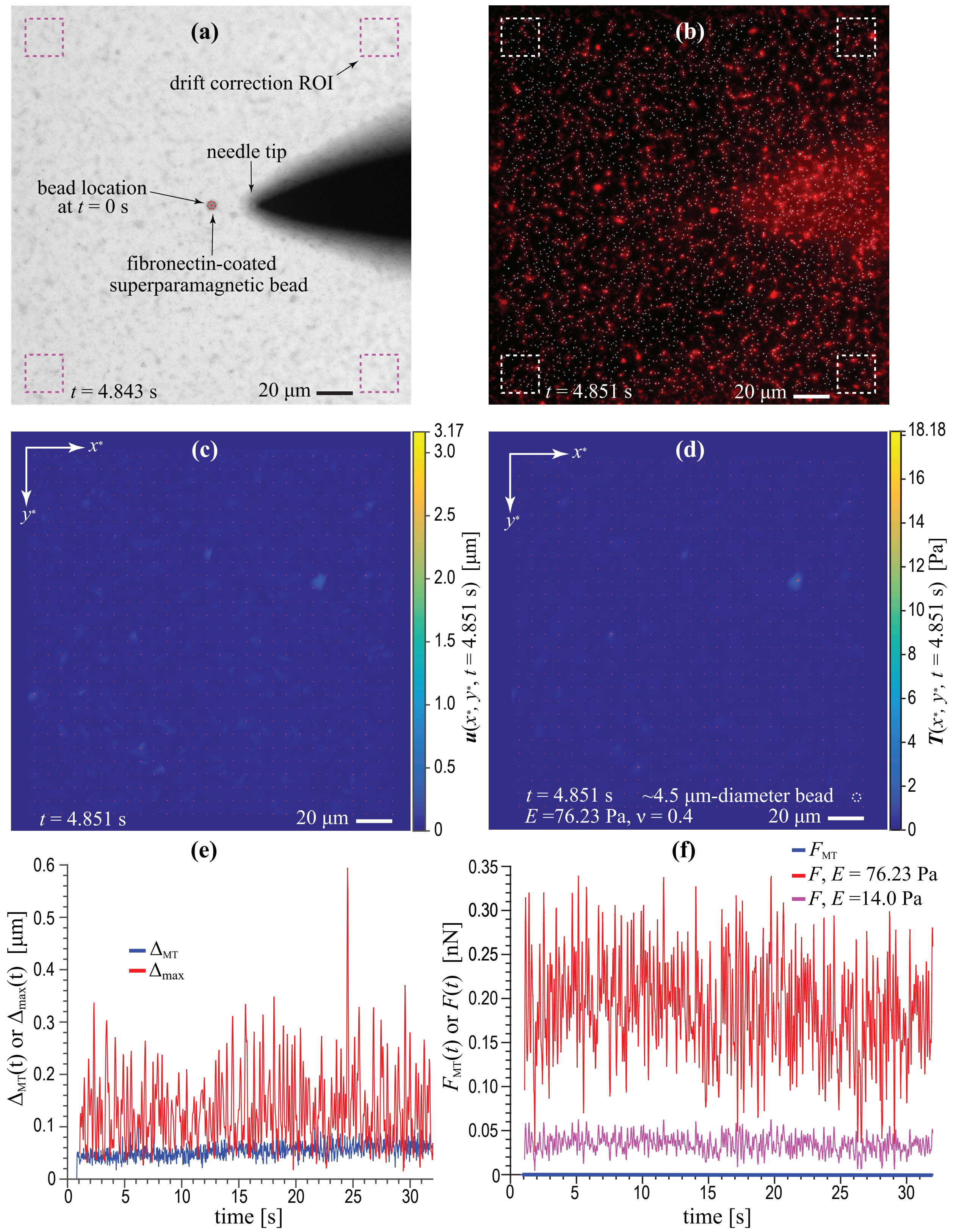


**FIG. S2.** FC mode null force control experiment.

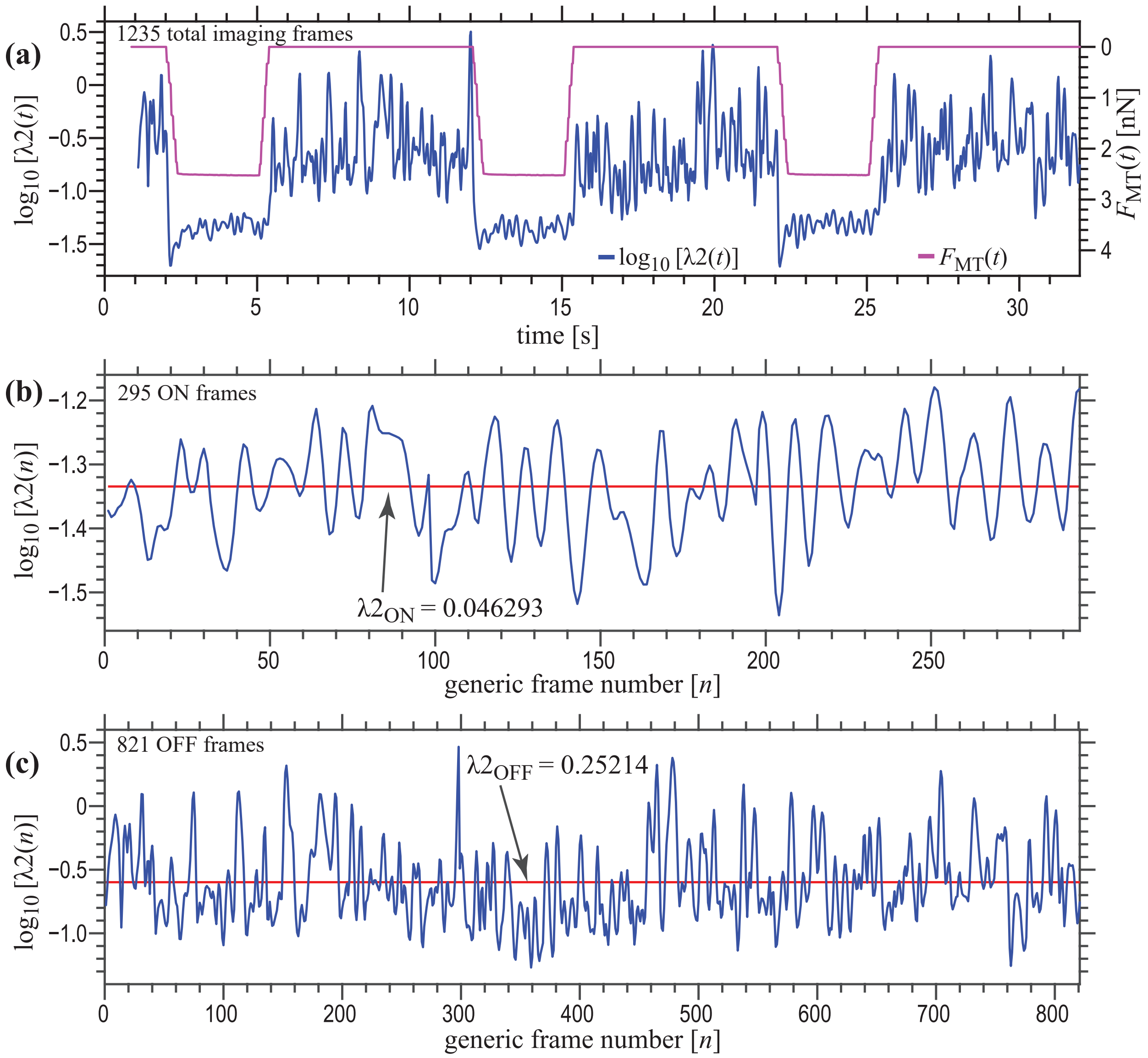


**FIG. S3. (a)** Frame-by-frame (raw) values of the L2 regularization parameter, λ2, found using a Bayesian approach applied to data from the prototype FC bead-on-gel experiment detailed in **Fig. 2** of the main article, assuming $E = 76.23$ Pa and $\upsilon = 0.4$. Fluctuations in the observed raw values of λ2 are temporally correlated with the ON/OFF fluctuations of $F_{\mathrm{MT}}(t)$. After omitting transient imaging frames and binning individual imaging frames as ON **(b)** or OFF **(c)**, we defined two global L2 regularization parameters, $\lambda 2_{\mathrm{ON}}$ =0.04629 and $\lambda 2_{\mathrm{OFF}}$ =0.25214, respectively, based on the mean of the logarithms of the binned data sets (see Sec. SI.A.2).

**FIG. S4.** Plots showing time-resolved vectoral components in the $x^*$- and $y^*$-directions of the displacements, $\boldsymbol{u}(\text{µ-sphere } [i], t)$, and accelerations, $\frac{\partial^2}{\partial t^2}\boldsymbol{u}(\text{µ-sphere } [i], t)$, of microspheres, $\text{µ-sphere } [i]$, embedded within the collagen substrate and subject to the FC mode null force control experiment described in Sec. IV.A.1 of the main article. Given that 3266 microspheres were tracked over 1234 imaging frames, displacement and acceleration plots show only decimated data points representative of the total acquired data set, specifically, 4% of all displacements and 4% of all accelerations (i.e., every 25$^{\text{th}}$ $\text{µ-sphere } [i]$) are plotted for every other imaging frame, $t$. Horizontal dashed lines in each plot represent the statistical upper and lower bounds of the data defined by +/-6 standard deviations (6SD), bounds that we assume to approximate the uncertainty in our measurements for microsphere displacements and accelerations.

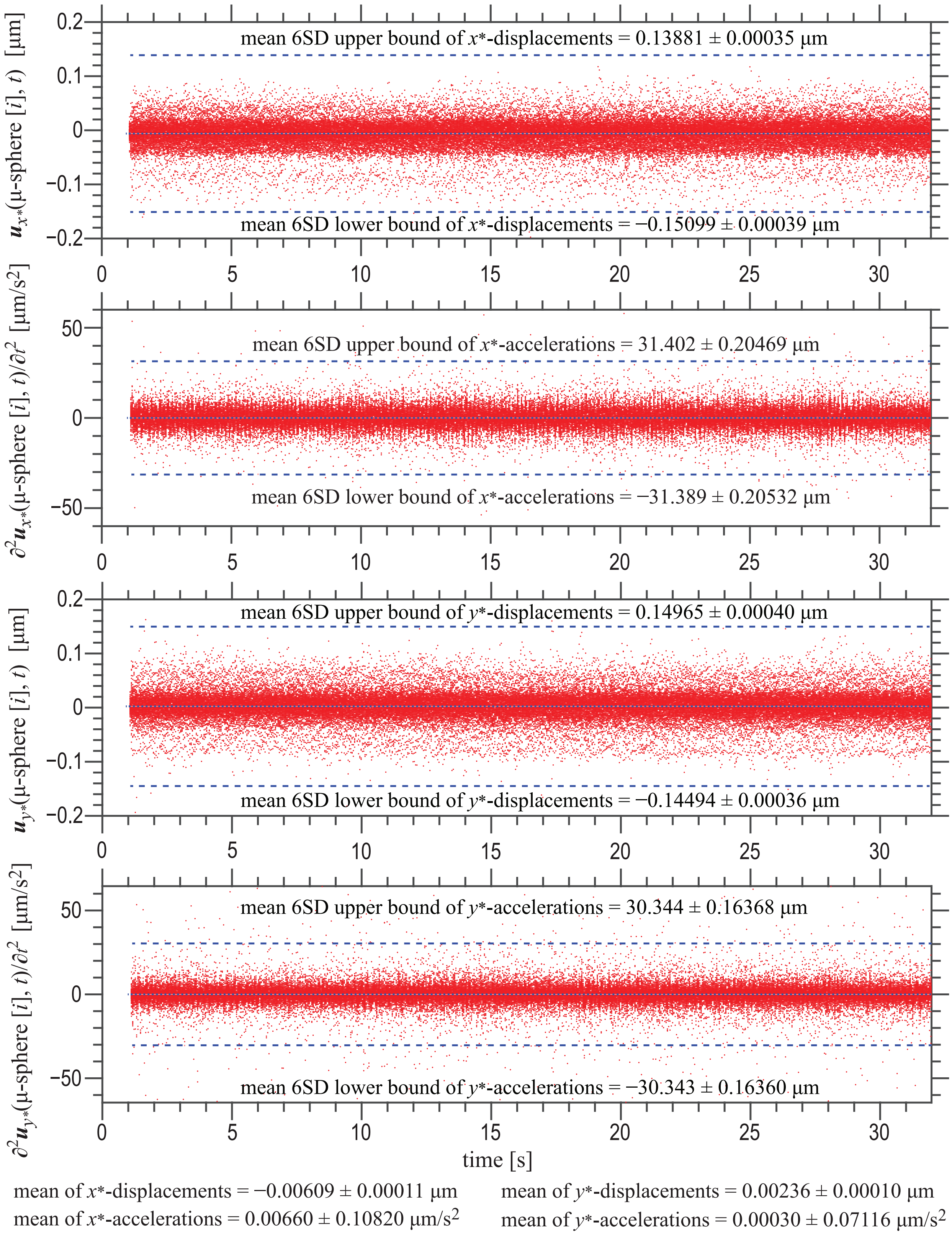


**FIG. S4.** Microsphere displacements and accelerations for the FC mode null force control experiment.

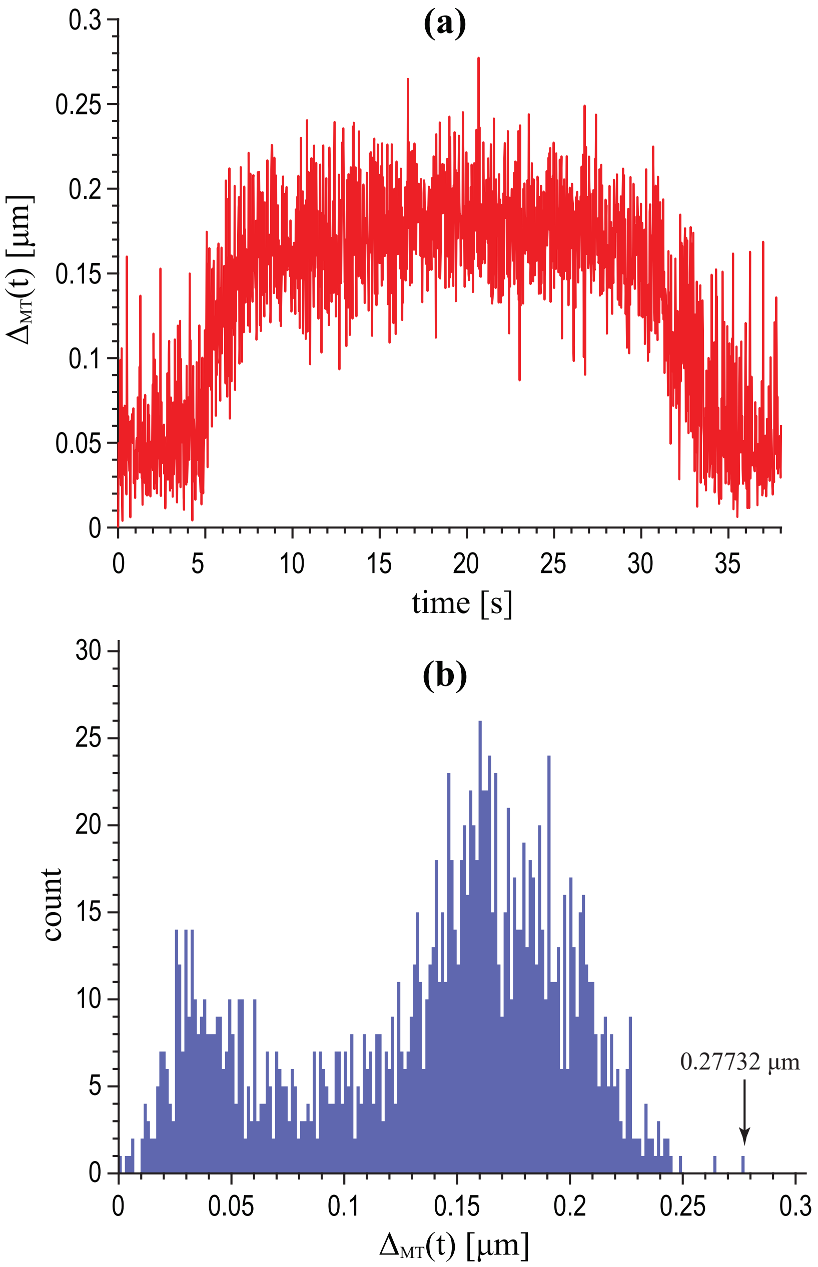


**FIG. S5.** Magnitude of the superparamagnetic bead displacement, $\Delta_{\mathrm{MT}}(t)$, as measured during a DC mode null displacement control experiment (see Sec. IV.A.2 of the main article) as a function of time **(a)** or alternatively, plotted as a histogram **(b)**. In this experiment, we knew *a priori* that the superparamagnetic bead was physically *static* throughout all 1520 acquired DIC imaging frames. As such, the data represent a measure of displacement uncertainty intrinsic to our bead tracking algorithms. Knowing that needle translation occurred between $t$ = ~5 s and $t$ = ~30 s, we note that our tracking uncertainty exhibits a dependence on the relative motion of the needle tip that can be observed within each DIC image. Given that the largest observed displacement for a *static* superparamagnetic bead was $\Delta_{\mathrm{MT}}$ = 0.27732 μm, for purposes of defining $t_{\mathrm{DIC}}$, we assume that the superparamagnetic bead first experiences a significant displacement away from its initial position when $\Delta_{\mathrm{MT}}(t_{\mathrm{DIC}}) > 0.28$ μm.

**FIG. S6.** In a DC mode null displacement control experiment (see Sec. IV.A.2 of the main article), the displacements of 6088 fluorescent microspheres were tracked over 1425 imaging frames. The above plot contains a histogram showing the magnitude of microsphere displacements, $|\boldsymbol{u}(\mu\text{-sphere }[i],\ 16.492\text{ s})|$, tracked for all microspheres, $i$, in frame 660 ($t$ = 16.492 s). Assuming that at each time point, $t$, the $x^*$- and $y^*$-components of $\boldsymbol{u}(\mu\text{-sphere }[i], t)$ are uncorrelated, normally distributed, and of equal variance with zero mean, $|\boldsymbol{u}(\mu\text{-sphere }[i], t)|$ for all $i$ will approximate a Rayleigh distribution. For each frame, then, the tracked microsphere displacements were used to find a best-fit Rayleigh distribution parameter, $\sigma$. Here σ = 0.02084 μm for frame 660. The mean value of all best-fit Rayleigh distribution parameters was then defined over all 1425 imaging frames as $\overline{\sigma}$, where for this negative displacement control experiment, $\overline{\sigma}$ = 0.02339 μm. Assuming a Rayleigh distribution with $\overline{\sigma}$ =0.02339 μm, we define $|\boldsymbol{u}_{6\overline{\sigma}}(\mu\text{-sphere }[i])|$ as the displacement magnitude that approximates the $6\overline{\sigma}$ upper limit of the microsphere displacement distribution, or in other words, the displacement that will be greater than 99.9999998026825% of any singly observed microsphere displacement measurement. Here, we find $|\boldsymbol{u}_{6\overline{\sigma}}(\mu\text{-sphere }[i])|$ = 0.14585 μm. Knowing *a priori* that the microspheres were static throughout the null displacement control experiment, ~0.15 μm represents a measure of microsphere displacement uncertainty intrinsic to our microsphere tracking algorithms. For purposes of defining $t_{\text{EPI}}$ and correlating DIC imaging data with epifluorescence imaging data in a DC mode experiment, we assume that the initial collagen substrate motion first occurs when $\Delta_{\max}(t_{\text{EPI}})$ > 0.15 μm (time with respect to raw imaging time, $t_{\text{r}}$). In comparing displacement uncertainties as derived from our FC experiments done at 30X magnification (see **Fig. S4**) and our DC experiments done at 20X magnification, we see that the uncertainty in microsphere displacements are similar (~0.15 μm), though one would expect greater uncertainty for experiments done at 20X magnification. To reconcile this finding, we note that different thresholds of spatial outlier detection were used in analyzing the imaging data obtained in our FC versus our DC experiments. Specifically, more strict outlier detection limits were employed for the DC controlled experiments, decreasing the overall displacement noise and permitting smaller measurement uncertainties than expected.

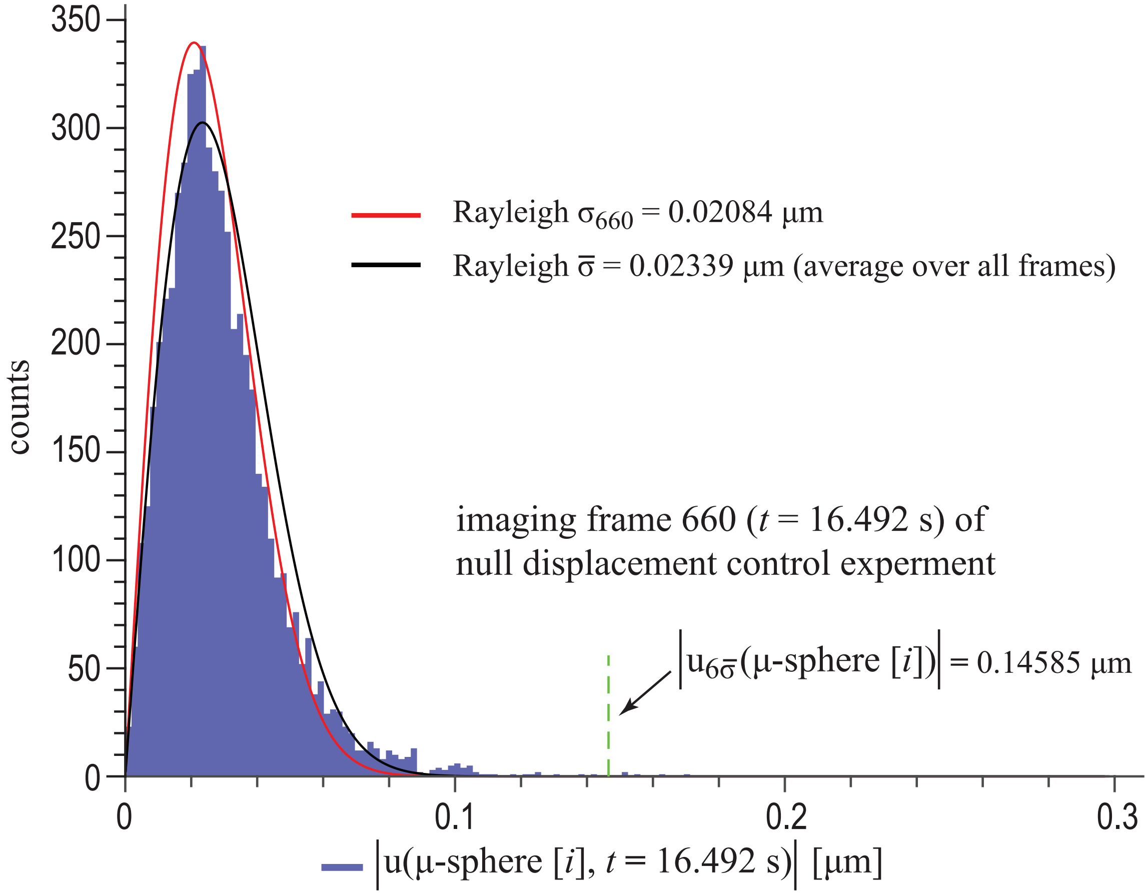


**FIG. S6.** Microsphere dynamics in a DC mode null displacement control experiment.

**FIG. S7.** Six-panel figure highlights data from a DC mode null displacement control experiment involving a fibronectin-coated 4.5 μm diameter superparamagnetic bead attached to the surface of a ~700 μm thick, 1.0 mg/mL type I collagen substrate embedded with a surface layer of covalently attached red fluorescent microspheres. In this experiment, by design, $F_{\mathrm{MT}}(t)$ = 0 nN for all time, $t$. Temporal correlation between the DIC and epifluorescence image sets could not be done as discussed in **Fig. S5** and **Fig. S6** given the absence of true superparamagnetic bead and microsphere displacements. **(a)** DIC image demonstrating that the superparamagnetic bead has no observable displacement despite stepwise translation of the needle tip ~10 μm from its initial position. **(b)** Texas Red epifluorescence image showing quivers representative of grossly zero displacement vectors for all individual microspheres embedded within the collagen substrate after the needle tip has been translated ~10 μm from its initial position. The displacement field, $\boldsymbol{u}(x^*, y^*)$, and stress traction vector field, $\boldsymbol{T}(x^*, y^*)$, computed for the microsphere displacements displayed in **(b)** are shown in **(c)** and **(d)**, respectively. A plot of the maximum substrate microsphere displacement, $\Delta_{\mathrm{max}}(t)$, versus superparamagnetic bead displacement, $\Delta_{\mathrm{MT}}(t)$, is shown in **(e)**. Knowing that stepwise needle translation to and from its maximum displacement occurred between $t$ = ~5 s and $t$ = ~30 s, one can observe that the noise in tracking of $\Delta_{\mathrm{MT}}(t)$ is correlated with motion of the needle tip in the acquired DIC images, despite the fact that there was no physical connection between the tip and the superparamagnetic bead during the experiment. The mean noise floor in microsphere displacements is ~0.075 μm. A plot of the integrated total traction force vector, $F(t)$, is shown in **(f)**, where $F(t)$ has been computed assuming υ = 0.4 and $E$ = 52.73 Pa. The mean noise floor in $F(t)$ for these assumed material properties is ~0.7 nN.

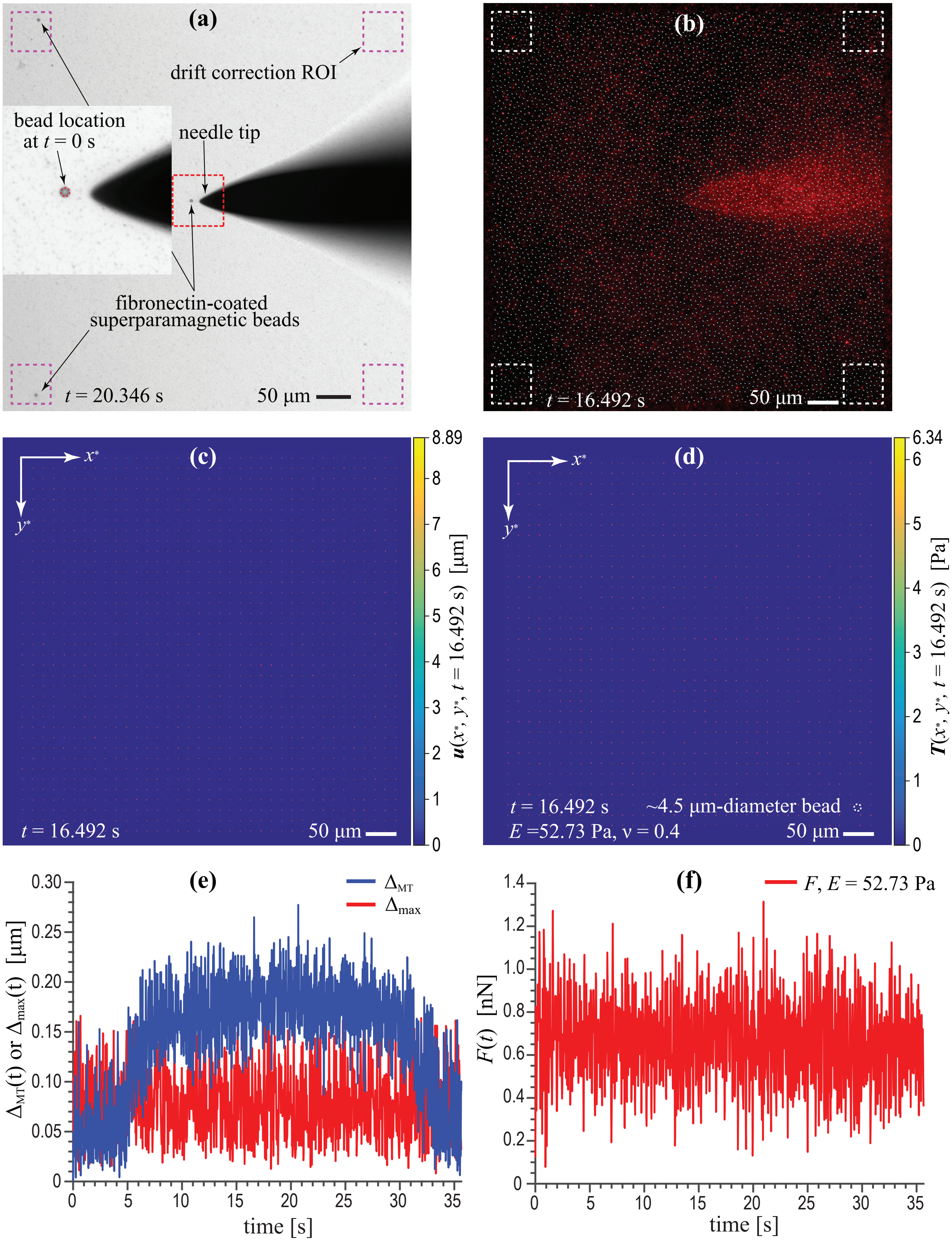


**FIG. 7.** DC mode null displacement control experiment.

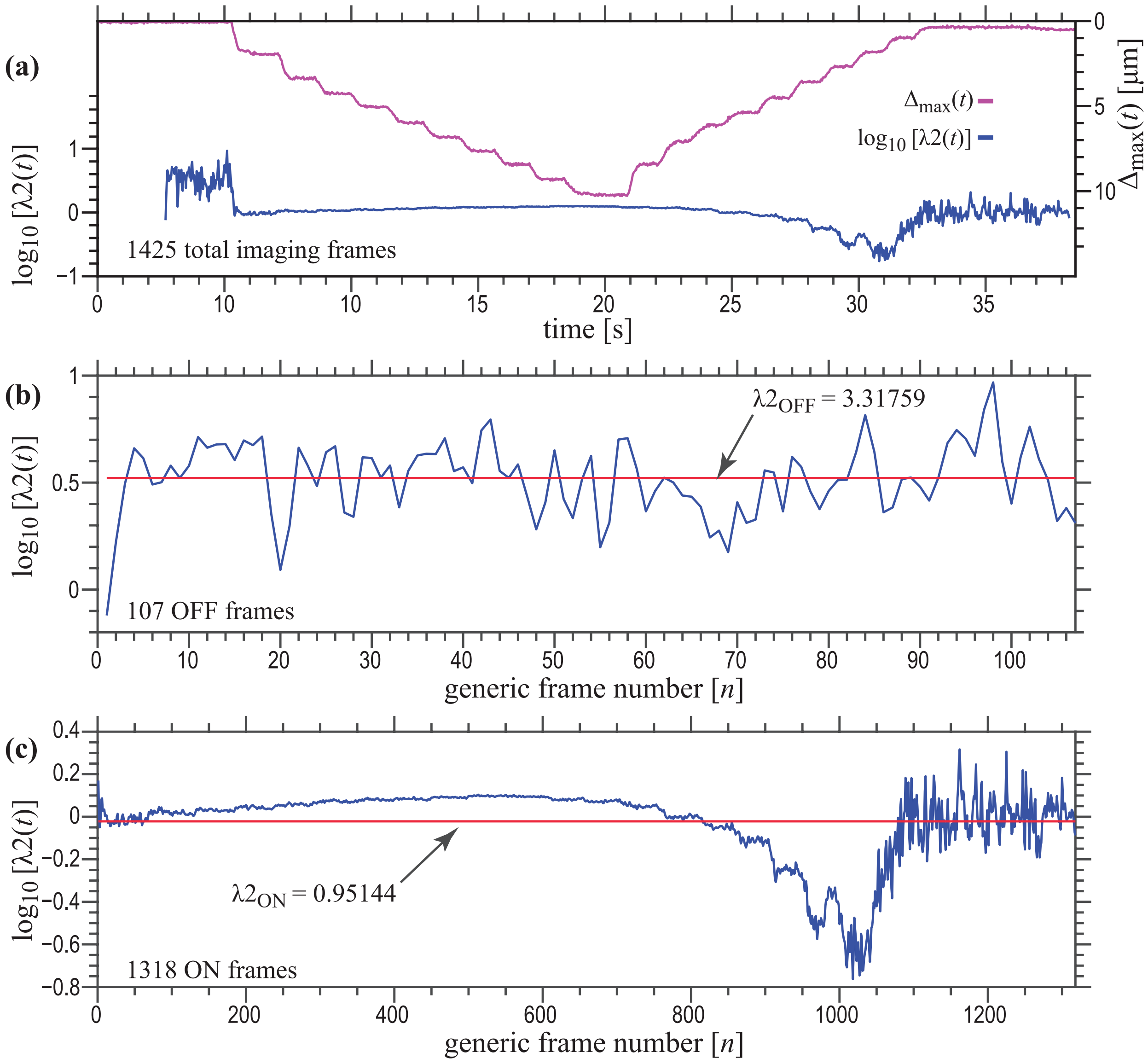


**FIG. S8. (a)** Frame-by-frame (raw) values of the L2 regularization parameter, λ2, found using a Bayesian approach applied to imaging data from the prototype DC mode bead-on-gel experiment detailed in **Fig. 5** of the main article, assuming $E = 52.73$ Pa and $\upsilon = 0.4$. $\Delta_{max}(t)$ is plotted in **(a)** to demonstrate the correlation between substrate displacements and the frame-by-frame Bayesian determination of λ2. After binning individual imaging frames as OFF **(b)** or ON **(c)**, we defined two global L2 regularization parameters, $\lambda 2_{ON} = 0.95144$ and $\lambda 2_{OFF} = 3.31759$, respectively, based on the mean of the logarithms of the binned data sets (see Sec. SI.B.2).

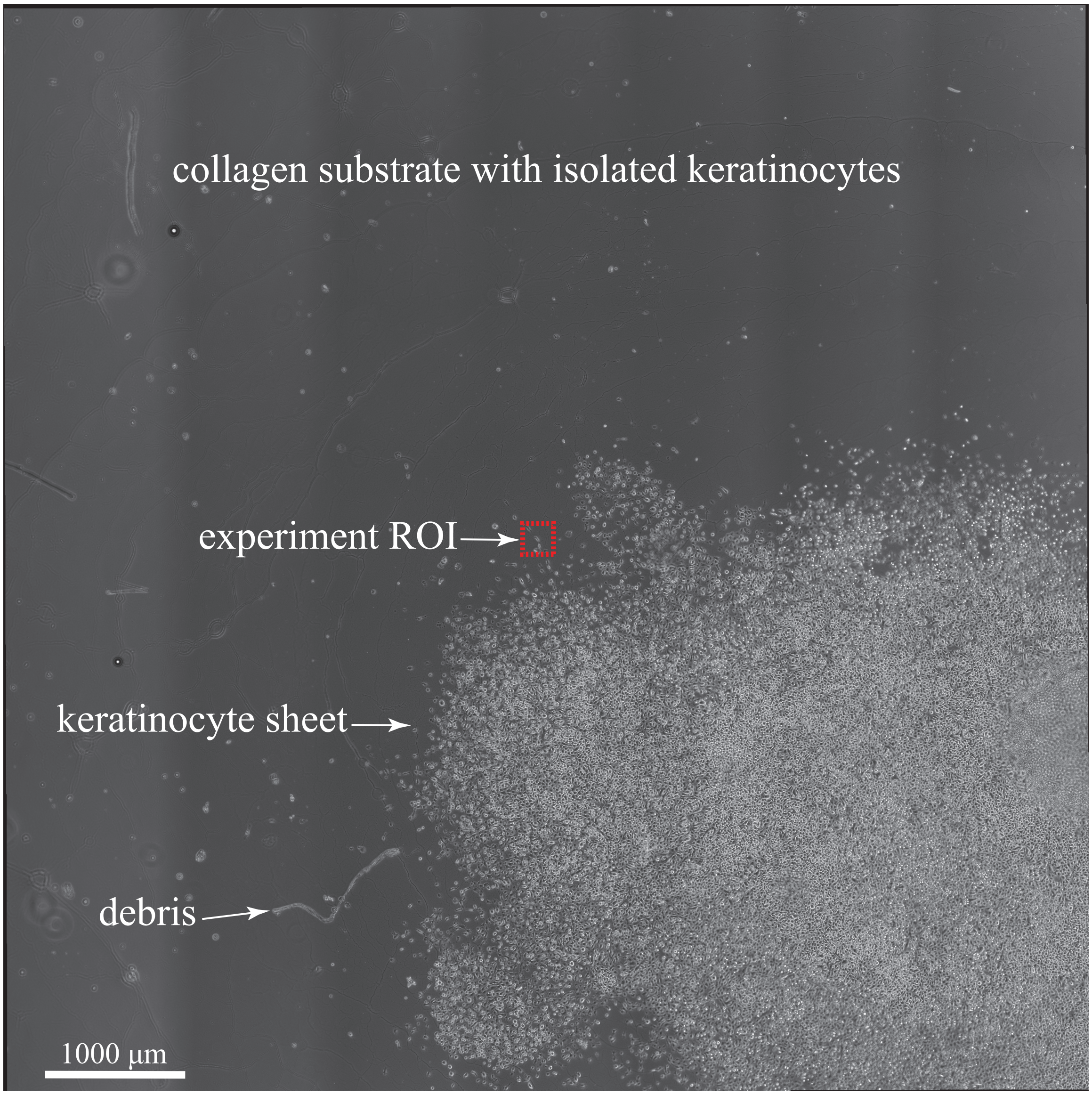


**FIG. S9.** Tiled and stitched phase contrast image showing the reconstituted multicellular keratinocyte sheet used for the FC mode bead-on-cell keratinocyte experiment detailed in Sec. IV.B of the main article. The keratinocyte sheet occupies the lower right-hand corner of the image, while isolated keratinocytes are adherent to the underlying 530 μm-thick 2.0 mg/mL type I collagen substrate reside in the upper third of the image. The red dashed box delineates the ROI containing keratinocyte #1 that was interrogated for the experiment detailed in **Fig. 7** and **Fig. 8** of the main article (also see **Vid 23**, **Vid 24**, **Vid 25**, **Vid 26**, and **Vid 27**).

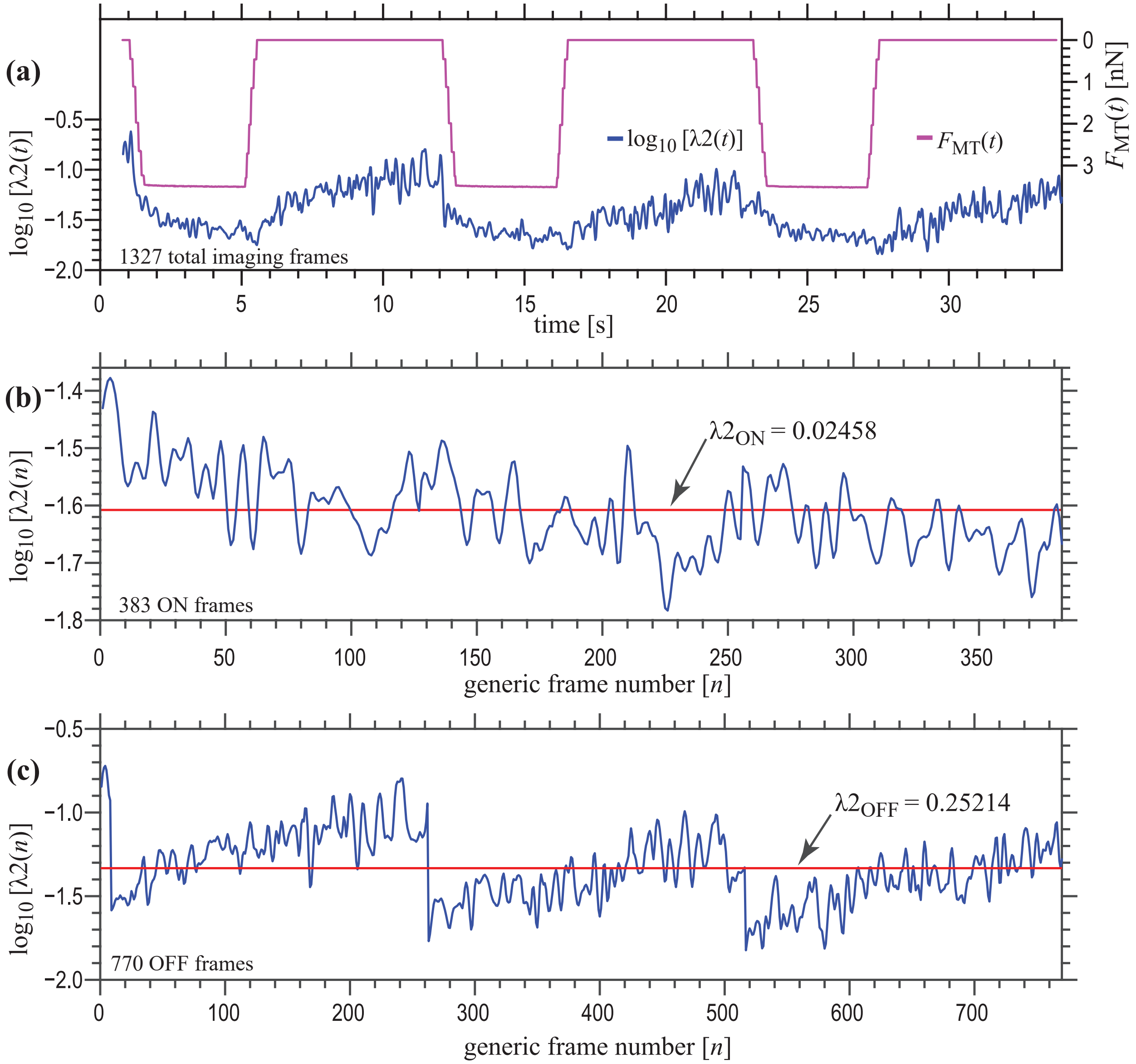


**FIG. S10. (a)** Frame-by-frame (raw) values of the L2 regularization parameter, λ2, found using a Bayesian approach applied to data from the FC mode bead-on-cell keratinocyte experiment detailed in **Fig. 7** of the main article, assuming $E = 138.3$ Pa and $\upsilon = 0.4$. Fluctuations in the observed raw values of λ2 are temporally correlated with the ON/OFF fluctuations of $F_{MT}(t)$. After omitting transient imaging frames and binning individual imaging frames as ON **(b)** or OFF **(c)**, we defined two global L2 regularization parameters, $\lambda 2_{ON}$ =0.02458 and $\lambda 2_{OFF}$ =0.04634, respectively, based on the mean of the logarithms of the binned data sets (see Sec. SI.A.2).

**FIG. S11.** Magnified view of **(a)** the displacement field, $\boldsymbol{u}(x^*, y^*, t = 27.139\ \text{s})$ and **(b, c)** the stress traction vector field, $\boldsymbol{T}(x^*, y^*, t = 27.139\ \text{s})$, of the collagen substrate subjacent to keratinocyte #1 computed near $t_{\text{OFF}}$ of cycle #3 of the bead-on-cell experiment described in Sec. IV.B of the main article. In this TFM solution for $\boldsymbol{T}$, we employed the boundary element method using 1024 mesh points and L1 regularization that used the "optimal" selection of the L1 regularization parameter, as described by Han et al.[4] For ease of visualizing the directionality of the stress traction vector field, traction quivers *external* to the keratinocyte have been removed in **(b)**, whereas quivers *within* the area of subjacent substrate have been omitted in **(c)**. Scale bars = 10 µm. As was observed for the FTTC-based solution (see **Figs. 8(a)** and **8(b)** of the main article), apparent "ringing" arises in the stress traction field due to the presence of large gradients within the displacement field.

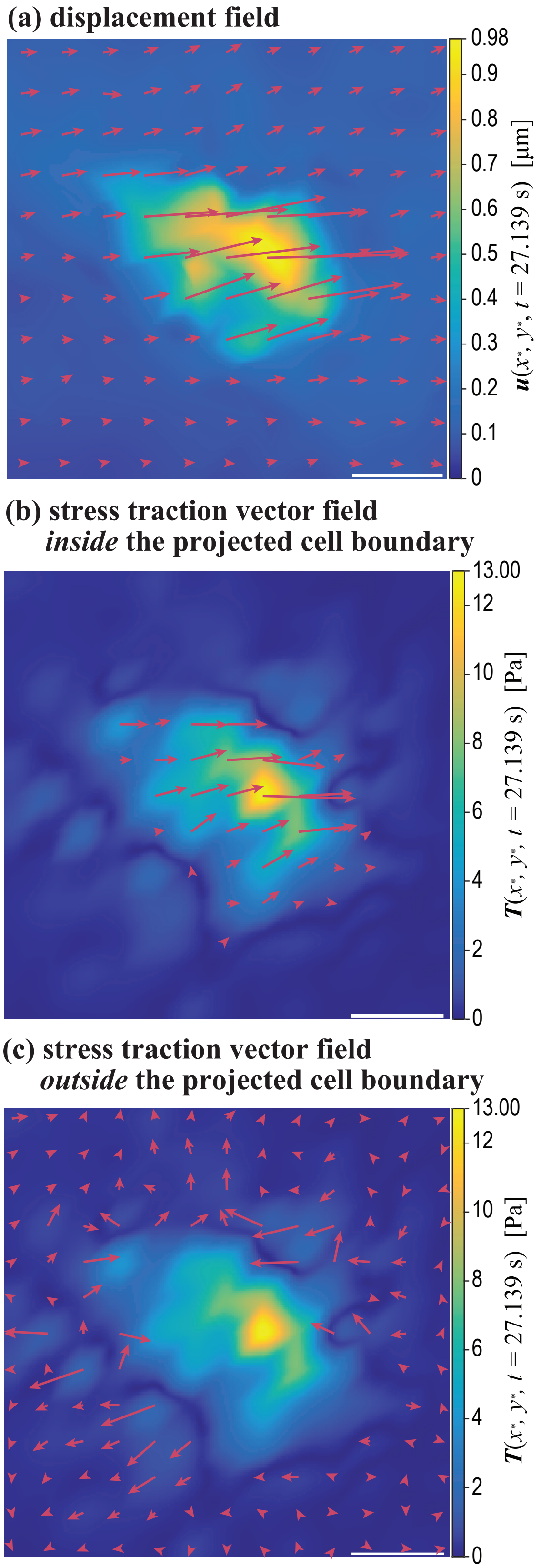


**FIG. S11.** $\boldsymbol{u}(x^*, y^*, 27.139\ \mathrm{s})$ and $\boldsymbol{T}(x^*, y^*, 27.139\ \mathrm{s})$ underlying keratinocyte #1.
